# Endothelial SOCS3 maintains homeostasis and promotes survival in endotoxemic mice

**DOI:** 10.1101/2020.12.28.424586

**Authors:** Nina Martino, Ramon Bossardi Ramos, Shuhan Lu, Kara Leyden, Lindsay Tomaszek, Sudeshna Sadhu, Gabrielle Fredman, Ariel Jaitovich, Peter A. Vincent, Alejandro P. Adam

## Abstract

SOCS3 is the main inhibitor of the JAK/STAT3 pathway. This pathway is activated by interleukin 6 (IL-6), a major mediator of the cytokine storm during shock. To determine its role in the vascular response to shock, we challenged mice lacking SOCS3 in the adult endothelium (SOCS3^iEKo^) with a non-lethal dose of lipopolysaccharide (LPS). SOCS3^iEKo^ mice died 16-24 hours post-injection after severe kidney failure. Loss of SOCS3 led to an LPS-induced type I interferon-like program, and high expression of pro-thrombotic and pro-adhesive genes. Consistently, we observed intraluminal leukocyte adhesion and NETosis, as well as retinal venular leukoembolization. Notably, heterozygous mice displayed an intermediate phenotype, suggesting a gene dose effect. In vitro studies were performed to study the role of SOCS3 protein levels in the regulation of the inflammatory response. In HUVEC, pulse-chase experiments showed that SOCS3 protein has a half-life below 20 minutes. Inhibition of SOCS3 ubiquitination and proteasomal degradation leads to protein accumulation and a stronger inhibition of IL-6 signaling and barrier function loss. Together, our data demonstrates that the regulation of SOCS3 protein levels is critical to inhibit IL-6-mediated endotheliopathy during shock and provides a promising new therapeutic avenue to prevent MODS though stabilization of endothelial SOCS3.

**Graphical abstract:** 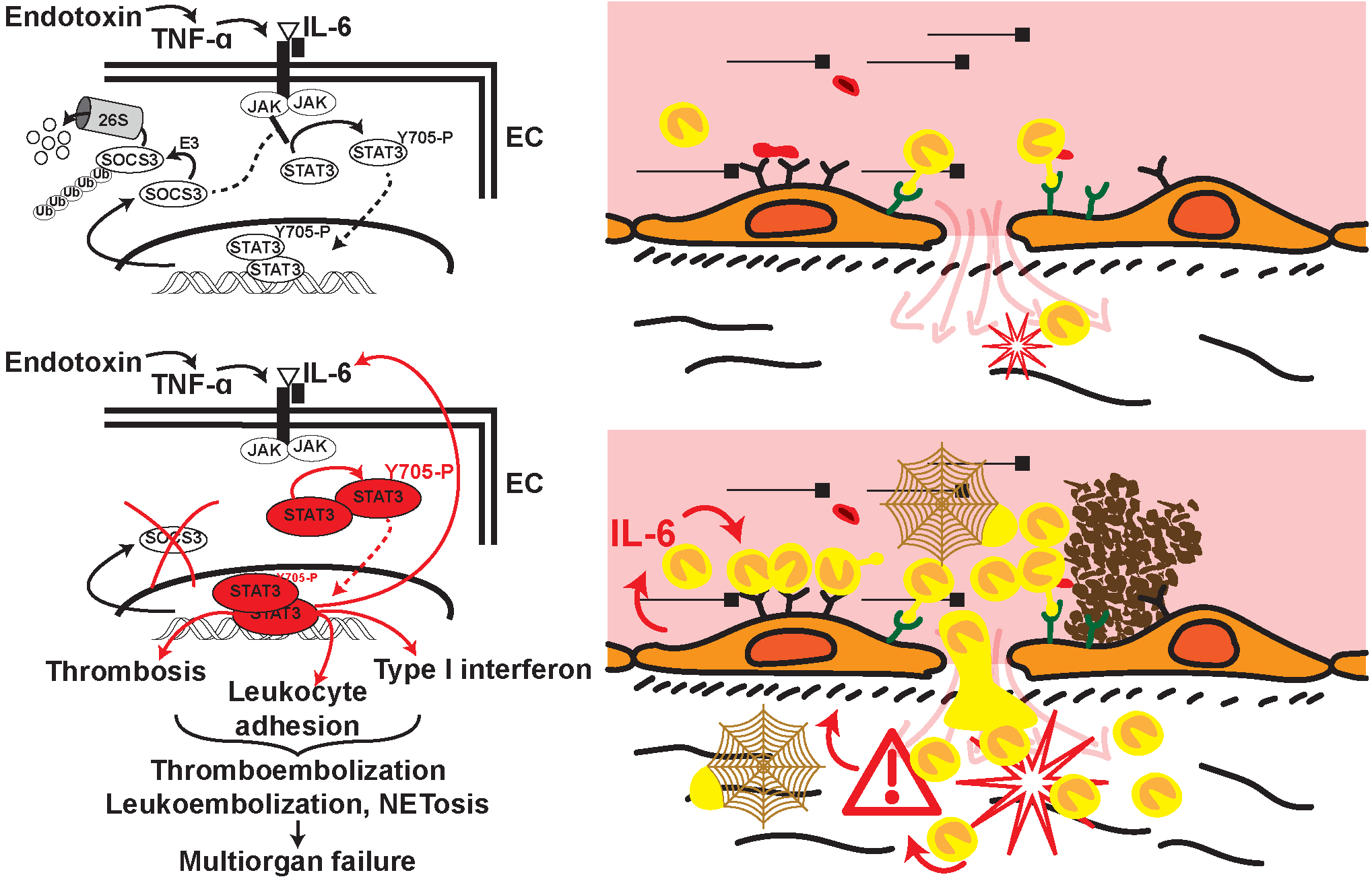

## Introduction

Systemic inflammatory response syndrome (1) often leads to multiorgan dysfunction (MODS) through an acute release of cytokines, in a process often called ‘cytokine storm’ (2). The release of these cytokines mediates the vascular dysfunction leading to shock, including refractory vasodilation, microvascular thrombosis, leukocyte plugging and increased vascular permeability. These defects lead to systemic hypotension, organ hypoperfusion and ultimately organ failure and death (3–6). These phenomena are mainly driven by the host’s immune response through a mechanism that involves, at least in part, cytokine-induced vascular dysfunction (7, 8). The circulating levels of the proinflammatory cytokine interleukin 6 (IL-6) correlate with disease severity (9–13) and are highly predictive of mortality (14, 15). In fact, an IL-6 amplifier mechanism that consists in a positive feedback regulation of this cytokine has been linked to worse prognosis (10). IL-6 binds to a heterodimeric receptor consisting of a transmembrane gp130 subunit that is responsible for signal transduction, and a smaller IL-6Rα, or gp80 subunit, that presents the cytokine to gp130 (10, 16). Very few cells express gp80, but a soluble form (sIL-6Rα) quickly appears in circulation during inflammation and presents the ligand to gp130 to activate its downstream signaling, in a process termed ‘trans-signaling’ (10, 16). Upon ligation, this receptor activates the Janus tyrosine kinases (JAK) to induce several downstream signal transduction cascades, including the activation of signal transduction and activator of transcription 3 (STAT3)-dependent gene expression through its phosphorylation on tyrosine 705 (10, 17). Multiple positive and negative feedback loops regulate the intensity and duration of this signal. Chiefly, STAT3-mediated expression of suppressor of cytokine signaling 3 (SOCS3) leads to a potent negative regulation. SOCS3 binds to the gp130/JAK complex to block further STAT3 phosphorylation (18–21).

The direct effects of IL-6 on the endothelium are not completely understood. Moreover, the role for its downstream signaling in promoting shock induced endotheliopathy is still unclear. Activation of this pathway leads to an increase in expression of leukocyte binding molecules (such as ICAM, E-Selectin, P-Selectin) at the surface of the endothelium, and a decrease in anti-thrombotic proteins (22). Many reports point to a role for IL-6 in promoting vascular leak in vivo (16, 23–26). Using in vitro models, we (27) and others (28–31) have shown that IL-6 directly promotes increases in endothelial permeability. In HUVEC, IL-6-induced increase in endothelial permeability is sustained for prolonged periods, which conflicts with the notion of a strong negative regulation via STAT3-induced SOCS3 expression. This barrier function loss requires STAT3 phosphorylation at Y705 and de novo protein synthesis (27). The transcriptional response in the endothelium leading to vascular leak, however, is not known.

Here, we sought to determine the role of SOCS3 in the regulation of the intensity and duration of IL-6 signaling in the endothelium by assessing the response to endotoxin in mice lacking SOCS3 specifically in the adult endothelial cells (SOCS3^iEKO^). We found that SOCS3 deletion leads to fast mortality less than 24 h after an endotoxin injection that is non-lethal in control mice. Surviving SOCS3^iEKO^ mice 16 hours post-challenge showed severe kidney failure, accumulation of intraluminal leukocytes in multiple organs, vascular leakage, and a pro-thrombotic and an adhesive transcriptional response that was associated with a strong induction of type I interferon-regulated genes. Notably, heterozygous mice displayed an intermediate phenotype, suggesting a gene dose effect. In vitro, we demonstrate that the sustained barrier function loss induced by an IL-6 treatment can be explained by fast SOCS3 ubiquitination and proteasomal degradation that prevents sufficient protein accumulation to act as an effective barrier to this pathway. Together, these findings demonstrate a crucial role for the regulation of endothelial SOCS3 levels to minimize vascular dysfunction and uncovers a promising and novel potential therapeutic target for balancing an uncontrolled inflammatory response.

## Results

### SOCS3^iEKO^ mice die within 24 hours after endotoxin challenge

To determine the role of endothelial SOCS3 in the vascular response to inflammation, we generated tamoxifen-inducible, endothelial-specific SOCS3 knockout mice (herein, SOCS3^iEKO^). For that purpose, we crossed SOCS3^fl/fl^ with mice carrying a cdh5-CreER^T2^ inducible endothelial Cre driver. A tdTomato reporter was also introduced. Control mice carried the Cre driver and tdTomato reporter, but wild type SOCS3 alleles. All mice were treated with tamoxifen for five consecutive days. Upon tamoxifen treatment, SOCS3^iEKO^ displayed a deletion of part of exon 2 (Figure 1A). Flow cytometry of cells obtained from the lung, a highly vascularized tissue, show that approximately 50% of cells were positive for tdTomato, while the fraction of cells positive in blood or in bone marrow was much lower (Figure 1B). Moreover, RT-qPCR of CD45 cells obtained from spleen (Supplemental Figure 1) demonstrates a lack of SOCS3 excision in these cells (Figure 1C), suggesting a low activity of this Cre driver in hematopoietic cells, a finding that is consistent with other reports (32). Mice were viable for several weeks post-excision, without any obvious phenotype. Then, we challenged SOCS3^iEKO^, control and heterozygous littermate controls two weeks post-excision with a single dose of endotoxin (250 μg/mouse of lipopolysaccharides from *E. coli* 0111:B4, LPS) or saline solution. As shown in Figure 1D, a dose of LPS that is not lethal in wild type mice, led to a very fast mortality in the SOCS3 knockouts. All LPS-treated SOCS3^iEKO^ mice died by 24 hours, with death starting at ~16 hours post treatment. To better understand the mortality in these mice, we developed a set of objective criteria based on recently described scoring systems for similar conditions, with small modifications (33–35) (described in Supplemental Table 5). As expected, endotoxemic mice showed an increased severity score, which was further aggravated by the SOCS3 deficiency (Figure 1E). Notably, even though all heterozygous littermates survived, they received an intermediate severity score, suggesting that loss of a single SOCS3 allele is sufficient to worsen the inflammatory response (Figure 1E). Further, LPS-induced hypothermia was much more pronounced in SOCS3^iEKO^ mice, again with heterozygous mice having intermediate values (Figure 1F).

**Figure 1.**
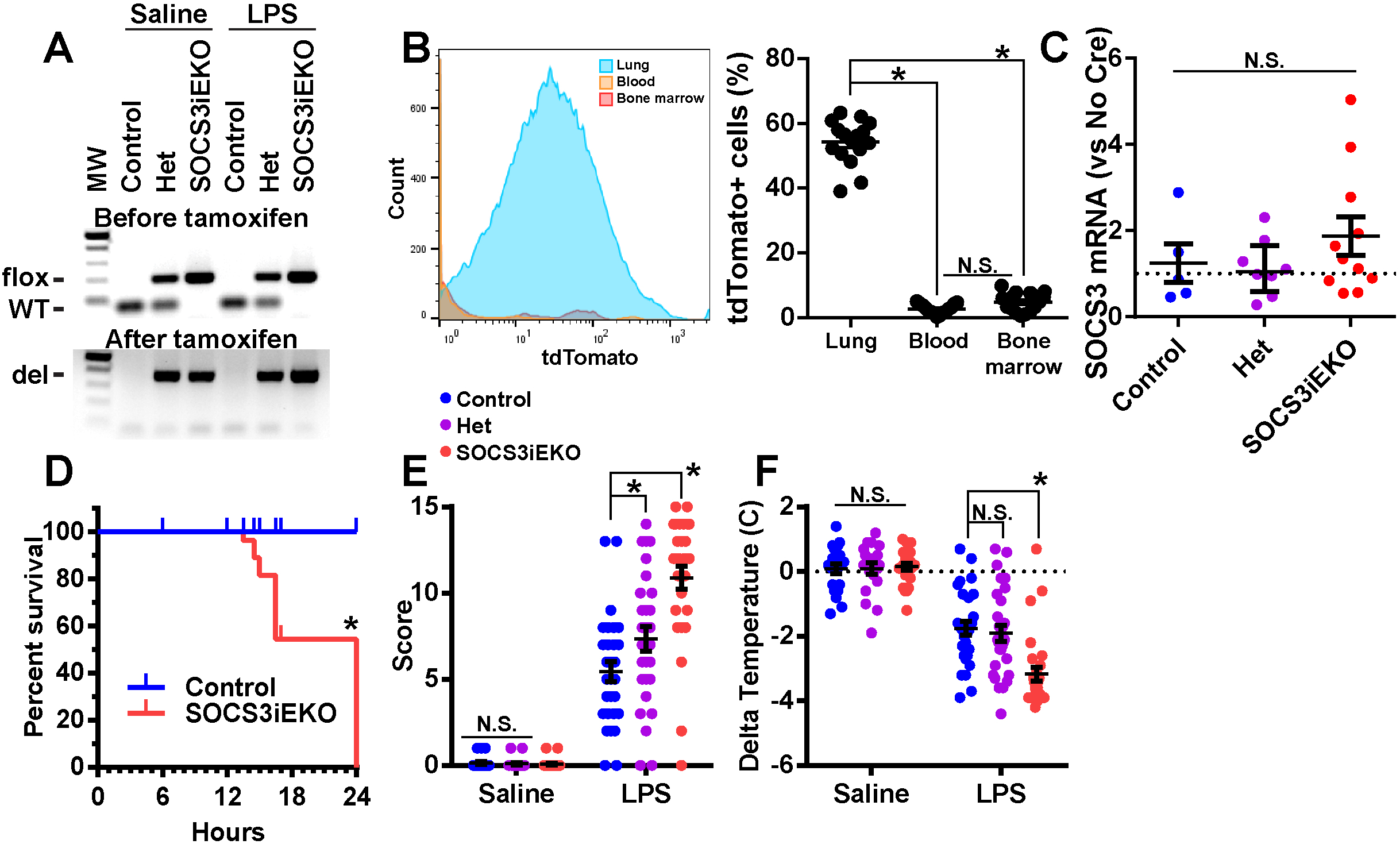
Loss of endothelial SOCS3 dramatically increase the severity of the response to endotoxin. A, Sample genotyping before and after tamoxifen addition confirming the deletion of SOCS3. B, Flow cytometry analysis of tomato expression showing only a minor expression in circulating and bone marrow cells and in a large proportion of lung cells, suggesting high selectivity and penetrance of Cre activation. C, RT-qPCR of CD45+ cells showing similar levels of SOCS3 expression in WT and SOCS3iEKO mice after tamoxifen treatment (Kruskal-Wallis). D, Survival curve of mice challenged with a single dose of 250 μg of endotoxin via intraperitoneal injection. Marks denote censored animals (those that were removed for diverse measurements) (Kaplan-Meier analysis). E, Severity score of mice 15-16 h after saline or LPS injection (Two-way ANOVA with Dunnett post-hoc test, het or SOCS3^iEKo^ vs control). F, Temperature difference for each mouse as measured immediately before and 15-16 h post-injection (Two-way ANOVA with Dunnett post-hoc test, het or SOCS3^iEKo^ vs control). Asterisks denote p < 0.05. Data combined from at least three independent experiments.

We had hypothesized that a loss of endothelial SOCS3 expression might mimic the effects of an IL-6 amplifier loop (10), leading to an overactivated pathway that would be associated with a worse outcome. Thus, we directly tested whether SOCS3^iEKO^ mice displayed increased IL-6 expression and signaling. In fact, we detected a dramatic increase in whole tissue IL-6 mRNA levels in lungs, kidneys and liver of these mice (Figure 2A), as well as an increase in SOCS3 mRNA itself (Figure 2B). Given that the SOCS3 primers were designed to detect full length, but not deleted SOCS3 mRNA, we speculate that non-endothelial cell types are responsible for this increase. Consistently, knockout mice show substantially increased levels of circulating IL-6, an effect that was not apparent when looking at the circulating TNF-α (Figure 2C).

**Figure 2.**
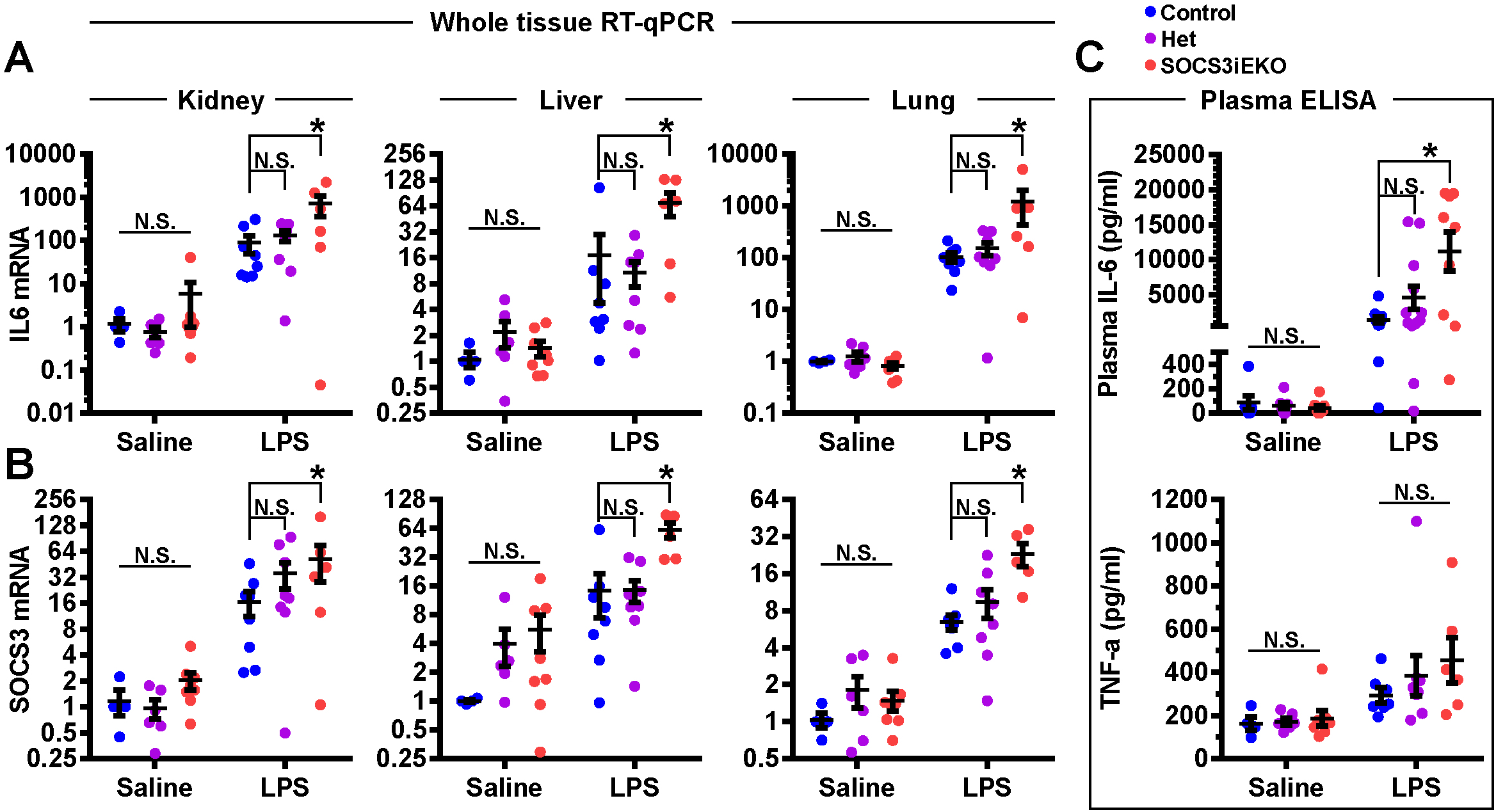
Overactivation of the IL-6/STAT3/SOCS3 pathway in endotoxemic SOCS3^iEKo^ mice. A, Levels of IL6 mRNA as measured by RT-qPCR obtained from whole organs. B, SOCS3 mRNA levels from the same sources. C, Plasma levels of IL-6 and TNF-a as measured by ELISA All data in this figure was analyzed by two-way ANOVA with Dunnett post-hoc test, comparing het or SOCS3^iEKo^ mice vs control. Asterisks denote p < 0.05. Data combined from at least three independent experiments.

### Endotoxemic SOCS3^iEKO^ mice have increased vascular permeability and multiple organ dysfunction

After the endotoxin challenge, all mice showed weight loss; however, SOCS3^iEKO^ mice consistently lost less weight than their control littermates (Figure 3A). Liquid retention is a hallmark of severe shock. Thus, we hypothesized that this reduced weight loss in SOCS3^iEKO^ mice might be due to increased edema caused by systemic vascular leak. To assess whether loss of SOCS3 increased vascular leak, we injected LPS- or saline-treated mice with FITC-dextran (70 kDa), allowed it to circulate for 30 minutes, and then euthanized the mice and perfused with PBS. Little dextran leaked from brain (Figure 3B) or lungs (Figure 3C) of control mice 15 hours post-LPS. However, dextran leak was evident in multiple spots in the organs of similarly treated SOCS3^iEKO^ mice (Figures 3B and 3C), demonstrating that loss of SOCS3 promoted a modest increase in vascular leak after endotoxin. The leak was particularly noticeable around vessels in the brain cortex and in the lung periphery.

**Figure 3.**
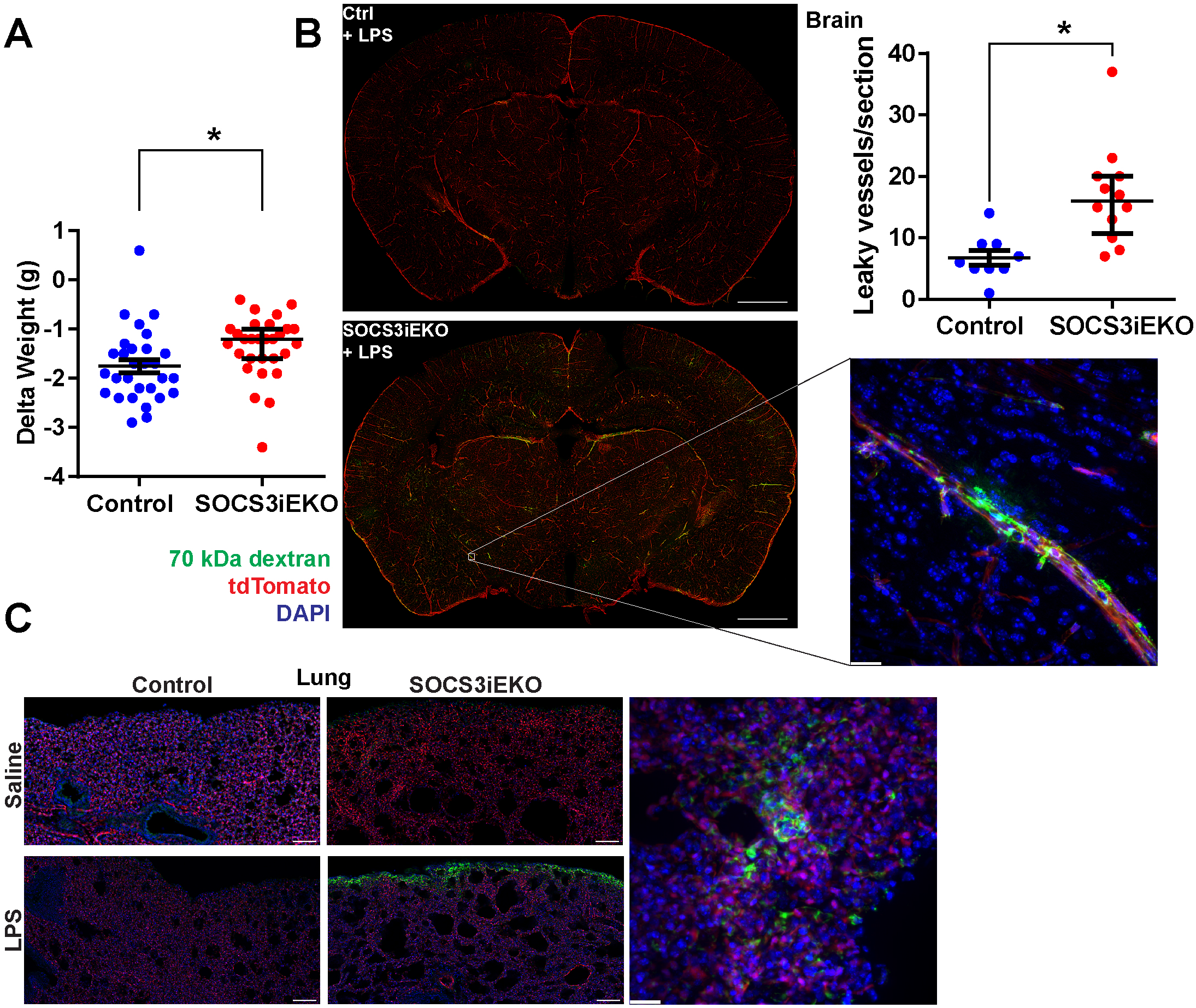
Increased vascular leakage in endotoxemic SOCS3^iEKo^ mice. A, weight difference before and 15-16 h after LPS injection (Mann-Whitney). B, Brain sections of endotoxemic mice injected with FITC-labeled 70 kDa dextran. Shown on the left are full section images (bregma ~-2 mm, bar = 1 mm) and on the right a 63x magnification of a cortical leaky vessel (maximum projection of a Z stack covering near 100 μm, bar = 20 μm). Shown in the top right is the quantification of the number of cortical vessels per section (1 section per mouse, Mann-Whitney). C, Representative regions of 50 μm thick lung sections (left, bars=100 μm) and a detail of a leaky spot near the lung edge (right, bar = 20 μm). Asterisks denote p < 0.05. Data combined from at least three independent experiments.

To better understand the mechanisms of lethality in SOCS3^iEKO^ mice, we performed a series of experiments aimed at determining the extent of organ dysfunction. Measurement of blood glucose show that endotoxin promotes a severe hypoglycemia independent of mouse genotype (Figure 4A). Surprisingly, lactate levels 15 h post-endotoxin challenge were lower in endotoxin than in saline treated mice. We attributed this finding to the severe hypoglycemia, which would limit the available source for the glycolysis needed for lactate generation. However, SOCS3^iEKO^ mice showed increased lactate levels compared to control endotoxemic mice, despite similar low glycemia (Figure 4B), suggesting higher glycolysis that may be due to increased hypoxia. Although we found a very mild lung infiltration in endotoxemic mice, loss of SOCS3 did not aggravate the histological findings (Figure 4C). Masked scoring of lung damage demonstrated no significant differences between control and SOCS3^iEKO^ (median=1 for both groups; score range from 1-normal to 4-severe pathology) endotoxemic mice. Moreover, oxygen saturation values higher than 96% at room air were consistently recorded on all animals even at an advanced stage of acute illness (Figure 4D), suggesting that lung injury cannot explain these findings.

**Figure 4.**
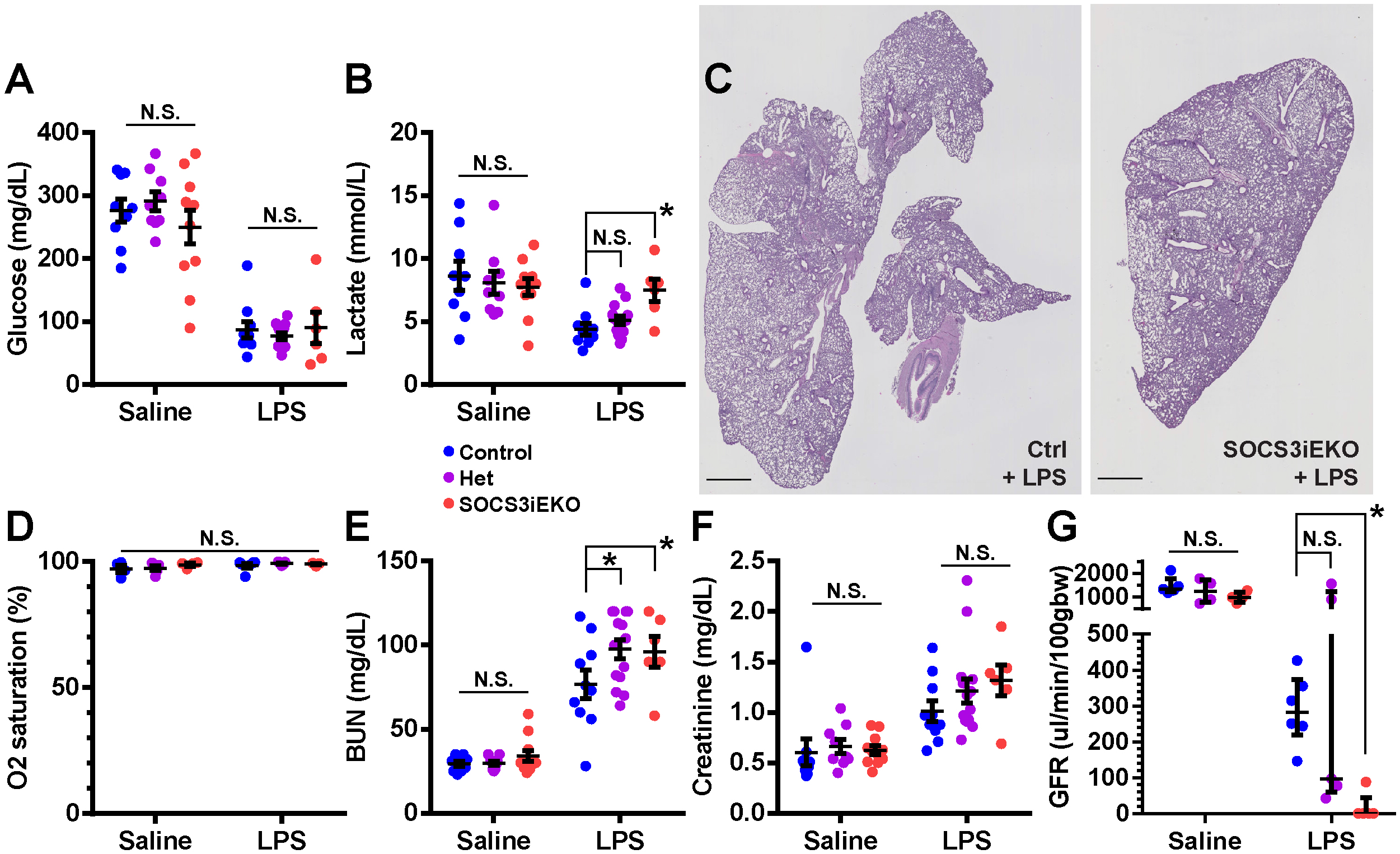
Mild lung injury and severe kidney failure in endotoxemic SOCS3^iEKo^ mice. A-B, Endotoxin-induced hypoglycemia (A) and changes in circulating lactate (B). Two-way ANOVA and Holm-Sidak post-hoc tests comparing het and SOCS3^iEKo^ mice to control within saline or LPS-treated groups. C, H&E staining of 5 μm lung sections showing mild infiltration. D, Normal blood O_2_ saturation levels in all groups (two-way ANOVA). E-F, Blood levels of urea nitrogen (E) and creatinine (F). Two-way ANOVA and Holm-Sidak post­ hoc tests comparing het and SOCS3^iEKo^ mice to control within saline or LPS-treated groups. G, Glomerular filtration rate measured for 90 minutes starting 14 h post-injection of endotoxin or saline. Two-way ANOVA and Holm-Sidak post-hoc tests comparing het and SOCS3^iEKo^ mice to control within saline or LPS-treated groups. Asterisks denote p < 0.05. Data combined from at least three independent experiments.

Liver glycogen content was severely reduced in endotoxemic mice (Supplemental Figure 2A). However, plasma albumin levels remained constant among experimental groups (Supplemental Figure 2B), suggesting that liver dysfunction is very limited in this model.

While changes in lung and liver function are minor, our studies suggest that kidney failure is the main driver of endotoxin-induced mortality in SOCS3^iEKO^ mice. Blood urea nitrogen (BUN) levels were elevated in SOCS3^iEKO^ mice compared to control mice after endotoxin (Figure 4E). Contrary to this, creatinine levels were increased by LPS, but changes were not statistically significant between SOCS3^iEKO^ and control mice (Figure 4F). Direct measurement of glomerular filtration rate (GFR) showed that, consistent with previous reports (36–38), endotoxin in control mice induced a strong reduction in kidney function (Figure 4G). Deletion of SOCS3 from the endothelium induced a much more severe response, suggesting complete kidney failure, as seen by the fact that there was no clearance of FITC-sinistrin within the 90 minutes period of this assay (Figure 4G).

### In HUVEC, SOCS3 rapid degradation limits its inhibitory function on IL-6 signaling

Our findings showed not only that endothelial SOCS3 has a critical role to sustain organ function and promote survival after an endotoxin shock, but also suggested that the regulation of SOCS3 levels critically affect its function. Thus, we set out to perform several in vitro studies to better understand how SOCS3 protein levels affected IL-6 signaling in endothelial cells. Previously, we have shown that HUVEC treated with a combination of recombinant IL-6 and sIL-6Rα (herein, IL-6+R) causes a 30% loss in barrier function (27) that is sustained for at least 24 hours. To assess if sustained signaling downstream of IL-6 receptor activation was required to maintain the barrier function loss, we treated HUVEC with IL-6+R for 6 h to allow for full barrier function loss prior to adding a blocking anti IL-6 antibody or non-specific IgG. As shown in Figure 5A, IL-6 inhibition completely reversed IL-6+R-induced increase in monolayer permeability. Similarly, addition of the JAK inhibitor ruxolitinib (rux) was sufficient to rescue the full barrier function in HUVEC even when added up to 24 h after the IL-6+R challenge (Figure 5B). Consistent with a sustained IL-6+R signaling throughout this long period, we detected large increases in SOCS3 and IL-6 mRNA levels 24 h and 48 h after IL-6+R treatment (Figure 5C).

**Figure 5.**
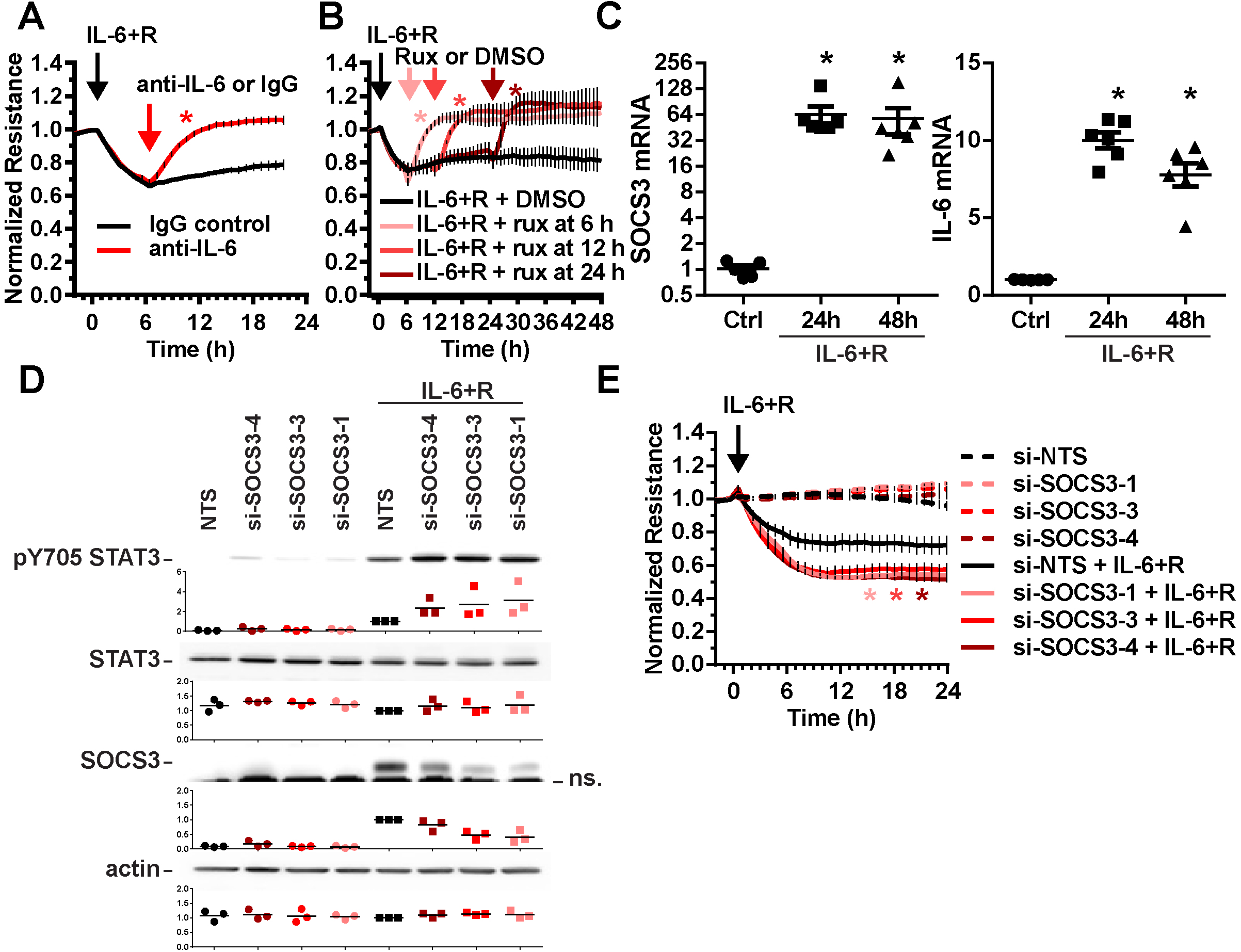
SOCS3 moderately reduces the effects of sustained IL-6 signaling in HUVEC. A-B, HUVEC treated with a combination of recombinant IL-6 and sll-6Ra displayed reduced barrier function that quickly recovered after antibody-mediated IL-6 blockade (A) or treatment with the JAK inhibitor ruxolitinib (B). Two-way ANOVA of repeated measurements and Dunnett post-hoc tests comparing antibody (A) or ruxolitinib (B) groups to the respective vehicle controls. C, IL-6-induced transcriptional response is sustained for at least 48 h (one-way ANOVA and Dunnett post-hoc test comparing IL-6+R treated groups to control. D, Western blot analysis showing a moderately increased level of STAT3 phosphorylation at tyrosine 705 in SOCS3 siRNA-treated cells compared to cells transfected with a non-targeting sequence (NTS). Levels of SOCS3 knockdown are shown above a non-specific band (ns). E, SOCS3 knockdown promotes a further loss of IL-6+R-induced barrier function loss but does not affect basal resistance levels. Two-way ANOVA of repeated measurements and Dunnett post-hoc tests comparing siSOCS3-transfected cells to NTS in the presence of IL-6+R. Asterisks denote p < 0.05. Data representative of at least three independent experiments.

Given that SOCS3 remains elevated even as IL6+R is causing a sustained barrier loss, we questioned whether SOCS3 has a protective, anti-inflammatory function in HUVEC. To assess this, we decreased SOCS3 expression in HUVEC by utilizing three individual siRNA sequences, followed by treatment with IL-6+R. As shown in Figure 5D, SOCS3 knockdown led to a minor, albeit consistent, increase in IL-6+R-induced STAT3 phosphorylation at tyrosine 705, demonstrating that SOCS3 can act as a weak negative regulator of this pathway. Moreover, SOCS3 depletion further exacerbated the barrier function loss induced by IL-6+R without having any effect in the absence of IL-6+R (Figure 5E).

While the above data demonstrated that SOCS3 was able to reduce IL-6+R signaling, it also showed that this inhibition was not robust. Given that prior reports showed that SOCS3 has a short protein life span due to ubiquitin-mediated proteasomal degradation (39, 40), we hypothesized that fast ubiquitination and degradation would not allow for sufficient SOCS3 protein accumulation needed for a more robust inhibition of the pathway, despite the very high mRNA levels. To test this, we pretreated HUVECs with the proteasome inhibitor MG-132, followed by an IL-6+R challenge. MG-132 pre-treatment effectively led to a robust increase in SOCS3 protein levels after IL-6+R treatment (Figure 6A). Importantly, the same protein lysates showed a strong inhibition of STAT3 phosphorylation two hours after challenge, suggesting that increased SOCS3 protein levels can act as a strong negative inhibitor of IL-6+R signaling (Figure 6A). We then determined the rate of turnover of SOCS3 by a cycloheximide (CHX)-mediated pulse and chase experiment. We treated cells with IL-6+R for 2 h (to allow for SOCS3 expression), followed by the addition of CHX or PBS 2-30 minutes prior to lysing. SOCS3 protein expression was measured by Western blot. We found that SOCS3 turnover is extremely rapid in these cells, with a half-life between 10 and 20 minutes (Figure 6B).

**Figure 6.**
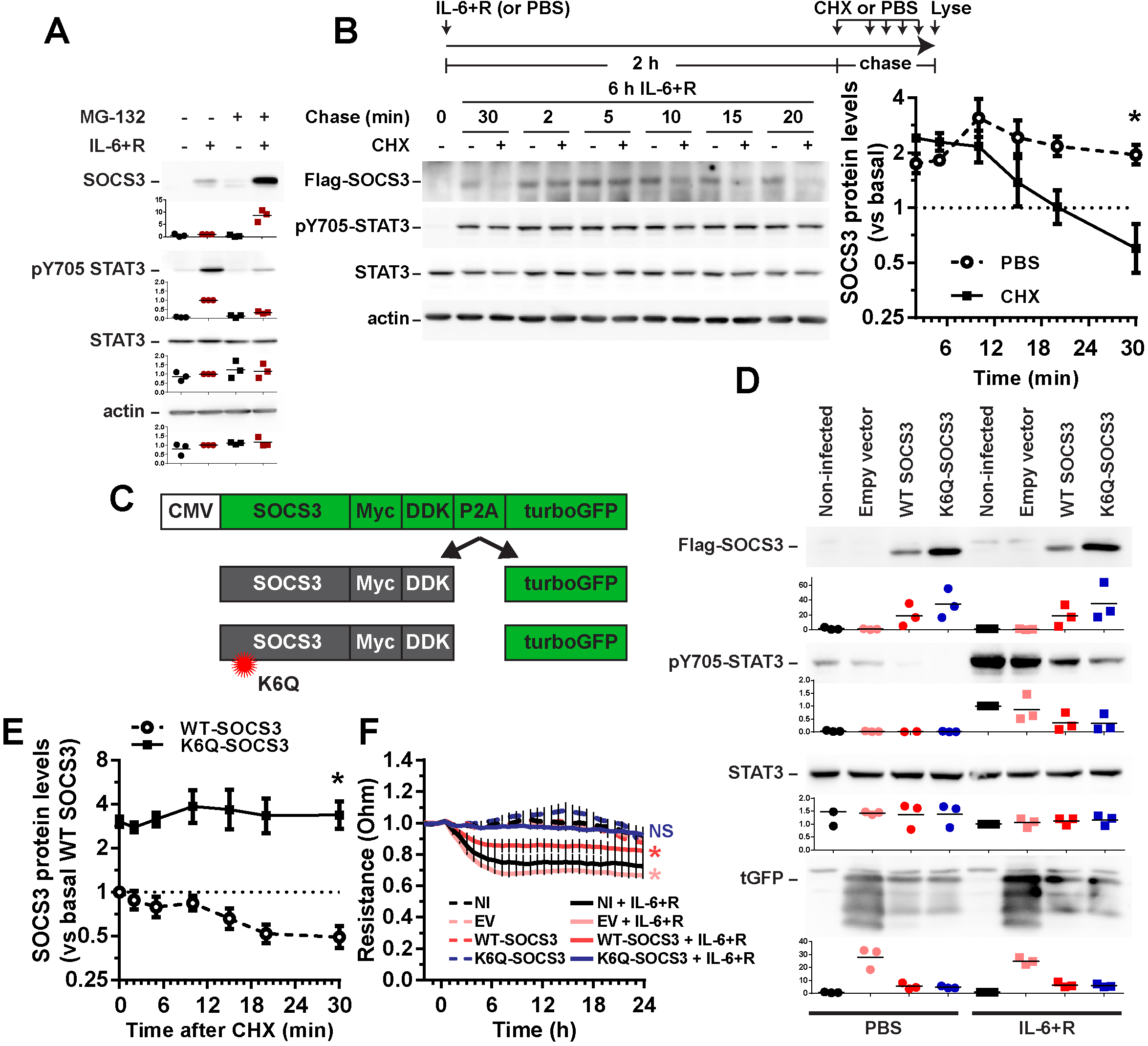
Levels of SOCS3 protein are rapidly degraded by the ubiquitin-proteasome pathway. A, Western blot analysis showing SOCS3 expression and STAT3 phosphorylation levels 2 h after treatment with a combination of recombinant IL-6 and sll-6Ra (IL-6+R) in the presence or absence of the proteasome inhibitor MG-132. B, Pulse-chase experiment of cells treated for 6 h with IL-6+R and then for the specified time with either PBS or the protein synthesis inhibitor cycloheximide prior to lysis. Representative Western blots (left) and quantification of three independent experiments (right). Two-way ANOVA with Dunnett post-hoc test (CHX vs PBS). C, Diagram of the SOCS3 overexpression strategy. D, Western blots showing the expression levels of exogenous SOCS3 and their effects on IL-6+R-induced STAT3 phosphorylation. E, Quantification of pulse-chase experiments by measuring the levels of Flag-SOCS3 after cycloheximide. Two-way ANOVA with Dunnett post-hoc test (K6Q vs WT SOCS3). F, TEER measurements of monolayers expressing the different constructs and treated with either PBS or IL-6+R. Two-way ANOVA of repeated measurements with Dunnett post-hoc test (vs PBS-treated, non-infected cells). Data representative of at least three independent experiments.

We then sought to directly test the hypothesis that SOCS3 ubiquitination limited its function in HUVEC. We generated lentiviral vectors to overexpress tagged versions of either wild-type SOCS3 or a degradation resistant mutant (K6Q-SOCS3) (40). The SOCS3 construct was joined to turboGFP (tGFP) via a P2A linker (41), thus allowing for expression of equimolar quantities of SOCS3 and free tGFP (Figure 6C). As expected, the mutation at lysine 6 led to higher accumulation of the exogenous SOCS3 protein even in the absence of IL-6+R (Figure 6D), that was associated with increased stability of this mutant (Figure 6E). Moreover, overexpression of K6Q-SOCS3 led to a strong reduction of IL-6+R-induced STAT3 phosphorylation, whereas overexpression of WT-SOCS3 promoted a more modest reduction (Figure 6D). Consistently, exogenous WT-SOCS3 reduced IL-6+R-induced barrier function loss, while K6Q-SOCS3 overexpression prevented barrier loss altogether (Figure 6F).

### IL-6 induced a pro-inflammatory gene expression profile in HUVEC involving a type I interferon-like response

We had previously shown that inhibition of de novo mRNA or protein synthesis prior to an IL-6+R challenge prevented the loss of barrier function (27). Because of the need for sustained signaling (Figure 5), we determined whether sustained protein synthesis was also required. CHX treatments up to 4 h post IL-6+R challenge completely reversed the barrier function loss (Supplemental Figure 3A). Thus, we sought to determine the transcriptional changes that occurred during this time frame by performing an RNA-Seq analysis of cells treated or not for 3 h with IL-6+R, in the presence or absence of rux. We identified 275 genes that were significantly (adjusted p<0.001) induced (>2-fold) and 64 significantly repressed genes by IL-6+R. The top 50 most significant genes are shown in Figure 7A. Of note, JAK inhibition completely abolished the IL-6+R response (1 gene upregulated, 4 genes downregulated vs DMSO). We performed RT-qPCR validation experiments on cells treated with IL-6+R for up to 24 hours, as well as in STAT3 knockdown cells treated with IL-6+R for 1 or 6 hours on over 30 genes of the top 100 induced gene set (Figure 7B, Supplemental Figure 3B and data not shown). Bioinformatic analysis of these data using the ISMARA package (42) shows that this IL-6+R treatment induced the expected enrichment of STAT3-responsive genes, but also a strong type I interferon (IFN)-like response (Figure 7C). STRING (43) analysis shows tight network of functional interactions linking multiple transcription factors of the STAT and IRF families (Figure 7D). Consistently, IL-6+R promoted an increase in IFN-responsive genes that was abrogated by overexpression of SOCS3 (Figure 7E). This appears to be due to a direct effect of STAT3 rather than an indirect mechanism through IFN autocrine signaling, since IL-6+R or SOCS3 overexpression did not induce any significant change in type I interferons expression (Supplemental Figure 3C) and the levels of IFNα, IFNβ and IFNγ in the conditioned medium of IL-6+R-treated HUVEC remained below the level of detection as measured by ELISA (data not shown).

**Figure 7.**
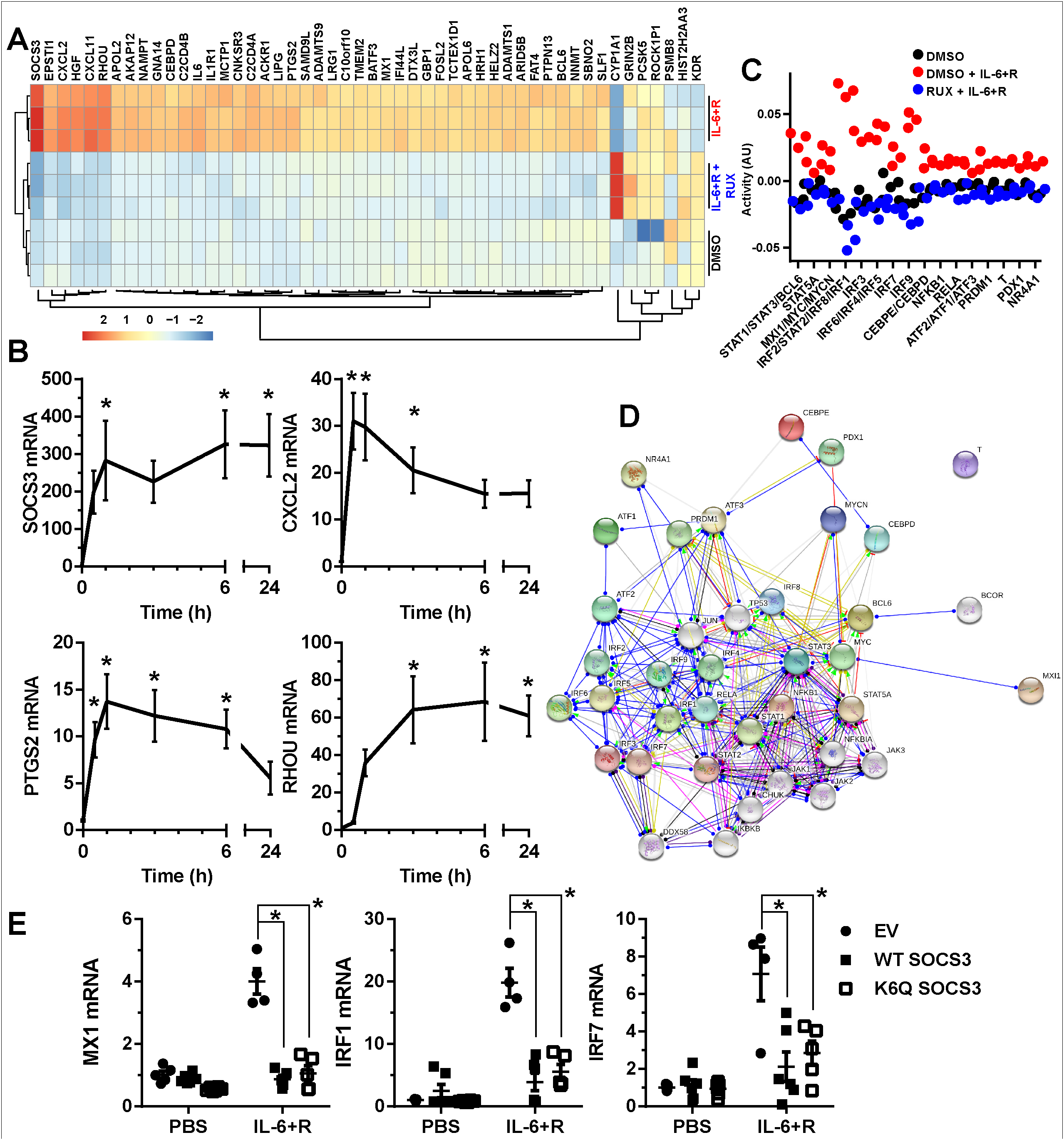
IL-6-induced gene expression profile in HUVEC. A, Heatmap and unbiased clustering of cells treated with either PBS or with a combination of recombinant IL-6 and sll-6Ra (IL-6+R) after 30 min incubation with either ruxolitinib (rux) or DMSO (vehicle control). Ribosomal RNA-depleted RNA was assayed by RNA-sequencing. Shown are the 50 most significant differences between IL-6+R and PBS, sorted by log2 fold difference. One experiment performed in triplicate. B, RT-qPCR analysis of an IL-6+R time course to confirm the changes in mRNA expression of selected candidate genes. One-way ANOVA and Dunnett post-hoc (vs no IL-6+R). Combined data from three independent experiments performed in duplicate each. C, ISMARA analysis of gene expression changes using the FASTQ files obtained in A showing a significant enrichment in type I interferon responses. D, STRING representation of the main interconnections of genes identified in C. E, RT-qPCR of cells treated with or without IL-6+R after infection with lentivirus to overexpress WT SOCS3 or K6Q-SOCS3. An empty vector lentivirus was used as control. Two-way ANOVA and Dunnett post-hoc tests. Combined data from three independent experiments performed in duplicate each. Asterisks denote p < 0.05.

### Endotoxin-induced transcriptional profiling in SOCS3^iEKO^ mice demonstrates a pro-adhesive, pro-thrombotic phenotype associated with a type I IFN-like response

To determine whether a similar transcriptional profile occurred in SOCS3^iEKO^ endotoxemic mice, mice were challenged for 15 h with or without endotoxin prior to euthanasia followed by mRNA isolation from lungs, kidneys and livers. We then assayed the expression of over 50 genes from each organ by RT-qPCR. De novo clustering of these data generated three distinct groups of mice (Figure 8A). Unsurprisingly, mice treated with saline solution clustered separate from those treated with LPS, with the exception of one LPS-treated heterozygous mouse that did not show any symptoms following the LPS injection. All other LPS-treated mice clustered in the two other groups: one included most control mice, while the other was comprised mostly of SOCS3^iEKO^ mice, demonstrating that the differences in the severity of the response closely correlated with their gene expression profile. Consistent with an intermediate response, both clusters contained several SOCS3 heterozygous mice. We then performed a principal component analysis (PCA) of all the data obtained from these mice, including the gene expression data and all physiologic parameters (temperature, weight loss, blood counts, plasma IL-6 levels, creatinine, BUN, lactate, etc.). As shown in Figure 8B, all endotoxemic SOCS3^iEKO^ mice clustered separate from LPS-treated control mice, with most heterozygous again with intermediate changes. A functional assessment of each parameter in the first two PCA dimensions suggests that many transcriptional changes found in the kidneys drive a different phenotype than changes in either lungs or livers (Figure 8C). The severity score, BUN, creatinine, and plasma IL-6 effects, however, were more closely associated with lung and liver changes. To further investigate how gene expression changes were associated with pathophysiologic parameters, we performed a complete cross-correlation analysis of all the data obtained from each endotoxemic mouse (Supplemental Figure 4A). We observed multiple clusters with close cross-correlation, including two large clusters that had a high correlation with the severity score (all significantly associated changes are shown in Supplemental Figure 4B). Notably, the levels of plasma IL-6 showed the highest correlation with the severity score, closely followed by BUN levels and the degree of temperature loss. The expression of lung PLAUR, liver ICAM and kidney RHOU were the transcriptional changes with the highest correlation with the severity score and plasma IL-6 (Supplemental Figure 4B). Interestingly, RHOU was also dramatically increased by an IL-6+R treatment in HUVEC (Figure 7). Kidney injury markers, such as SYNPO, also significantly correlated with the severity score, albeit with a much lower R value (Supplemental Figure 4C).

**Figure 8.**
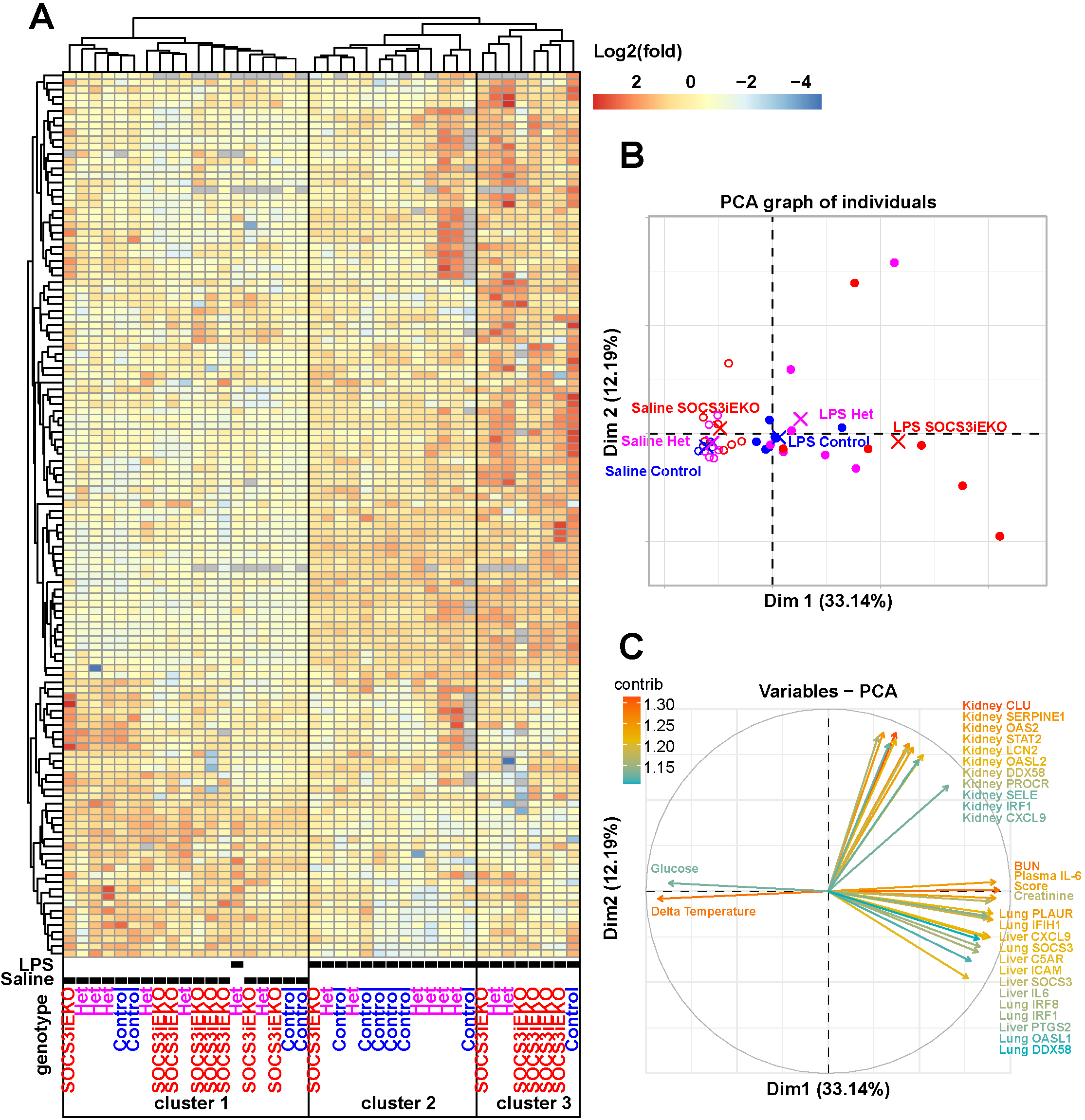
Loss of endothelial SOCS3 aggravates the transcriptional response to endotoxin in multiple organs. A, Heatmap and unbiased clustering of mice 15-16 h after injection of endotoxin based on gene expression changes as measured by RT-qPCR from whole organ RNA (lungs, livers and kidneys) and transformed through a base 2 logarithm. Data combined from three independent experiments. B-C, Principal component analysis of the raw expression data combined with all physiological parameters. Shown are the first two dimensions demonstrating a differential clustering of SOCS3^iEKo^ mice (B) and the main factors weighed in these two dimensions (C). Log2 gene expression data is available as Supplemental Table 6 and all data including physiological parameters as Supplemental Table 7.

In agreement with a strong type I IFN-like response observed in IL-6+R-treated HUVEC, all organs examined showed a dramatic increase in many IFN-responsive genes (including MX1, IRF8 and OASL1) in endotoxemic mice, which was further aggravated by the loss of endothelial SOCS3 (Figure 9A and Supplemental Figure 5). This increased IFN-like transcriptional response was also associated with altered mRNA levels coding for leukocyte adhesion molecules and with a pro-thrombotic phenotype (Figures 9B and 9C and Supplemental Figure 5). Notably, P-selectin expression was increased nearly 100-fold in the kidneys of endotoxemic SOCS3^iEKO^ mice. Similarly, tissue factor expression (coded by the F3 gene) increased only two-fold in control mice but over 16-fold in SOCS3^iEKO^ mice, suggesting a severe dysregulation of the leukocyte adhesive and thrombomodulatory capacity of the endothelium of these knockout mice (Figure 9C). Immunofluorescence staining confirmed the increased expression of P-selectin in every organ assessed of endotoxemic SOCS3^iEKO^ mice (Figure 10). Moreover, in these mice, P-selectin was exposed on the luminal surface of the endothelium and not stored in Weibel-Palade bodies, suggesting that the transcriptional changes indeed led to an increased adhesivity to leukocytes of the affected vasculature.

**Figure 9.**
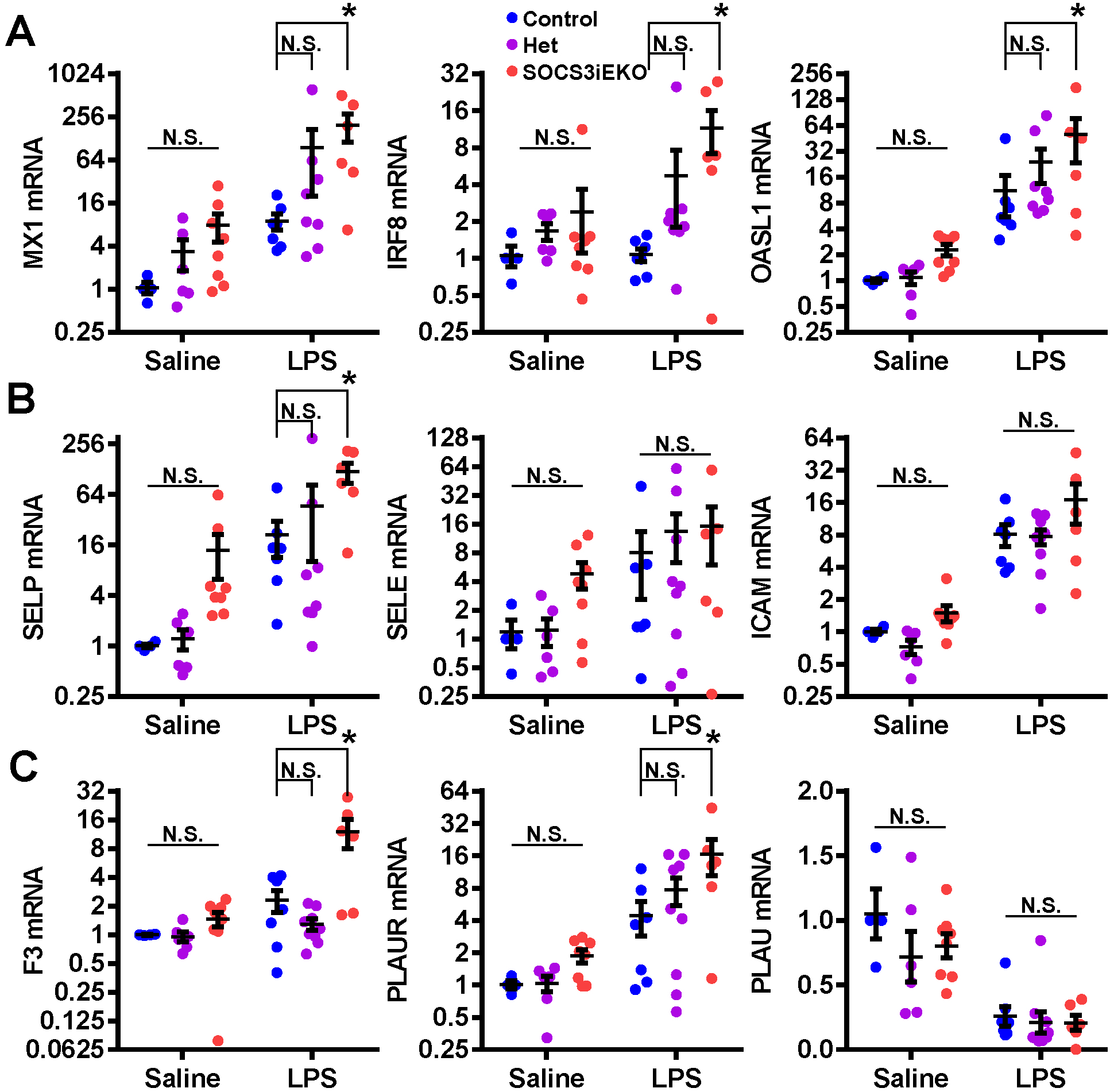
Transcriptional changes in whole kidneys of mice treated with endotoxin in the presence or absence of endothelial SOCS3. Loss of SOCS3 promotes a type I IFN-like response (A), as well as adhesive (B) and prothrombotic (C) changes after endotoxin. Two-way ANOVA and Holm-Sidak post-hoc tests comparing het and SOCS3^iEKo^ mice to control within saline or LPS-treated groups. Data combined from three independent experiments. Asterisks denote p < 0.05.

**Figure 10.**
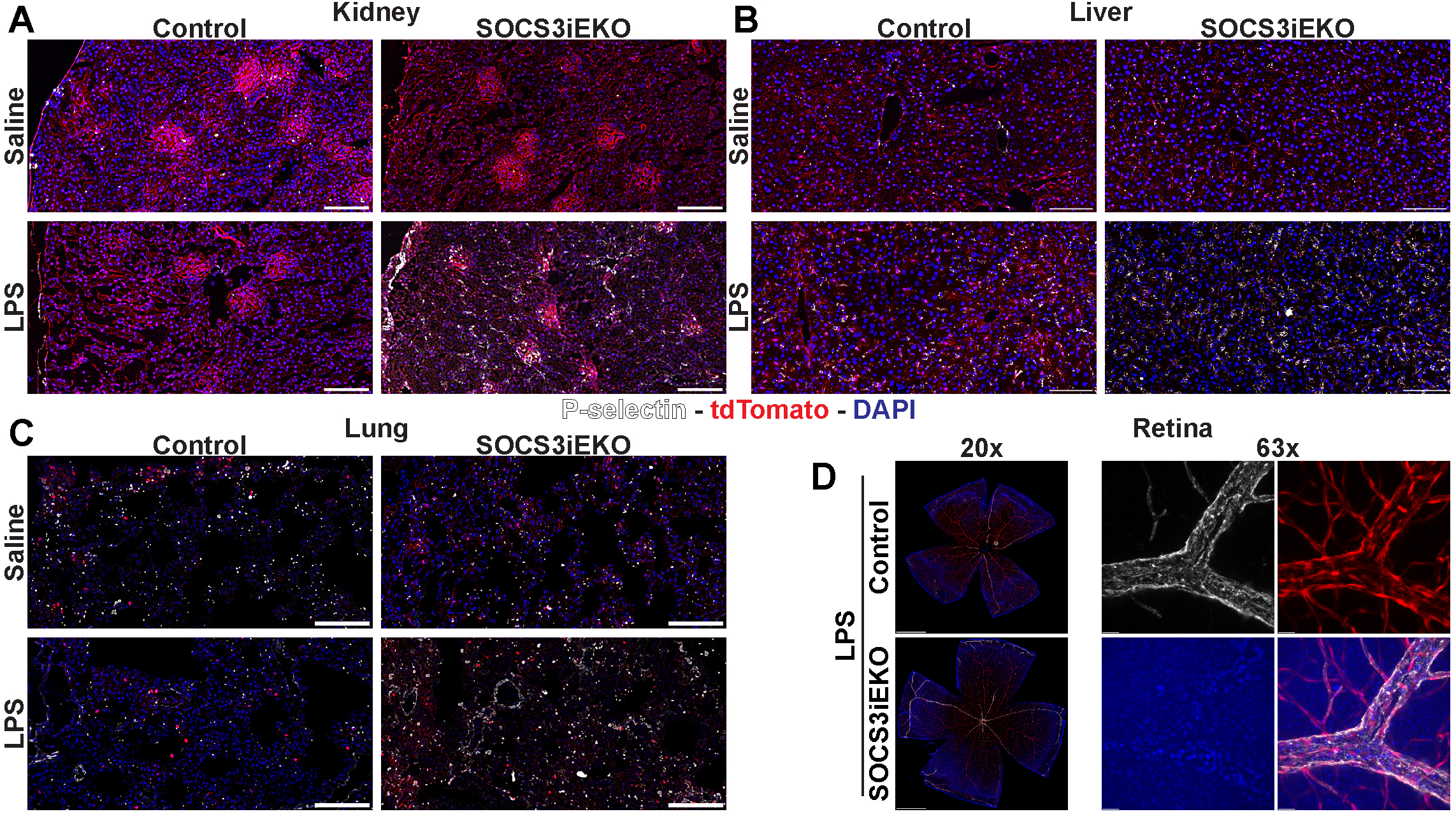
Loss of SOCS3 promotes an increase in P-selectin expression after endotoxin. A-C, Representative images of thin sections of kidney (A), liver (B) and lung (C) stained with anti P-selectin (white) and DAPI (blue). tdTomato expression is shown in red. D, Whole retina flat mounts (left) and a detail of a vein shown as a maximum projection of a Z stack. Bars, 100 μm (A-C), 500 μm (D, left) or 20 μm (D, right). Representative images of three independent experiments.

Consistent with these findings, we observed accumulation of intravascular leukocytes in multiple tissues. We identified a large number of leukocytes adhered to the vessels of kidneys and lungs, as determined by H&E staining of FFPE sections (Figure 11A). Flow cytometry analysis of lungs showed a >2-fold increase in neutrophils upon treatment with LPS regardless of genotype (Supplemental Figure 6A). Consistent with the low levels of tdTomato+ cells in blood and bone marrow (Figure 1), few Ly6G+ cells were positive for tdTomato in lungs (Supplemental Figure 6A). Histochemical staining of FFPE sections for myeloperoxidase (MPO) or F4/80 shows that both neutrophils and monocytes accumulated within the vessel lumens in lungs and kidneys (Supplemental Figure 6B). To better interrogate the extent of this intravascular accumulation, we obtained flat mounts of the retinas from these mice, which allowed us to study the whole tissue vasculature. Staining with an anti-CD45 (pan leukocyte) antibody demonstrated a dramatic accumulation of intravascular leukocytes (tdTomato-; CD45+) that, in most cases, filled all retinal venules and colocalized with E-selectin expression within the vessel lumens (Figure 11B). This accumulation was not evident on retinal arterioles or capillaries, further suggesting that this is due to increased adhesion to venules and not simply due to increased circulation of leukocytes within the retinal vasculature.

**Figure 11.**
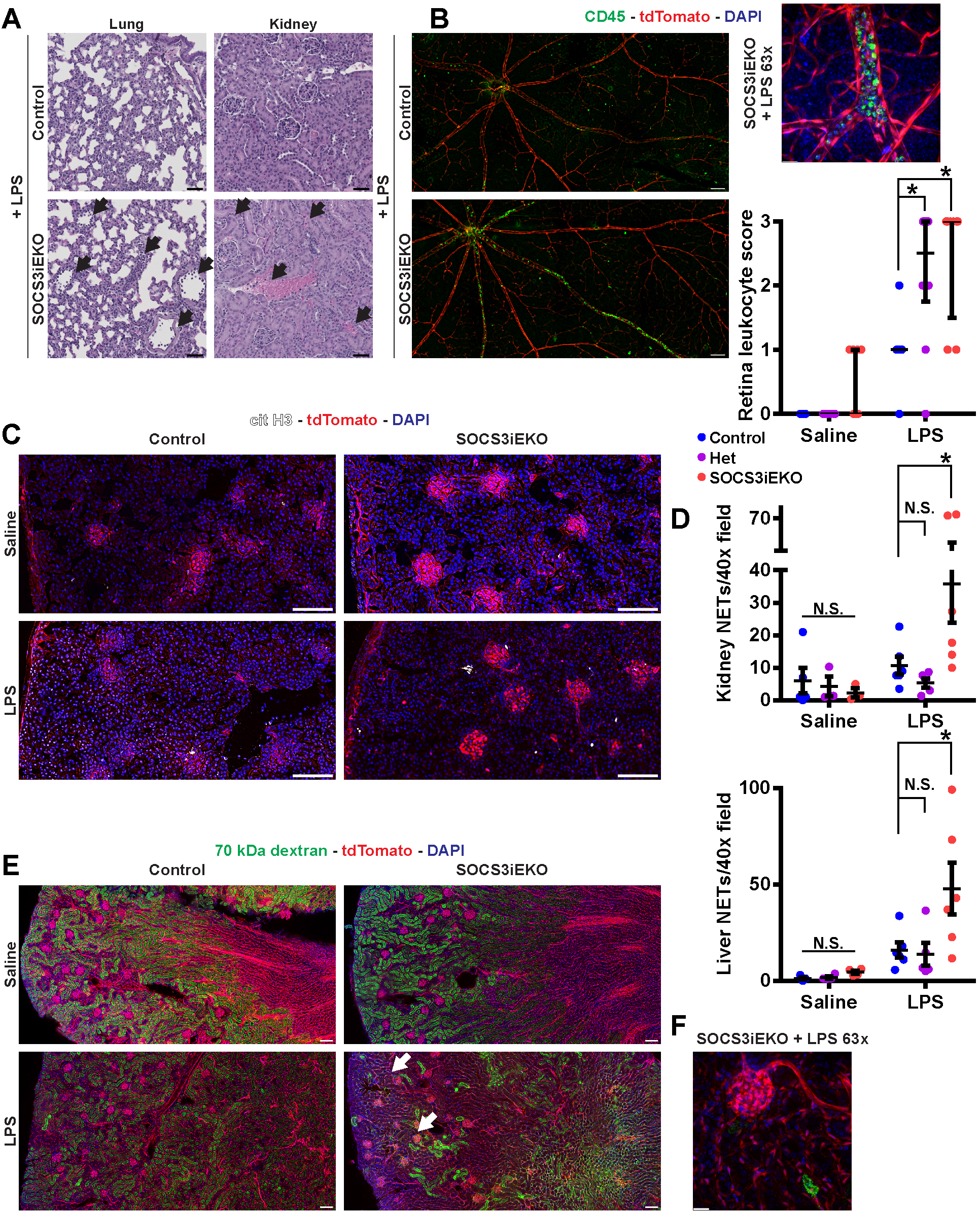
Increased intraluminal leukocyte accumulation and NETosis in endotoxemic SOCS3^iEKo^ mice. A, Representative H&E images of thin FFPE sections of lungs and kidneys of LPS-treated control or SOCS3^iEKo^ mice. Arrows point to regions of leukocytes adhered to the vessel walls. B, Whole retina flat mounts were stained with CD45 (green) and counterstained with DAPI (blue). tdTomato is shown in red. Left, 20x tiled images (bar= 100 μm). Top right, detail of a vein showing dramatic leukocyte accumulation (bar = 20 μm). Lower right, scoring of three independent experiments (0: No intravascular CD45+ cells. 1: Occasional (<10 in total) CD45+ cells. 2: Multiple cells in> 1 vein, occasional small clump. 3: >3 veins compromised, several clumps> 10 cells). Two-way ANOVA and Dunnett post-hoc tests comparing het and SOCS3^iEKo^ mice to control within saline or LPS-treated groups. Asterisks denote p < 0.05. C, Representative images of 5 μm kidney sections stained with an antibody against citrullinated histone H3 (cit H3). Blue, DAPI and red, tdTomato (bar = 100 μm). D, Quantification of data from 5 μm kidney and liver sections from three independent experiments. Mann-Whitney test. E, Representative images of thick (50 μm) kidney sections from mice injected with FITC-labeled 70 kDa dextran showing a reduction of green fluorescence intensity in control LPS mice and regions with convoluted tubules devoid of fluorescence in endotoxemic SOCS3^iEKo^ mice (white arrow). F, A detail of a glomerulus and surrounding tubules lacking any dextran in endotoxemic SOCS3^iEKo^ mice (bar = 20 μm). Representative images of three independent experiments.

### Increased NETosis is associated with organ injury in SOCS3^iEKO^ mice

Intravascular neutrophil extracellular traps (NET) are commonly associated with increased tissue damage and vascular dysfunction (44, 45). Thus, we next questioned whether loss of endothelial SOCS3 may lead to increased NETosis upon LPS, by staining frozen sections for citrullinated histone H3. As shown in figure 11C, LPS-induced increase in kidney NET formation in control mice was largely augmented in SOCS3^iEKO^ mice, supporting the notion that the accumulated intravascular leukocytes may have a causal effect in promoting tissue injury. NETs were found within the glomeruli and surrounding convoluted tubules. We observed a similar increase in NETosis in SOCS3^iEKO^ livers. Quantification of liver and kidney NETosis is presented in Figure 11D. Increased leukocyte adhesion, thrombosis and NET formation may lead to reduced kidney perfusion, explaining the observed lack of kidney function in endotoxemic SOCS3^iEKO^ mice. Kidneys from these mice show a dramatically reduced accumulation of 70 kDa FITC-dextran within the convoluted tubules 30 minutes after inoculation, with many regions lacking any fluorescence (Figure 11E-F), suggesting lack of perfusion in these areas.

## Discussion

Here, we present evidence to support the notion that endothelial SOCS3 is a crucial limiting factor for IL-6 signaling, promoting vascular homeostasis and survival during critical illness. An amplified JAK/STAT3 signaling in the endothelium caused severe vasculopathy, including a proinflammatory transcriptional profile leading to increased adhesivity and thrombogenicity of the endothelial surface, that is associated with a strong type I IFN-like response. This, in turn, leads to leukocyte accumulation within the affected vascular lumens, NETosis, and kidney failure. Mechanistically, the exact levels of SOCS3 are maintained by a balance between a large transcriptional increase and very fast protein degradation via ubiquitination of lysine 6 and proteasomal degradation. Overall, the findings of this study reinforce the critical role of the endothelium in acute, systemic inflammation and point to a pivotal role for SOCS3 stabilization as a therapeutic target.

An overactivated endothelium is key in the body’s unmanaged response during inflammation and MODS (3–6, 46, 47), and recently, COVID-19 (48, 49). Thus, there is an urgent need to understand critical signaling pathways that influence the endothelium to respond in this manner. Our research focused on a signaling mechanism that we hypothesized may act as the tipping point between a beneficial immune response and an un-controlled response that increases the risk of death or long-term sequelae. The circulating levels of IL-6 are not only closely associated with APACHE II and SOFA scores in septic patients, but also are very valuable as a prognostic marker of shock mortality (9–15). The murine model used here uses a single endotoxin injection to induce a severe systemic inflammatory reaction that is not lethal in control mice but led to MODS in mice lacking SOCS3 in the endothelium, at least in part through vascular dysfunction including thrombosis, leukoembolization and intravascular NETosis. These defects, in turn, resulted in systemic vascular leakage, organ hypoperfusion and ultimately organ failure and death, thus closely mimicking the disease progression of critically ill patients (3–5).

Current published data argues for both pro- and anti-inflammatory activities of endothelial STAT3. An increase in endothelial STAT3 phosphorylation is commonly associated with the inflammatory response (50–58). For example, LPS promotes a rapid increase in endothelial STAT3 phosphorylation (55–58) and the expression of IL-6Rα (59) and JAK3 (60). Confirming a critical role for this pathway in the inflammatory response, JAK or STAT3 pharmacological inhibitors improved survival of septic mice and rats (61–64) and a JAK inhibitor reduced ICAM-1 expression and vascular leakage in myocardial endothelium of LPS-injected mice (65). In contrast, disruption of STAT3 in the endothelium and hematopoietic cells using a Tie2-Cre transgene driving STAT3^fl/fl^ excision led to a marked intestinal infiltration after an LPS challenge (66), lack of dendritic cell development (67) and increased myocardial apoptosis after ischemia/reperfusion injury (68). Using a similar approach with a Tie2e-Cre driver that shows increased endothelial specificity (69), it was shown that endothelial STAT3 ablation leads to increased susceptibility to LPS (69), hyperoxia-induced lung injury (70) and hepatic inflammation after ethanol ingestion (71). These findings demonstrate unequivocally that STAT3 expression is critical to sustain endothelial function but contradict the pharmacological data discussed above. Thus, we sought to determine the effect of STAT3 overactivation in the context of systemic inflammation. To achieve that, we took advantage of the negative regulation through expression of SOCS3 (10, 18). SOCS3 is integral to survival, since homozygous germline deletion of SOCS3 leads to embryonic lethality between days E11 and E13, mainly due to placental defects (72). Further, removal of SOCS3 from varying cell types using various tissue-specific Cre drivers demonstrated important, non-redundant roles in hepatocytes, myeloid cells, all hematopoietic cells, and others through increased signaling in response to inflammatory mediators and other stimuli (73), further suggesting that STAT3 overactivation has deleterious effects. Here, we show that removal of SOCS3 from the adult endothelium creates a primed endothelium that is more reactive to a pro-inflammatory insult. Moreover, our in vitro data suggests that inhibition of endothelial SOCS3 degradation is a promising avenue to prevent and/or revert vasculopathy during shock.

The data from this study indicates that removal of endothelial SOCS3 may indeed be one of the contributors to critical illness, at least in part by promoting an IL-6 amplifier mechanism, similar to that previously linked to worse prognosis (10). We noted that endotoxin treated SOCS3^iEKO^ mice became extremely hypothermic and this physiologic change was statistically linked to the severity score and progression of inflammation. When comparing blood panels of these mice, versus their control counterparts, we were surprised to see that the levels of circulating leukocytes and platelets were not different between the groups. Histochemical, functional, and transcriptomic analyses demonstrate that loss of SOCS3 led to systemic alterations in multiple organs. Histological assessment showed modestly increased numbers of leukocytes in the lung interstitium, but not a gross increase in edema, suggesting that lymphatic draining was sufficient to balance the increased vascular permeability we observed in the fluorescent dextran experiments. Given the lack of overt histological changes in lungs and normal blood oxygen levels, it is clear that the intraluminal accumulation per se was not sufficient to induce tissue damage. This finding is consistent with the normal O_2_ saturation we measured even in severely compromised mice, demonstrating that the lethality seen in endotoxemic SOCS3^iEKO^ mice was not due to respiratory distress. Similar findings were previously reported in a model of cecal ligation and puncture (74). Similarly, we observed a loss of glycogen, but no overt histological changes in liver sections. This finding, together with similar plasma albumin in all experimental groups, suggest that liver function was mostly maintained. Again, it has been previously shown that sepsis does not induce severe liver dysfunction (38), but instead, acute kidney injury. In fact, SOCS3^iEKO^ mice showed severe kidney failure in response to LPS, as determined by direct glomerular filtration rate assays, and confirmed by increased BUN levels, altered expression of kidney injury markers, and increased NETosis; increased NET formation has been previously associated with kidney dysfunction (75). How NETs form in this situation remains to be determined. Given low amount of tdTomato positive circulating cells (arguing against a direct effect of SOCS3 loss in neutrophils), we propose that signals derived from the endothelium (either soluble or through cell-cell contacts) are the most likely cause. Surprisingly, we also found massive leukoembolization in retinal venules, and increased vascular permeability in large vessels irrigating the brain cortex. Although the blood-brain barrier is tightly regulated, it is often compromised in such disease states as Parkinson’s, Alzheimer’s, and acute inflammation (76). Importantly, many critically ill patients exhibit mental confusion and delirium (77–79), which is suggested to be due to impaired microvascular perfusion (80). Moreover, ICU survivors have increased risk of long-term cognitive impairment, which is correlated with the severity of acute illness (81). Although the fast lethality in this model did not allow for any assessment of cognitive function in these mice, this finding suggests a potential mechanism that would explain short-term and long-term cognitive dysfunction via localized edemagenic events driving brain damage.

Tissue perfusion in septic patients is drastically reduced, and microvascular changes are independent predictors of sepsis mortality (82–84). Many of our findings suggest that localized hypoperfusion was a main driver of lethality in the knockout mice. Lactate measurements after endotoxin in control mice were surprisingly lower than baseline levels. We hypothesized that this could be explained by the severe hypoglycemia induced by the endotoxin shock. However, SOCS3^iEKO^ mice showed much increased lactate levels compared to control mice, even if plasma glucose levels remained similarly low. This may suggest that there may be regions with increased hypoxic conditions. Consistent with this finding, we found that fluorescent dextran was unable to reach many regions of the kidney cortex in endotoxemic SOCS3^iEKO^ mice, demonstrating that, at least in the most severely affected organ, tissue hypoperfusion and thus localized hypoxia are features of this model. We did not observe a lack of perfusion of either lungs or livers, consistent with their increased function.

In our studies, we detected dramatically increased leukocyte adhesion in endotoxemic SOCS3^iEKO^ mice as measured by mRNA and protein levels of leukocyte receptors, as well as direct binding of leukocytes, including neutrophils and monocytes, to the vessel lumens. This increase is similar to that observed in septic patients (7, 85, 86). This was particularly evident in the retinal bed vasculature, where we detected increased E- and P-selectin levels, as well as dramatic leukoembolization throughout the venular system. Little is known about the involvement of the retinal microvasculature during shock. Notably, one study demonstrated a high incidence of retinal angiopathies in septic patients (87). Whether similar leukoembolization occurs in critically ill patients remains to be determined. In addition to this increase, SOCS3^iEKO^ mice had higher levels of pro-thrombotic factors over those seen in the endotoxic control mouse. In fact, tissue factor (F3) mRNA was greatly increased, notably in the kidney. This, along with reduced amounts of PLAU mRNA (coding for the urokinase plasminogen activator), led us to hypothesize that an increase in clotting may further aggravate the kidney hypoperfusion. We also detected increased levels of PLAUR expression (coding for uPAR). Although we do not know the source(s) or function of this change, it is consistent with clinical findings which demonstrate that circulating soluble uPAR levels are associated with systemic inflammation severity (88) and produced by neutrophils (89). The hypothesis that intraluminal neutrophils in endotoxemic SOCS3^iEKO^ mice may drive a hyper-coagulatory state in this model remains however to be tested.

In vitro studies using HUVEC showed that IL-6 signaling leads to a strong type I IFN-like response. Notably, we found that loss of endothelial SOCS3 greatly increased the expression of multiple IFN-responsive genes in all organs tested. Moreover, the expression of many of these genes was increased 2- to 4-fold in SOCS3^iEKO^ mice even in the absence of an endotoxin challenge. Conversely, a prior report showed that constitutive expression of type I IFNs sustains expression of JAK and STATs and predisposes cells to necroptotic death (90). In turn, necroptotic pathways may lead to NETosis (91), another feature of this model. The similarities observed between our in vitro experiments in IL-6+R-treated HUVEC and SOCS3^iEKO^ mice strongly support the argument that the described phenotype corresponds to an activation of the observed IL-6 amplifier loop. However, SOCS3 can block other gp130-mediated signals, and thus it remains possible that other gp130 receptor family ligands are involved in this response; further studies (e.g., using IL-6 inhibiting antibodies) will be required to assess this possibility.

In summary, we demonstrate that endothelial SOCS3 promotes vascular homeostasis and survival in a murine model of endotoxemia. Loss of SOCS3 in the adult endothelium leads to severe vasculopathy and kidney failure under systemic inflammation that is associated with an increased type I IFN-like transcriptional program. Importantly, loss of a single allele induced an intermediate phenotype, suggesting that maintenance of high SOCS3 levels is an absolute requirement. In vitro experiments showed that endothelial SOCS3 is a highly unstable protein, due to ubiquitin dependent proteasomal degradation. Thus, inhibition of endothelial SOCS3 degradation is a promising therapeutic opportunity to alleviate the vascular dysfunction during shock, increasing survival rates and reducing the risk of long-term sequelae.

## Methods

### Materials

The commercial sources for critical reagents and their catalog numbers are listed in Supplemental Table 1. Supplemental Table 2 lists all antibodies used. Supplemental Table 3 provides a list of sequences for RT-qPCR primers.

### Mice

#### Genetics

Endothelial specific, tamoxifen inducible SOCS3 knockout mice were generated by breeding B6.Tg(Cdh5-cre/ERT2)1Rha (Cdh5-CreER^T2^ endothelial driver) mice (92) (a kind gift from Dr Kevin Pumiglia, Albany Medical College) with B6.Gt(ROSA)26Sor^tm9(CAG-tdTomato)Hze^ (Rosa26-tdTomato reporter) (93) and B6;129S4-Socs3^tm1Ayos^/J (SOCS3^fl/fl^ conditional knockout) (94) mice (The Jackson Laboratory). Mice were backcrossed to a full C57Bl6/J background by breeding to C57Bl6/J mice (The Jackson Laboratory) for at least 10 generations. Knockouts, heterozygous and control littermates were obtained by crossing Cre^+^; tdTomato^+^; SOCS3^fl/+^ mice. Genotypes and gender were confirmed by PCR genotyping (Supplemental Table 4). All mice received tamoxifen (2mg tamoxifen in 100ul via intra-peritoneal (IP) injection) at 6-9 weeks old for 5 consecutive days. Deletion of target gene was confirmed by post-tamoxifen tail digestion and PCR. All the experiments were conducted between 2 and 3 weeks after the end of tamoxifen treatment.

#### Housing

Mice were housed in specific pathogen free rooms with 12 h light/dark cycles and controlled temperature and humidity. Mice were kept in groups of five or less in Allentown cages with access to food and water ad libitum.

#### Endotoxemic model

Severe, acute inflammation was induced by a single IP injection of a bolus of 250 μg/250 μl LPS. Control mice were given 250ul of sterile saline via IP injection. A severity scoring system was used to assess the response to LPS based on recently described scoring systems for similar conditions (33–35) (Supplemental Table 5). Body weight and temperature were obtained prior to LPS or saline injection and immediately after scoring.

#### Experimental design

Mice from at least one litter were used in each experiment. Mice from a second litter were used in some experiments to balance the number of mice for each gender in each experimental group. No mice were excluded from the studies. Assignment to the saline or LPS groups were performed through randomization of mice within each genotype for every litter. All handling, measurements and scoring were performed blindly to treatment and genotype groups and based on mouse ID# (ear tags). At least three independent experiments (from three or more litters) were performed for each assay. Experimental groups were unmasked at the end of each experiment.

### Blood measurements

Blood for complete blood count (CBC) was attained via a 5 mm lancet applied to the submandibular vein. Blood was collected into an EDTA blood collection tube and run on a Heska Element HT5 instrument. The remaining blood was centrifuged (1200 g, 15’, RT). Plasma was removed and stored at −80 °C until further processing.

To collect blood for chemistry panel, the inferior vena cava was snipped after pentobarbital administration and blood was collected into capillary tubes connected to a lithium heparin coated blood collection tube. Approximately 100ul were loaded into an Element point of care (POC) card and run on a Heska Element POC Blood Gas and Electrolyte Analyzer. Circulating levels of IL-6 and TNF-α were measured from plasma by ELISA following manufacturer’s instructions and plates were read on a Molecular Devices Spectramax I3. Circulating albumin from plasma was measured using the bromocresol purple (BCP) method following the manufacturer’s instructions. In addition to the supplied standards, mouse serum albumin was used as reference.

### Glomerular Filtration Rate

Glomerular filtration rate (GFR) was measured as previously described (95). Briefly, one to two days prior to experiment, a small area of dorsal fur was removed using an electric shaver followed by application of depilatory cream. Approximately 14-15 hours post LPS or saline injection, mice were briefly anaesthetized (3.5% v/v isoflurane) and placed on a heated pad at 37 °C. A MediBeacon transdermal Mini GFR Monitor was attached to the depilated region using a double-sided adhesive patch and adhesive, medical tape. The unit gathered a baseline measurement for approximately 1-2 minutes prior to injection of FITC-Sinistrin (MediBeacon) (7mg/100 gbw) in sterile saline solution via retro-orbital injection. Upon recovery from anesthesia, each mouse was returned to its own cage without access to food and water. After 90 min the instrument was removed and GFR was calculated with MediBeacon’s proprietary software.

### Assessment of Vascular Leak

Approximately 14-15 hours post LPS or saline injection, mice were briefly anaesthetized (3.5% v/v isoflurane) and placed on a heated pad at 37 °C. Then, mice received 100 μl of saline containing 2 mg/ml of FITC-labeled 70 kDa dextran and 0.13 mg/ml Alexa Fluor 647-labeled 10 kDa dextran was injected via retro-orbital injection. Upon recovery from anesthesia, mice were returned to the cages. After 30 min, mice were injected with pentobarbital. After confirmation of lack of paw reflex, the chest cavity was opened, and mice were perfused for 3 min with 5 ml/min RT PBS. Livers, kidneys, lungs and brains were fixed in 10% NBF for 24-48 h. Fresh NBF was replaced 24 h after organ collection. Then, kidneys, livers and lungs were maintained at 4 °C for two days or until organs sunk in 15% sucrose and 30% sucrose (both in PBS with 0.1 g/L sodium azide) respectively before embedding and freezing in Tissue-Tek optimum cutting temperature (OCT). Frozen 50 um sections on charged coverslips were stored at −20C until needed. Brains were transferred directly to 30% sucrose, 0.1 g/L sodium azide in PBS at 4 °C until the organs sunk (approximately 48 h). Brains were embedded and frozen in OCT prior to sectioning. 100 um sections were stored in 30% sucrose, 1% polyvinyl-pyrrolidone (average MW 40 kDa), 30% ethylene glycol, 0.1g/L sodium azide in PBS at 4 °C until use. After mounting, images were taken at 20x and 63x using a Leica Thunder microscope and processed for computational clearing. 3D 63x stacks were processed with Imaris software (Bitplane). Vascular leak was observed as patches of green fluorescence outside the vessel lumens.

### Histology and Immunofluorescence

#### Histochemistry

Lungs, liver and kidney were collected and placed into 10% Neutral Buffered Formalin (NBF) and stored in 70% ethanol prior to embedding in paraffin. 5 μm sections were then stained with Harris Hematoxylin and Eosin Y (H&E) following manufacturer’s protocols. Slides were scanned at 40x on a Nanozoomer (Hammamatsu C10730-12).

#### Frozen tissue immunofluorescence

Harvested tissue was embedded in OCT and snap frozen in a dry ice and methanol slurry. Blocks were stored at −80 °C until processed. 5 μm sections were post-fixed in 4% PFA for 15 minutes. After 3 washes in PBS, slides were permeabilized in PBS + 0.1% Triton X-100 (PBS-TX) for 15 minutes. Then tissue was blocked in 5% FBS in PBS-TX for 1 hour. Slides were stained with primary antibodies in blocking buffer overnight at 4 °C or 2 hours at RT. Slides were then washed in PBS-TX and stained with Alexa Fluor-conjugated secondary antibodies and 0.5 μg/ml DAPI for 1 hour at RT. Slides were then washed in PBS and mounted with Fluoroshield. Images were taken at 20x and 63x using a Leica Thunder microscope and processed for computational clearing.

#### Retina whole mount

After euthanasia, eyes were placed in 4% PFA at 4 °C. After 24 hours, eyes were transferred to PBS and stored at 4 °C until the retinas were removed. After removal of the cornea and lens, the edge of the retina was gently detached from the sclera. Four small cuts were made to the retina to allow flattening. Then, retinas were permeabilized and blocked by incubating for 1 h at 37 °C with 1% bovine serum albumin (BSA), 5% FBS, 0.5% Triton X-100 in PBS with gentle shaking. Retinas were then incubated with primary antibodies in permeabilization/blocking buffer overnight at 37 °C with gentle shaking. Retinas were then washed (x3) in PBS and incubated with Alexa Fluor-conjugated secondary antibodies (if required) and 1ug/ml DAPI in PBS at 37 °C for 4 hours. Retinas were then washed (x3) in PBS for 20 minutes/wash, flattened on the slides and mounted with Fluoroshield. Images were taken at 20x and 63x using a Leica Thunder microscope and processed for computational clearing.

### Cell culture and treatment

Human umbilical cord endothelial cells (HUVEC) were isolated in-house according to established protocols (27, 96–98). Briefly, umbilical cords (20-30 cm) from scheduled Cesarean sections were stored at 4 °C in phosphate-buffered saline (PBS) containing a mixture of penicillin and streptomycin and used within 24 h. Samples were anonymized following the recommendations of the Albany Medical Center IRB. After quick rinses in 70% ethanol and sterile PBS, vein lumens were washed with PBS to remove remaining blood and clots and then incubated for 30 minutes at RT in sterile PBS containing 0.2% collagenase, pH 7.4 with gentle massaging. Released cells were collected in growth media containing phenol red-free EBM 2 media supplemented with EGM-2 Growth Medium 2 Supplement Mix, penicillin, streptomycin and amphotericin B. Cells were then centrifuged, resuspended in fresh growth media and plated in plastic culture flasks pre-coated with 0.1% gelatin. Upon reaching initial sub-confluence (4-7 days), cells were passaged three times per week in the absence of antibiotics. Identity and purity of the HUVEC isolations was confirmed each time by >99% positive immunostaining with endothelial cell markers (FITC-*Ulex europaeus* lectin, VE-cadherin) and >99.9% negative for α-smooth muscle actin. Cells were assayed between passages 3 and 8. Unless otherwise stated, for all assays, cells were plated at full confluency at a density of 8×10^4^ cells/cm^2^ on plates precoated for 30 min with 0.1% gelatin and incubated at least 48 h prior to the start of experiments.

HUVEC were treated with inhibitors to JAK activity (ruxolitinib), proteasome (MG132) or protein synthesis (cycloheximide) dissolved in DMSO at a maximum use concentration of 0.1%. A similar amount of DMSO was added to vehicle control wells. To induce IL-6 signaling, cells were treated with a combination of 200 ng/ml recombinant human IL-6 and 100 ng/ml sIL-6Rα (IL-6+R).

### Lentiviral delivery

We created lentiviral constructs to express SOCS3 with C-terminal Myc and DDK tags by subcloning the human SOCS3 CDS and turbo green fluorescent protein (tGFP) in a single mRNA using a P2A linker following Origene’s TrueORF cloning instructions. Briefly, SOCS3 cDNA was cut with EcoRI and XhoI and subcloned into pLenti-C-Myc-DDK-P2A-tGFP lentiviral gene expression vector. A lysine 6 mutation (K6Q-SOCS3) was obtained by PCR mutagenesis using PfuUltra II Fusion High-fidelity DNA Polymerase. All constructs were confirmed by Sanger sequencing (Genewiz).

Lentiviral particles were grown in HEK293FT cells by co-transfecting cells with the SOCS3-expressing lentiviral constructs or pLenti-C-Myc-DDK-P2A-tGFP (empty vector control) and pCMV-dR8.2 dvpr and pCMV-VSVG packaging plasmids (99). After 48 hours, media was concentrated using a 30 kDa cutoff filter, aliquoted and stored at −80 °C until use. Viral titer was determined by the proportion of infected cells with green fluorescence 72h post-infection.

### Measurement of monolayer permeability

Monolayer permeability was determined in real time by measuring changes in electrical resistance using Electrical Cell-Substrate Impedance Sensor (ECIS, Applied Biophysics). HUVEC were seeded onto 8 or 96-well electrode arrays pre-coated with 0.1% gelatin at confluence. Following treatments, the electrical impedance across the monolayer was measured at 1 V, 4000 Hz AC for 24-72 continuous hours every 5 minutes and then used to calculate electric resistance by the manufacturer’s software.

### RNA-Sequencing

Confluent HUVEC grown on 0.1% gelatin pre-coated 6-well plates were pre-treated for 30 min with 2 μM ruxolitinib or 0.1% DMSO control and then treated with IL-6+R or PBS for 3 hours in triplicate. RNA was isolated with Trizol following manufacturer’s instructions. RNA enrichment and next-gen sequencing was performed by Genewiz. Briefly, after RNA quality testing, rRNA was eliminated using a magnetic bead approach and enriched RNA was subject to fragmentation and random priming for cDNA synthesis. Following end repair, 5’ phosphorylation, and dA-tailing, adaptor ligation was performed using NEBNext adaptors. Fastq files were obtained by collecting ~40×10^6^ reads (2×150 PE) per sample on an Illumina HiSeq instrument. The bioinformatics analysis was performed by following a workflow (100) using Salmon V0.8.1 (101), R/Bioconductor 3.5 (102), tximport (103) and DESeq2 (104). The data discussed in this publication have been deposited in NCBI’s Gene Expression Omnibus (105) and are accessible through GEO Series accession number GSE163649 (https://www.ncbi.nlm.nih.gov/geo/query/acc.cgi?acc=GSE163649).

### Statistics

All statistical analysis and graphs were made in GraphPad Prism version 6 or in R. Analysis for RNA and protein expression levels were performed using one-way ANOVA and Dunnett post-hoc test or a two-way ANOVA and Holm-Sidak post-hoc tests comparing all samples vs a control (designated in each figure). Two group comparisons were made with either Student’s T test (for parametric data) or Mann-Whitney tests for non-parametric data. A two-tailed p value of less than 0.05 was considered significant. For Western blot experiments, normalized band intensity values (protein/actin) were used. ECIS data were analyzed by two-way ANOVA of repeated measurements and post-hoc analysis comparing main column (treatment) effects. All unsupervised clustering for heatmaps, principal component analyses and cross-correlation studies were performed in R following a procedure described in the Supplemental Methods.

### Study approval

Mice were housed at the Animal Research Facility (ARF) at Albany Medical Center, a facility that has been accredited by AAALAC and licensed by the USDA and N.Y.S. Department of Public Health, Division of Laboratories and Research. All animal work was performed according to Institutional Animal Care and Use Committee (IACUC)-approved protocols.

## Author contributions

N.M. and A.P.A. conceived and designed research; N.M., R.B.R, S.L, K.L, L.T., S.S. and A.P.A. performed experiments; N.M., R.B.R, S.L, K.L, L.T., S.S., G.F., A.J., P.A.V. and A.P.A. analyzed data; N.M. and A.P.A. interpreted results of experiments; N.M., R.B.R., S.S. and A.P.A prepared figures; N.M. drafted manuscript; N.M. and A.P.A. edited and revised manuscript with input from all authors; N.M., R.B.R, S.L, K.L, L.T., S.S., G.F., A.J., P.A.V. and A.P.A. approved final version of manuscript.

## Acknowledgements

This project was supported by the National Institute of General Medical Sciences Grant R01GM124133 and an American Heart Association Transformational Project Award 18TPA34170561 to A.P.A. and National Heart, Lung and Blood Institute Grants R01HL141127 and R01HL153019 to G.F. and K01-HL130704 to A. J. We are grateful to Joseph Balnis for his help in measuring O_2_ saturation levels.

