## Supplemental data for "Endothelial SOCS3 maintains homeostasis and promotes survival in endotoxemic mice"

### Supplemental methods

#### Unbiased clustering and principal component analysis of mouse gene expression data

Graphs from mouse data for unsupervised clustering, principal component analysis and cross-correlation heatmaps with missing data were generated in R V4.0.2 following the workflow described below. The complete datasets (“All data.csv” and “Log2fold_data.csv” are provided on Supplemental Tables 6 and 7, respectively.

```
library(missMDA)
library(VIM)
library(FactoMineR)
library(factoextra)
library (pheatmap)

#load the cormat file to get the correlation matrix function
source("http://www.sthda.com/upload/rquery_cormat.r")

alldata <- read.csv ("All data.csv",stringsAsFactor=FALSE)
rownames (alldata) <- alldata[,1]
numdata <- alldata[,c(11,12,18:38,40:166)]
group <- alldata[, 5]

nb <- estim_ncpPCA(numdata,method.cv = "Kfold", verbose = FALSE)

res.comp <- imputePCA(numdata, ncp = nb$ncp)
res.comp$completeObs

imp <- cbind.data.frame(res.comp$completeObs,group)
res.pca <- PCA(imp, quali.sup = 151, ncp = nb$ncp, graph=FALSE)

pdf("plots.pdf")
plot(res.pca, hab=151, lab="quali") # colors by group
fviz_pca_ind(res.pca, habillage=151, legend.title = "Groups", palette="lancet", repel = TRUE,
geom = "point") # colors by group

# 30 most important variables colored by contribution
fviz_pca_var(res.pca, repel = TRUE, select.var = list(cos2 = 30), col.var = "contrib",
gradient.cols = c("#00AFBB", "#E7B800", "#FC4E07"))

contribs <- res.pca$var$contrib[,1:2]
contribs <- contribs[order(contribs[,1],decreasing=TRUE),]

rquery.cormat(numdata, graphType="heatmap")
corrmatrix<-rquery.cormat(numdata, type="flatten", graph=FALSE)
write.table (corrmatrix$r, file = "corrmatrix.txt", append = FALSE, quote = FALSE, sep = "\t",
eol = "\n", na = "NA", dec = ".", row.names = TRUE, col.names = TRUE)

LPS_numdata <- imp[grep("LPS", imp$group), ]
LPS_numdata <- LPS_numdata[,1:148]

rquery.cormat(LPS_numdata, graphType="heatmap")
LPS_corrmatrix<-rquery.cormat(LPS_numdata, type="flatten", graph=FALSE)
write.table (LPS_corrmatrix$r, file = "LPS_corrmatrix.txt", append = FALSE, quote = FALSE, sep =
"\t", eol = "\n", na = "NA", dec = ".", row.names = TRUE, col.names = TRUE)

logdata <- read.csv ("Log2fold_data.csv")
numlogdata <- logdata [,6:132]
rownames(numlogdata) <- logdata [,1]
head (numlogdata)

numdata_scale = scale(numlogdata)
pheatmap(numdata_scale,main = "pheatmap default", na_col="gray")

dev.off()
```

#### Gel Electrophoresis and Immunoblotting

Confluent HUVEC were lysed in Laemmli buffer containing the following protease and phosphatase inhibitors: cOmplete protease inhibitor mixture, PhosSTOP phosphatase inhibitor mixture, 0.1 M NaF, 0.1 mM phenyl arsine oxide, 10 mM pyrophosphate and 0.1 mM pervanadate. After boiling, a total of 15 µl of cell lysate per lane was loaded on standard SDS-PAGE gels and transferred to nitrocellulose membranes. Immunoblots were performed by blocking the membranes with 5% nonfat dry milk or 5% BSA in PBS/Tween and incubating overnight at 4°C with respective primary antibodies. Secondary HRP-conjugated anti-mouse, anti-rabbit, or anti-goat antibodies were incubated for 1 h at room temperature. Membranes were developed via chemiluminescence detected with either a LAS-3000 (Fujifilm) or a Chemidoc MP (Bio-Rad) imaging systems.

#### RNA isolation and RT-qPCR

RNA isolation from whole organs: Approximately 30-50mg of a target organ was removed and placed into an RNAse free 2ml tube containing zirconia/silicone beads and 1ml Trizol reagent. Tissue was homogenized using a Mini-Bead beater-16 (BioSpec) for 1 minute. Homogenate was transferred to a new RNAse free tube and the RNA isolation from Trizol was continued as per manufacturer's instructions.

RNA isolation from HUVEC: Confluent monolayers grown in multi-well plates were lysed with Trizol reagent and total RNA was isolated following manufacturer's instructions.

RT-qPCR: 400 ng of total RNA were used to prepare cDNA using Primescript RT Master Mix at 42°C following manufacturer's instructions. cDNA was diluted 10-fold in nuclease-free water, and then, 2µL of cDNA were used per PCR reaction. qPCR was performed in a StepOnePlus (Applied Biosystems) instrument using SYBR green-based iTaq supermix and 2 pmol primers (Thermo Fisher). Fold induction was calculated using the ΔΔCt method using GAPDH as housekeeping gene (HUVEC) or HPRT (mouse RNA).

#### Lung histological scoring

5 µm sections from FFPE lungs were processed for standard H&E and scanned with a Hammamatsu C10730-12 Nanozoomer at 40x. Scans identified only by mouse ID number were scored by a single investigator (A.J.) that remained masked to experimental groups (both treatment and genotype) until all slides were scored. Inflammatory infiltration, intra alveolar and interstitial exudate accumulation, tissue necrosis, septal and alveolar wall thickening, microhemorrhage and thrombosis were all scored in a scale of 1-4 (1 being the least and 4 being the most abnormal); in four different areas identified first with lower and then higher magnification. These magnifications were variably selected by the operator as needed to make the observation based in the dynamic view provided by the software.

#### Immunohistochemistry

5 µm FFPE sections from lung, liver of kidney were processed for colorimetric histochemistry. Briefly, slides were deparaffinated and rehydrated in sequential steps in xylene and ethanol solutions (100%/ 95%/70%). Endogenous peroxidase activity was blocked with 0.5% $H_2O_2$/MeOH for 10 minutes at room temperature (RT) and antigen retrieval (EZ-Retriever System, Biogenex) was done for 15 minutes at 98 C in 10 mM citrate buffer, pH 6 (for F4/80) or

in 1mM EDTA pH 8 (for MPO). Samples were blocked with FBS for 1 hour at room temperature. Primary antibodies were incubated at a dilution of 1:200 (F4/80) or 1:500 (MPO) in PBS overnight at 4 C. Biotinylated anti-rabbit secondary antibodies 1:500 in PBS were incubated for 1 hour at RT. Then, samples were incubated with avidin/biotin peroxidase (Elite ABC-HRP, Vector) in the dark for 30 minutes at RT and signal was detected with 3,3'-diaminobenzidine (Immpact DAB, Vector) and counterstained with hematoxylin prior to dehydration and mounting with VectaMount. Whole slides were scanned with a Hammamatsu C10730-12 Nanozoomer at 40x.

**Flow cytometry**

Mice were euthanized with an overdose of pentobarbital. The chest cavity was opened, and 100 µl of blood were collected from a nicked aorta into EDTA-containing vials. Lungs were excised, washed several times with PBS, and minced finely with scissors. Then, the minced lungs were transferred to a 50ml conical tube containing 6 ml of pre-warmed collagenase type I/dispase II/DNAse I mix (CDD) and incubated at 37 ºC with gentle agitation for 1 hour. Single cell suspensions were obtained after the suspensions were passaged multiple times through 20-gauge cannulas and filtered through a 70 µm mesh. Cells were then spun at 400 g for 8 minutes at 4 ºC and resuspended in 3 ml of cold PBS containing 0.1% BSA.

In parallel, bone marrow cells were collected by flushing the right femur with 10ml PBS onto a 70 µm cell strainer and then centrifuged at 6000 rpm for 5 min 4ºC. Red blood cells from blood and bone marrow cells were lysed using RBC lysis buffer according to manufacturer's instructions. Live and dead cells from lungs, blood and bone marrow were distinguished using Pacific blue Annexin V Apoptosis Detection kit with 7-AAD. Lung single cell suspensions (300 µl / sample) were then stained with ly6G-APC-Cy7 to label neutrophils. Cells were then subjected to flow cytometry analysis using a BD FACS symphony. 30,000 events were acquired for each sample and the data was analyzed using FlowJo v10.7.1. Single cells were gated using FSC-H vs FSC-A, dead cells were excluded based on 7AAD-Annexin-V and only live cells were gated for further analysis. Results are expressed as percentage of total cells.

**Isolation of RNA from spleen CD45+ cells**

Control, SOCS3iEKO or heterozygous mice were treated with tamoxifen to induce Cre activation as described in the main text. Two weeks after, mice were sacrificed to collect spleens. Organs were then washed in PBS. One third of each spleen was transferred into a tube with Trizol to have RNA of total organ isolated later following the Trizol protocol. The remaining portion of each spleen was pressed with a plunger from a 10 cc syringe. Released cells and tissue remains were collected and pushed through 70 µm cell strainers to obtain single cells isolates. Cells were centrifuged at 400g for 8 min at 4 ºC. The cell pellet was then resuspended in 1.5 ml ice-cold PBS+0.1%BSA. The cell suspension was incubated with anti-CD45 conjugated Dynabeads M280 streptavidin beads (3 µg ab,300 µg beads/sample) for 10 minutes at RT under gentle rotation. CD45+ cells were separated using a magnetic separator (Dynal MPC-S) and washed six times with PBS+0.1%BSA. Cells were placed on magnet for 3 minutes, the supernatant was removed and cells were lysed in Trizol reagent. RNA was extracted following a modified protocol. Briefly, 0.1 ml chloroform was added to 0.5 ml of Trizol per sample. After a 2 min incubation, tubes were centrifuged for 15 min at 12,000 g at 4 ºC. The aqueous solution was then recovered, mixed 1:1 with 70% EtOH, and applied to a silica column (RNeasy Plus Micro kit, Qiagen). Then, the columns were washed, and the RNA was eluted following the RNeasy Plus Micro standard protocol. Efficient gene deletion was confirmed by tail snip genotyping as described in the main text.

#### ELISA

HUVEC cells were cultured in six-well plates at a density of $7.2\times10^5$ cells/well. 14 hours before treatment, the media was reduced to 800 μL/well. Cells were then treated with IL6+R or vehicle control (PBS). Culture media was collected after 6 hours and centrifuged at 18,000 g for 10 min at 4 ˚C. Then, supernatant was aliquoted and stored at -80˚C until analysis. The levels of IFNα, IFNβ and IFNγ in culture media were measured using ELISA kits according to the manufacturer's instructions.

#### Gene knockdown

HUVEC were transfected by plating cells onto multi-well plates containing individual small interference RNA (siRNA) against SOCS3 or a non-targeting control pool complexed with lipofectamine RNAiMAX transfection reagent in suspension and seeded at $10^5$ cells/cm$^2$. Knockdown efficiency was determined by Western blot analysis.

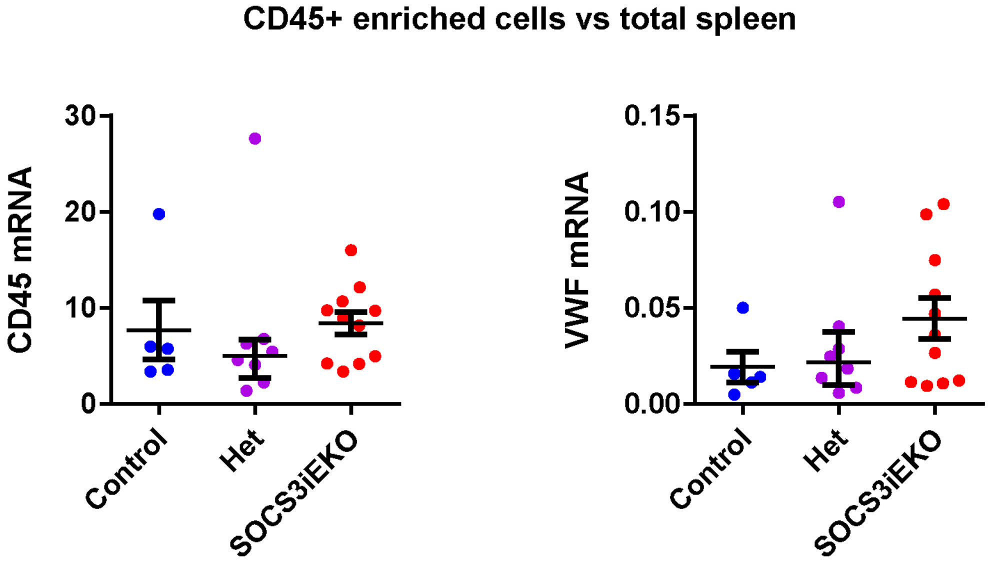

**Supplemental Figure 1. Enrichment of CD45 cells from spleen.** Spleens from control, heterozygous or SOCS3iEKO mice (corresponding to those shown in Figure 1C) were collected 14 days after tamoxifen treatment. CD45 cells were isolated using magnetic beads prior to RNA extraction and RT-qPCR. Enrichment was confirmed by a 5-10-fold increase in CD45 expression and >10-fold decrease in VWF expression compared to total RNA from the same spleens. NS: not statistically significant (Kruskal-Wallis). Data combined from three independent experiments.

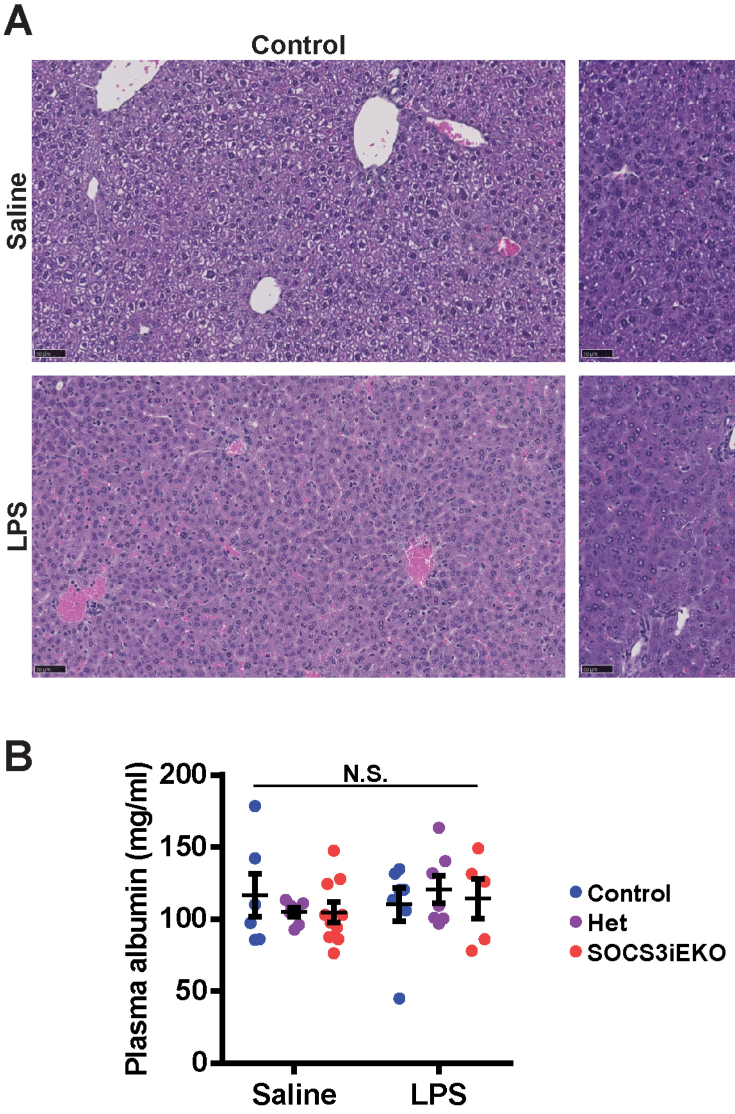

**Supplemental Figure 2. Mild liver dysfunction in endotoxemic SOCS3$^{iEKO}$**. A, H&E staining showing a reduction in glycogen content but no other overt histological features in endotoxemic SOCS3$^{iEKO}$ mice. B, Plasma albumin levels remained similar among all experimental groups (two-way ANOVA). Data combined from at least three independent experiments.

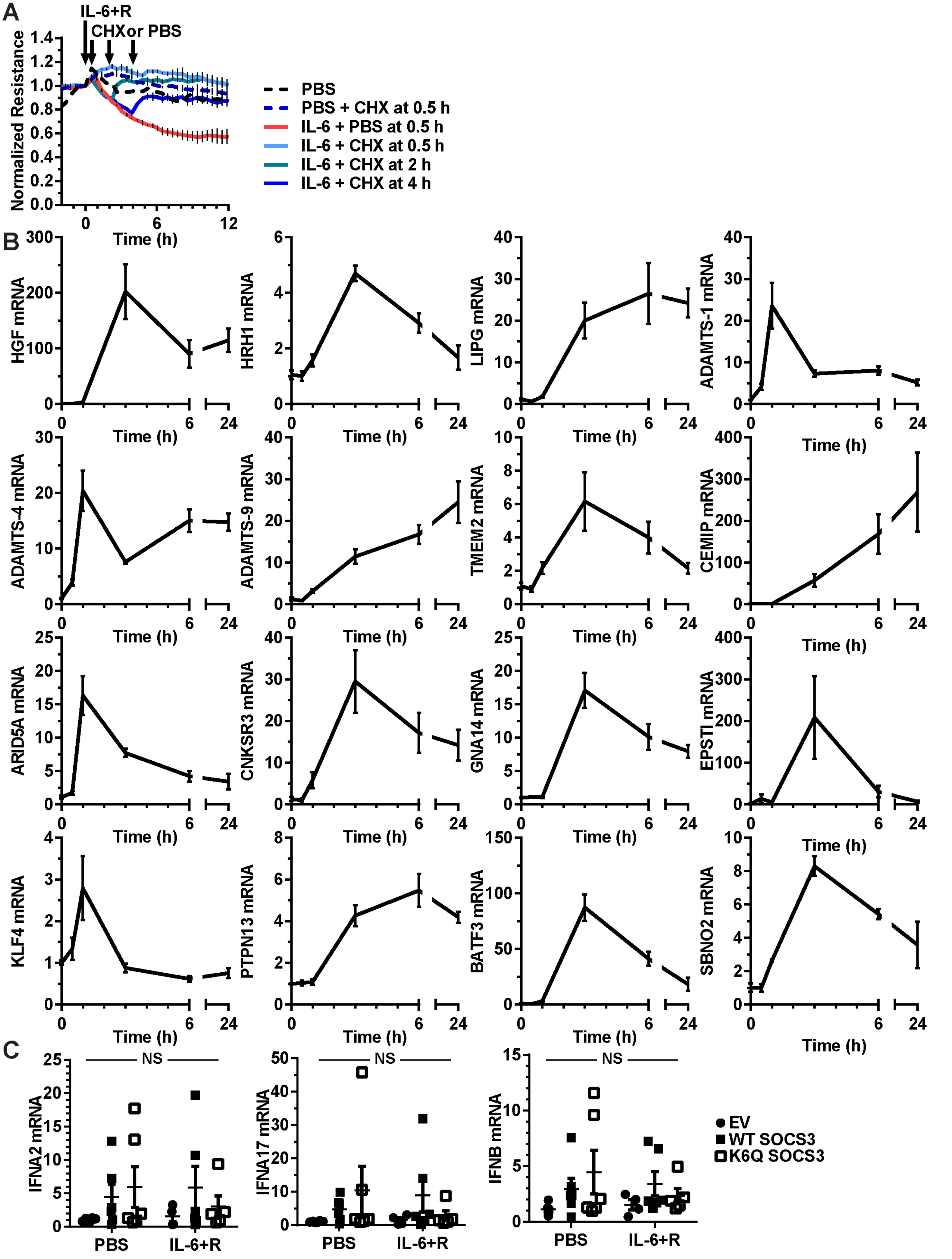

**Supplemental Figure 3. Gene expression changes in HUVEC treated with IL-6+R**. A, IL-6+R-mediated loss of barrier function requires continuous protein synthesis. B, RT-qPCR of IL-6+R-treated HUVEC. Data was obtained as described in Figure 7. Data combined from three independent experiments. C, RT-qPCR of cells treated with or without IL-6+R after infection with lentivirus to overexpress WT SOCS3 or K6Q-SOCS3. An empty vector lentivirus was used as control. Two-way ANOVA. Combined data from three independent experiments performed in duplicate each.

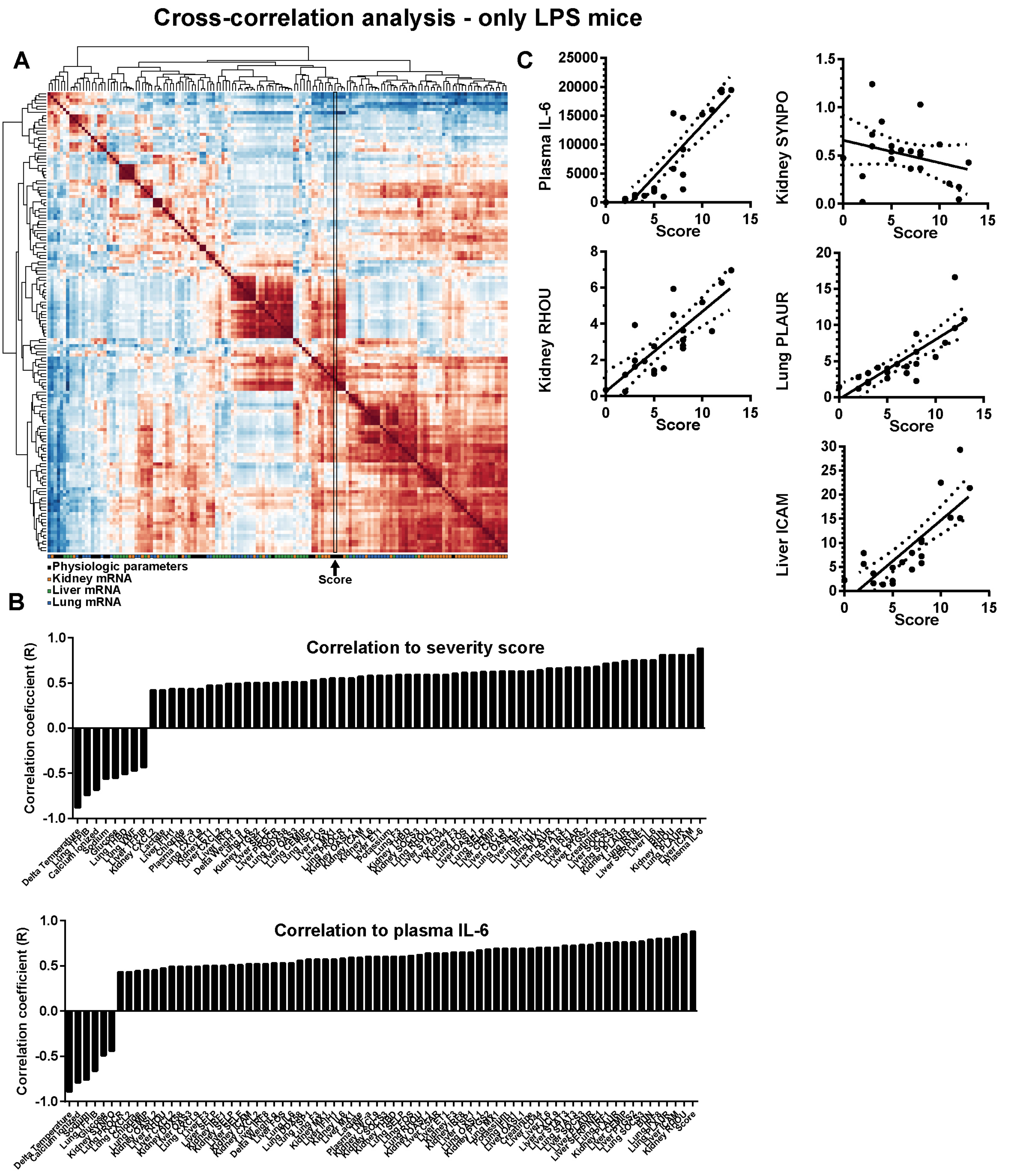

**Supplemental Figure 4. Multiple significant parameters correlate with the severity score and plasma IL-6 levels**. A, Cross-correlation analysis for all parameters measured in LPS-treated mice (full data available as Supplemental Table 6). B, Graph showing all significantly correlated parameters to the severity score or plasma IL-6. C, Examples of the correlation of expression of specific genes (expressed as a base 2 logarithm of fold changes) with the severity score for each mouse. Data combined from three independent experiments.

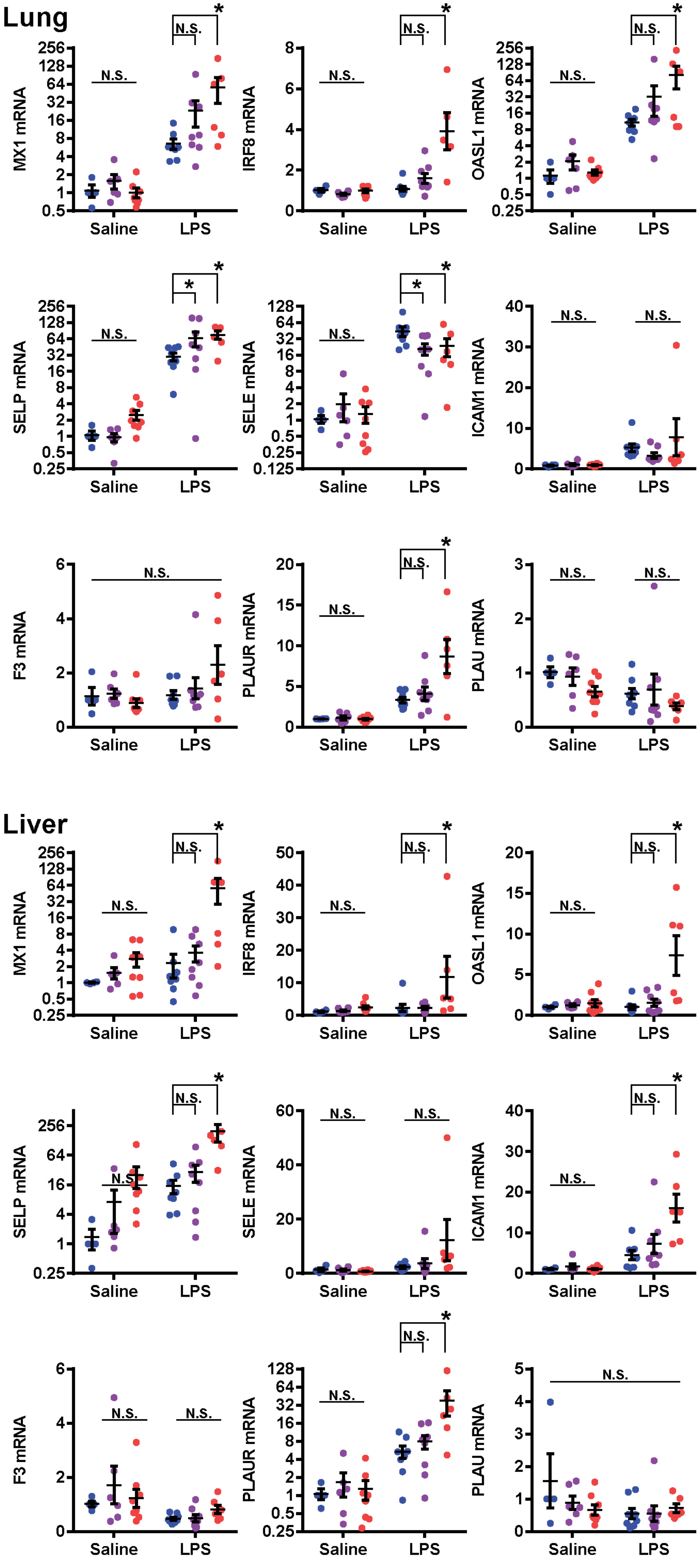

**Supplemental Figure 5. Type I IFN-like, adhesive and prothrombotic gene expression in endotoxemic SOCS3$^{iEKO}$ mice**. RT-qPCR analysis of whole lung (A) or liver (B) RNA levels. Two-way ANOVA and Holm-Sidak post-hoc tests comparing het and SOCS3$^{iEKO}$ mice to control within saline or LPS-treated groups. Asterisks denote $p < 0.05$. Data combined from at least three independent experiments.

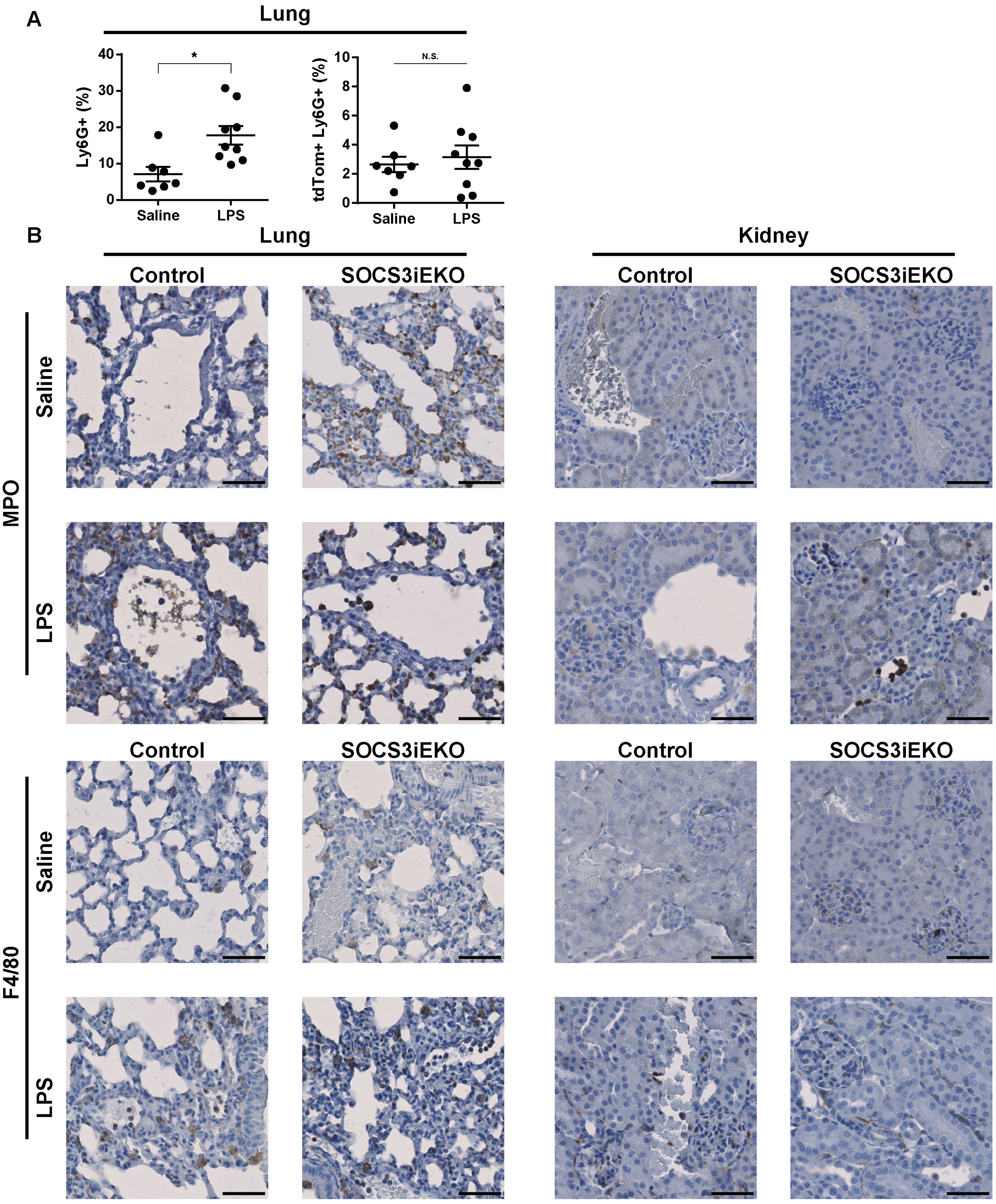

**Supplemental Figure 6. Neutrophil and monocyte infiltration in endotoxemic organs**. A, flow cytometry analysis of Ly6G+ cells in lungs. Asterisks denote $p < 0.05$. Data from two independent experiments (n=7 saline- and n=9 LPS-treated mice). B, Immunohistochemical staining of FFPE sections for myeloperoxidase (MPO, a neutrophil marker) and F4/80 (a monocyte marker) in lungs and kidneys. Bars, 50 µm. Images are representative from three independent experiments.

**Supplemental Table 1**

| M&M section | Reagent or disposable | Vendor | Catalog Number |
|---|---|---|---|
| **Mice** | Lipopolysaccharides from *Escherichia coli* O111:B4 | Millipore Sigma | L4391 |
| | B6.Gt(ROSA)26Sor$^{tm9(CAG-tdTomato)Hze}$ | The Jackson Laboratory | 07909 |
| | B6;129S4-Socs3$^{tm1Ayos}$/J | The Jackson Laboratory | 010944 |
| | Goldenrod 5 mm lancet | Medipoint | 5mm |
| | EDTA Vacutainer tube | BD | 367836 |
| | Lithium Heparin Capillary Blood Collection Tube | Sarstedt | 20.1292.100 |
| **Blood measurements** | Element POC Test Card with BUN | Heska | 5424 |
| | IL-6 ELISA | Proteintech | KE10007 |
| | TNF-α ELISA | Proteintech | KE10002 |
| | BCP Albumin Assay Kit | Abcam | AB272526 |
| | Mouse serum albumin | Millipore Sigma | A3139 |
| **Glomerular Filtration Rate** | Transdermal Mini GFR Monitor | MediBeacon | TDM-MD004 |
| | FITC-Sinistrin | MediBeacon | FTC-FS001 |
| **Assessment of Vascular Leak** | FITC-labeled 70 kDa dextran | Invitrogen | D1822 |
| | Alexa Fluor 647-labeled 10 kDa dextran | Invitrogen | D22914 |
| | sucrose | VWR | 97061-426 |
| | Polyvinyl-pyrrolidone (avg MW 40 kDa) | Millipore Sigma | PVP-40 |
| | Ethylene Glycol | VWR | 97061-964 |
| | Tissue-Tek optimum cutting temperature | Sakura Finetek | 4583 |
| | DAPI | Invitrogen | D3571 |
| | Fluoroshield with 1,4-Diazabicyclo [2.2.2] octane | Millipore Sigma | F937 |
| **Histology and Immunofluorescence** | Harris Hematoxylin | Millipore Sigma | HHS16 |
| | Eosin Y | Richard Allen | 7111 |
| | Periodic Acid-Schiff (PAS) Kit | Millipore Sigma | 395B |
| | Gill No. 3 Hematoxylin | Millipore Sigma | GHS332 |
| | Triton X-100 | Millipore Sigma | T8787 |
| | Fetal bovine serum | Millipore Sigma | 12306C |
| | BOVINE SERUM ALBUMIN - Fraction V (Immunoglobulin and Protease Free) | Rockland | BSA-50 |
| **Cell culture** | collagenase | Worthington | LS004196 |
| | phenol red-free EBM 2 media | PromoCell | C-22211 |
| | EGM-2 Growth Medium 2 Supplement Mix | PromoCell | C-39216 |
| | Trypsin-EDTA Solution 1X | Millipore Sigma | 54917C-100ML |
| | Penicillin-Streptomycin Solution, 100x | Corning | 30-002-CI-PK |

| | | | |
|---|---|---|---|
| | Antibiotic Antimycotic Solution (100×), Stabilized | Millipore Sigma | A5955-20ML |
| | FITC-Ulex europaeus lectin | Millipore Sigma | L9006 |
| | gelatin | Millipore Sigma | ES-006-B |
| | Ruxolitinib | Selleck Chem | S1378 |
| | MG132 | Calbiochem | 474790 |
| | Cycloheximide | Acros Organics | 357420010 |
| | DMSO | Millipore Sigma | 472301 |
| | Recombinant human IL-6 | R&D Systems | 206-IL-050 |
| | Recombinant human sIL-6Rα | R&D Systems | 227-SR-025/CF |
| **Lentiviral delivery** | SOCS3 (NM_003955) Human Tagged ORF Clone | Origene | RC209305 |
| | pLenti-C-Myc-DDK-P2A-tGFP Lentiviral Gene Expression Vector | Origene | PS100088 |
| | PfuUltra II Fusion High-fidelity DNA Polymerase | Agilent | 600670 |
| | HEK293FT | Invitrogen | R70007 |
| | pCMV-dR8.2 dvpr | Addgene | 8455 |
| | pCMV-VSVG | Addgene | 8454 |
| | Vivacell 100, 30,000 MWCO PES, 10pc | Sartorius | VC1022 |
| **Measurement of monolayer permeability** | 8-well electrode array | Applied Biophysics | 8W10E PET |
| | 96-well electrode array | Applied Biophysics | 96W10idf PET |
| **Gene knockdown** | ON-TARGETplus set of 4 SOCS3 siRNA | Dharmacon | LQ-004299-00-0005 |
| | ON-TARGETplus non-targeting control pool | Dharmacon | D-001810-10-20 |
| | Opti-MEM | Thermo Scientific | 31985070 |
| | lipofectamine RNAiMAX transfection reagent | Invitrogen | 13778150 |
| **Gel Electrophoresis and Immunoblotting** | complete protease inhibitor mixture | Roche Applied Science | 11697498001 |
| | PhosSTOP phosphatase inhibitor mixture | Roche Applied Science | 4906845001 |
| | Sodium fluoride | Millipore Sigma | S-1504 |
| | phenyl arsine oxide | Millipore Sigma | P-3075 |
| | Sodium pyrophosphate decahydrate | Millipore Sigma | 221368-100G |
| | Sodium orthovanadate | Millipore Sigma | S6508 |
| | Transblot Turbo RTA Mini Nitrocellulose Transfer kit | Bio-Rad | 1704270 |
| | Clarity Western ECL Substrate | Bio-Rad | 1705061 |
| | Clarity Max Western ECL Substrate | Bio-Rad | 1705062 |
| | SuperSignal West Femto Maximum Sensitivity Substrate | Pierce | 34094 |

| | | | |
|---|---|---|---|
| **RNA isolation and RT-qPCR** | 1.0 mm dia zirconia/silica beads | Biospec | 11079110z |
| | TRIzol reagent | Invitrogen | 15596018 |
| | PrimeScript RT Master Mix | Takara Bio | RR036B |
| | iTaq Universal SYBR Green Supermix | Bio-Rad | 1725125 |
| **Flow cytometry** | Collagenase type I | Worthington | LS004197 |
| | Dispase II | Roche Applied Science | 10888700 |
| | DNAse I | Roche Applied Science | 10104159001 |
| | EDTA Capillary Blood Collection Tube | Sarstedt | 20.1288.100 |
| | Pacific blue Annexin V Apoptosis Detection kit with 7-AAD | Biolegend | 640926 |
| | RBC lysis buffer | Roche Applied Science | 11814389001 |
| **ELISA** | IFNα kit | Thermo Scientific | BMS216 |
| | IFNβ kit | PBL Assay Science | 41410 |
| | IFNγ kit | Thermo Scientific | EHIFNG |
| **Immunohistochemistry** | VECTASTAIN Elite ABC-HRP Kit | Vector Labs | PK-6100 |
| | ImmPACT® DAB Substrate, Peroxidase | Vector Labs | SK-4105 |
| | VectaMount Permanent Mounting Medium | Vector Labs | H-5000 |
| **Isolation of RNA from spleen CD45+ cells** | Dynabeads M-280 Streptavidin | Invitrogen | 11205D |
| | RNeasy Plus Micro Kit | Qiagen | 74034 |
| | Falcon 70 µm Cell Strainer | Corning | 352350 |

**Supplemental Table 2**
**List of antibodies**

| Target | Species | Clone or isotype | Conjugate | Vendor | Catalog No | RRID |
|---|---|---|---|---|---|---|
| **STAT3** | rabbit | clone D1A5 | | Cell Signaling Technology | 8768S | AB_2722529 |
| **pY705-STAT3** | rabbit | clone D3A7 | | Cell Signaling Technology | 9145L | AB_2491009 |
| **SOCS3** | rabbit | polyclonal IgG | | Abcam | AB16030 | AB_443287 |
| **E-selectin** | rabbit | polyclonal IgG | | Abcam | AB18981 | AB_470289 |
| **P-selectin** | goat | polyclonal IgG | | R&D Systems | AF737 | AB_2285644 |
| **CD45** | mouse | clone 30-F11 | FITC | BioLegend | 103107 | AB_312972 |
| **CD45** | mouse | clone 30-F11 | Biotin | BioLegend | 103104 | AB_312969 |
| **Histone H3 (citrulline R2 + R8 + R17)** | rabbit | polyclonal IgG | | Abcam | AB5103 | AB_304752 |
| **DDK** | mouse | clone OTI4C5 | | Origene | TA50011100 | AB_2622345 |
| **Turbo GFP** | rabbit | polyclonal IgG | | Fujifilm Wako Chemicals | AB513 | |
| **β-actin** | mouse | clone AC-15 | | Millipore Sigma | A5441 | AB_476744 |
| **α-smooth muscle actin** | mouse | clone 1A4 | | Millipore Sigma | A2547 | AB_476701 |
| **VE-cadherin** | goat | polyclonal | | R&D Systems | AF938 | AB_355726 |
| **Myeloperoxidase** | rabbit | clone EPR20257 | | Abcam | ab208670 | AB_2864724 |
| **F4/80** | rabbit | clone D2S9R | | Cell Signaling | 70076S | AB_2799771 |
| **Ly6G** | rat | clone 1A8 | APC/Cy7 | BioLegend | 127624 | AB_10640819 |
| **mouse IgG** | goat | polyclonal | HRP | Jackson Immunoresearch | 115-035-062 | AB_2338504 |
| **rabbit IgG** | goat | polyclonal | HRP | Jackson Immunoresearch | 111-035-003 | AB_2313567 |
| **goat IgG** | bovine | polyclonal | HRP | Jackson Immunoresearch | 805-035-180 | AB_2340874 |
| **mouse IgG** | donkey | polyclonal | Alexa Fluor 647 | Invitrogen | A-31571 | AB_162542 |
| **rabbit IgG** | donkey | polyclonal | Alexa Fluor 647 | Invitrogen | A-31573 | AB_2536183 |
| **goat IgG** | donkey | polyclonal | Alexa Fluor 647 | Invitrogen | A-21447 | AB_2535864 |
| **rabbit IgG** | goat | polyclonal | Biotin | Vector Labs | BA-1000 | AB_2313606 |

**Supplemental Table 3**
**Mouse RT-qPCR primers (SYBR green)**

| Gene symbol | Forward | Reverse |
|---|---|---|
| **HPRT** | TGGCCCTCTGTGTGCTCAA | TGATCATTACAGTAGCTCTTCAGTCTGA |
| **C5AR** | CCATTAGTGCCGACCGTTTCCT | CACGAAGGATGGAATGGTGAGG |
| **CD44** | CGGAACCACAGCCTCCTTTCAA | TGCCATCCGTTCTGAAACCACG |
| **CEMIP** | CTCTACACAGGTCAAGGTGGCA | CCATCACCACAATGTTCCGAGTC |
| **CLU** | GATGATCCACCAGGCTCAACAG | ACACAGTGCGGTCATCTTCACC |
| **CXCL2** | CATCCAGAGCTTGAGTGTGACG | GGCTTCAGGGTCAAGGCAAACT |
| **CXCL9** | CCTAGTGATAAGGAATGCACGATG | CTAGGCAGGTTTGATCTCCGTTC |
| **DDX58** | AGCCAAGGATGTCTCCGAGGAA | ACACTGAGCACGCTTTGTGGAC |
| **ET1** | CTACTTCTGCCACCTGGACATC | CGCACTGACATCTAACTGCCTG |
| **F3** | GCACCGAGCAATGGAAGAGTTTC | CTTTCTGTCCCGCTCGGTTCTT |
| **FOS** | GGGAATGGTGAAGACCGTGTCA | GCAGCCATCTTATTCCGTTCCC |
| **HAVCR1** | CTGGAATGGCACTGTGACATCC | GCAGATGCCAACATAGAAGCCC |
| **ICAM** | AAACCAGACCCTGGAACTGCAC | GCCTGGCATTTCAGAGTCTGCT |
| **HNF1** | AGAGACCTTGGTGGAGGAGTGT | GGCAAACCAGTTGTAGACACGC |
| **IFIH1** | TGCGGAAGTTGGAGTCAAAGCG | CACCGTCGTAGCGATAAGCAGA |
| **IL18** | GACAGCCTGTGTTCGAGGATATG | TGTTCTTACAGGAGAGGGTAGAC |
| **IL6** | TACCACTTCACAAGTCGGAGGC | CTGCAAGTGCATCATCGTTGTTC |
| **IRF1** | TCCAAGTCCAGCCGAGACACTA | ACTGCTGTGGTCATCAGGTAGG |
| **IRF7** | CCTCTGCTTTCTAGTGATGCCG | CGTAAACACGGTCTTGCTCCTG |
| **IRF8** | AGGTCTTCGACACCAGCCAGTT | GCACGAGAATGAGTTTGGAGCG |
| **KLF2** | CGCCGCCACACATACTTG | AACTTCCAGCCGCATCCTT |
| **LCN2** | ATGTCACCTCCATCCTGGTCAG | GCCACTTGCACATTGTAGCTCTG |
| **MX1** | TGGACATTGCTACCACAGAGGC | TTGCCTTCAGCACCTCTGTCCA |
| **OAS2** | CACCAAAGTCCTGAAGACCGTC | AGAGTCGTAACTCTCCAGCGAG |
| **OAS3** | TTCTCTGCCAGCTTCGGAAAGC | CTCTGAAGGCAGACTTGTGACC |
| **OASL1** | TGAAGAGCCTCCTTCGGTTGGT | TCCAGCCTGAAGTTGGCATCCT |
| **OASL2** | CCAAAACGAGGTCGTCAGGAAC | AGCCACCTGTTCCCATCCCTTT |
| **PLAT** | GTTACACAGCGTGGAGGACCAA | CACGTCAGCTTTCGGTCCTTCA |
| **PLAU** | AGAAGCGACCCTGGTGCTATGT | CCACACTGGAAGCCTTGTTGGT |
| **PLAUR** | AGGACTACCGTGCTTCGGGAAT | ACACGGTCTCTGTCAGGCTGAT |
| **PROCR** | CCTCCAAAGCAGCCAACTTCAC | GTGTAAGAGCGACCTGTTTGGC |
| **PTGS2** | GCGACATACTCAAGCAGGAGCA | AGTGGTAACCGCTCAGGTGTTG |
| **RHOU** | TGGTCAGCTACACCACTAACGG | TGCAGTGTCACAGAGCTGGAGT |
| **SELE** | GGACACCACAAATCCCAGTCTG | TCGCAGGAGAACTCACAACTGG |
| **SELP** | AAGATGCCTGGCTACTGGACAC | CAAGAGGCTGAACGCAGGTCAT |
| **SERPINE1** | CCTCTTCCACAAGTCTGATGGC | GCAGTTCCACAACGTCATACTCG |
| **SOCS3** | GGACCAAGAACCTACGCATCCA | CACCAGCTTGAGTACACAGTCG |
| **STAT2** | GAACCAACTCTCCATTGCCTGG | CGTAAGAGGAGAACTGCCAGCT |
| **STAT3** | AGGAGTCTAACAACGGCAGCCT | GTGGTACACCTCAGTCTCGAAG |
| **SYNPO** | TCCTTCTCCACCCGGAATGCTG | AGCCGTCCAGGCTGCTAGGAG |
| **TFPIA** | CTGTGAGAATCCAGTCCACTCC | GCCACGATAATCCCGACG |
| **TFPIB** | CTGTGAGAATCCAGTCCACTCC | TCTTCTTTTGTAACCCGACGC |
| **TFPIC** | CTGTGAGAATCCAGTCCACTCC | TACCAAGGCAGCCCGAC |
| **THBD** | GGAGAATGGTGGCTGTGAGTAC | GCACGATTGAACCACAGGTCTTG |

| | | |
|---|---|---|
| **TSP1** | TGGCCAGCGTTGCCA | TCTGCAGCACCCCCTGAA |
| **VEGFA** | CTGCTGTAACGATGAAGCCCTG | GCTGTAGGAAGCTCATCTCTCC |
| **VWF** | AACAGACGATGGTGGACTCAGC | CGATGGACTCACAGGAGCAAGT |
| **PTPRC** | CTTCAGTGGTCCCATTGTGGTG | TCAGACACCTCTGTCGCCTTAG |

**Human RT-qPCR primers (SYBR green)**

| Gene symbol | Forward | Reverse |
|---|---|---|
| **GAPDH** | GTCTCCTCTGACTTCAACAGCG | ACCACCCTGTTGCTGTAGCCAA |
| **SOCS3** | CATCTCTGTCGGAAGACCGTCA | GCATCGTACTGGTCCAGGAACT |
| **IL6** | AGACAGCCACTCACCTCTTCAG | TTCTGCCAGTGCCTCTTTGCTG |
| **CXCL2** | GGCAGAAAGCTTGTCTCAACCC | CTCCTTCAGGAACAGCCACCAA |
| **RHOU** | ACTGCCTTCGACAACTTCTCCG | GAGCAGGAAGATGTCTGTGTTGG |
| **PTGS2** | CGGTGAAACTCTGGCTAGACAG | GCAAACCGTAGATGCTCAGGGA |
| **IFNA2** | TGGGCTGTGATCTGCCTCAAAC | CAGCCTTTTGGAACTGGTTGCC |
| **IFNA17** | GAAGACTCAAGCCATCTCTGTCC | TAGGAGGCTCTGTTCCCAAGCA |
| **IFNB** | CTTGGATTCCTACAAAGAAGCAGC | TCCTCCTTCTGGAACTGCTGCA |
| **MX1** | GGCTGTTTACCAGACTCCGACA | CACAAAGCCTGGCAGCTCTCTA |
| **IRF1** | GAGGAGGTGAAAGACCAGAGCA | TAGCATCTCGGCTGGACTTCGA |
| **IRF7** | CCACGCTATACCATCTACCTGG | GCTGCTATCCAGGGAAGACACA |
| **ADAMTS1** | GCGTCAATGCTTTCCAACCTGG | GGGATTCTGAGGCTTGTCCATC |
| **ADAMTS4** | TCACTGACTTCCTGGACAATGGC | GGTCAGCATCATAGTCCTTGCC |
| **ADAMTS9** | CCATTCAGAGGTGCAGTGAGTTC | ACCAGACCTGGCGGTGCTTATG |
| **TMEM2** | GGAATAGGACTGACCTTTGCCAG | TTCTGACCACCCTGAAAGCCGT |
| **CEMIP** | ACCGAGCACATTCCAACTACCG | GGCAGAGATGATTGAGAGGAACG |
| **ARID5A** | TGGCAAGCAGAACGGAATCCAG | CTTGTAGAGGCTGACCAGGAAG |
| **CNKSR3** | AAAACCTACGGTGGAAGCCACC | ATCCAGGATGGCTGACTTCTCC |
| **EPSTI** | ACTGAAACGGCAGCAGCAAGAG | TCCAACAGCCTCCAGATTGCTC |
| **GNA14** | AAGCAGCTCTGGCAAGATCCAG | GTAGGCACGAATGATGGTGTGG |
| **KLF4** | ACCTACACAAAGAGTTCCCATC | TGTGTTTACGGTAGTGCCTG |
| **PTPN13** | GGATGAAGCCACTTACTCCAGC | CTCCAGGCTTAGGAGGTGATGA |
| **BATF3** | ACCGAGTTGCTGCTCAGAGAAG | AGGTGCTTCAGCTCCTCTGTCA |
| **SBNO2** | GAGAACGATGGGCACCTCAACT | TGTCCCGCTTTCTCTTGGTGGA |

**Supplemental Table 4**
**Primers for genotyping**

| | | Male | Female |
|---|---|---|---|
| **SRY** | | **273 bp** | **-** |
| Fwd | TTG TCT AGA GAG CAT GGA GGG CCA TGT CAA | | |
| Rev | CCA CTC CTC TGT GAC ACT TTA GCC CTC CGA | | |

| | | WT | Flox |
|---|---|---|---|
| **SOCS3 flox** | | **52 bp** | **181 bp** |
| Fwd | CGG GCA GGG GAA GAG ACT GT | | |
| Rev | TCG ACT GTC CTC GGT CAC | | |

| | | WT | Tom |
|---|---|---|---|
| **Rosa26 locus** | | **297 bp** | **196 bp** |
| Wild type Fwd | AAG GGA GCT GCA GTG GAG TA | | |
| Wild type Rev | CCG AAA ATC TGT GGG AAG TC | | |
| Mutant Fwd | CTG TTC CTG TAC GGC ATG G | | |
| Mutant Rev | GGC ATT AAA GCA GCG TAT CC | | |

| | | Present | Deleted |
|---|---|---|---|
| **SOCS3 Deletion** | | **>1400 bp** | **424 bp** |
| Fwd | GCG GGC AGG GGA AGA GAC TGT CTG GGG TTG | | |
| Rev | CCT TTT CTC TTC CAT CCT TCC | | |

| | | WT | Cre |
|---|---|---|---|
| **Cre** | | **-** | **199 bp** |
| Fwd | ACC AGC CAG CTA TCA ACT CG | | |
| Rev | TTA CAT TGG TCC AGC CAC C | | |

**Supplemental Table 5**

| Score | 0 | 1 | 2 | 3 |
|---|---|---|---|---|
| **Appearance** | Smooth coat | Slightly roughed fur | Majority of fur on back is ruffled | Piloerection, puffy appearance |
| **Consciousness** | Active | Active, avoids standing upright | Active only when provoked | Non-responsive, even when provoked |
| **Activity** | Normal | Suppressed eating, drinking, or running | Stationary | Stationary, even when provoked |
| **Eyes** | Open | Not fully open, potential secretions | Half closed, potential secretions | No response to touch stimulus |
| **Posture** | Normal | Hunched, moving freely | Hunched, strained or stiff movement | Hunched, little or no movement |

**Supplemental Table 6**

| Mouse ID | Gender | Genotype | Tx | Group | Pre-TX Wei | Pre-TX Ten | Time post- | Post-TX We |
|---|---|---|---|---|---|---|---|---|
| 5129 | F | fl/+ | Saline | Saline_fl/+ | 21.2 | 37.5 | 16.5 | 21 |
| 5130 | F | fl/fl | Saline | Saline_fl/fl | 21.3 | 37.7 | 16.5 | 21.2 |
| 5132 | M | fl/fl | Saline | Saline_fl/fl | 19.2 | 37.4 | 16.5 | 18.1 |
| 5133 | F | fl/fl | Saline | Saline_fl/fl | 20.4 | 37.2 | 16.5 | 19.1 |
| 5135 | M | fl/fl | Saline | Saline_fl/fl | 21.9 | 37.9 | 16.5 | 21 |
| 5137 | M | fl/fl | LPS | LPS_fl/fl | 21.6 | 37.9 | 16.5 | 19.8 |
| 5139 | M | +/+ | Saline | Saline_+/+ | 23.2 | 37.2 | 16.5 | 22.4 |
| 5140 | F | +/+ | LPS | LPS_+/+ | 20.7 | 38.1 | 16.5 | 18.3 |
| 5142 | F | fl/+ | Saline | Saline_fl/+ | 17.3 | 38.1 | 16.5 | 16.7 |
| 5143 | F | fl/+ | LPS | LPS_fl/+ | 21.2 | 37.6 | 16.5 | 19.2 |
| 5144 | M | fl/+ | LPS | LPS_fl/+ | 24.9 | 37.9 | 16.5 | 22.9 |
| 5150 | M | fl/fl | Saline | Saline_fl/fl | 25.3 | 37.6 | 14.5 | 25.6 |
| 5151 | F | fl/fl | Saline | Saline_fl/fl | 19.9 | 37.4 | 14.5 | 19.4 |
| 5155 | F | fl/fl | LPS | LPS_fl/fl | 20.8 | 37.8 | 14.5 | 19.7 |
| 5156 | F | fl/fl | LPS | LPS_fl/fl | 22.4 | 37.8 | 14.5 | 19.9 |
| 5178 | M | fl/fl | LPS | LPS_fl/fl | 22.7 | 37.3 | 14.5 | 21.7 |
| 5179 | M | fl/fl | Saline | Saline_fl/fl | 24.4 | 37.2 | 14.5 | 24.9 |
| 5180 | F | +/+ | LPS | LPS_+/+ | 19.4 | 38.1 | 14.5 | 17.9 |
| 5181 | M | fl/+ | LPS | LPS_fl/+ | 22.5 | 37.1 | 14.5 | 21.9 |
| 5182 | F | fl/fl | Saline | Saline_fl/fl | 21.2 | 37.6 | 14.5 | 19.2 |
| 5183 | F | +/+ | Saline | Saline_+/+ | 19.8 | 38 | 14.5 | 21 |
| 5184 | M | fl/+ | Saline | Saline_fl/+ | 25.7 | 37.8 | 14.5 | 26.9 |
| 5185 | M | fl/+ | LPS | LPS_fl/+ | 26.1 | 37.8 | 14.5 | 23.9 |
| 5219 | M | +/+ | LPS | LPS_+/+ | 27.1 | 37.5 | 14.5 | 24.2 |
| 5220 | M | fl/+ | LPS | LPS_fl/+ | 28.2 | 37 | 14.5 | 26.5 |
| 5221 | M | fl/+ | saline | Saline_fl/+ | 25.4 | 37.2 | 14.5 | 25.6 |
| 5226 | M | +/+ | saline | Saline_+/+ | 25.1 | 37.7 | 14.5 | 25.4 |
| 5228 | M | fl/+ | LPS | LPS_fl/+ | 25.3 | 37.6 | 14.5 | 23 |
| 5230 | F | +/+ | LPS | LPS_+/+ | 20.2 | 37.5 | 14.5 | 18.5 |
| 5231 | F | fl/+ | LPS | LPS_fl/+ | 19.4 | 37.3 | 14.5 | 18.3 |
| 5232 | F | fl/fl | LPS | LPS_fl/fl | 20.7 | 37.9 | 14.5 | 19.6 |
| 5233 | M | fl/+ | LPS | LPS_fl/+ | 26.2 | 36.9 | 14.5 | 25 |
| 5234 | M | +/+ | LPS | LPS_+/+ | 28.1 | 36.9 | 14.5 | 26.1 |
| 5239 | F | fl/+ | LPS | LPS_fl/+ | 21.2 | 37.3 | 14.5 | 19.3 |
| 5240 | F | fl/+ | saline | Saline_fl/+ | 19.8 | 37.6 | 14.5 | 20 |
| 5241 | M | fl/+ | saline | Saline_fl/+ | 26.6 | 37.2 | 14.5 | 27.1 |
| 5242 | M | fl/fl | LPS | LPS_fl/fl | 22.9 | 37.6 | 14.5 | 21.3 |
| 5243 | M | +/+ | LPS | LPS_+/+ | 28.3 | 37.8 | 14.5 | 26.1 |
| 5244 | M | +/+ | LPS | LPS_+/+ | 28.4 | 37 | 14.5 | 26 |
| 5245 | F | +/+ | saline | Saline_+/+ | 20.8 | 37.4 | 14.5 | 21.2 |

| Post-TX Te | Delta Weig | Delta Temp | Appearanc | Consciousn | Activity | Eyes | Posture | Score |
|---|---|---|---|---|---|---|---|---|
| 38.2 | -0.2 | 0.7 | 0 | 0 | 0 | 0 | 0 | 0 |
| 37.9 | -0.1 | 0.2 | 0 | 0 | 0 | 0 | 0 | 0 |
| 38.4 | -1.1 | 1 | 0 | 0 | 0 | 1 | 0 | 1 |
| 38.1 | -1.3 | 0.9 | 0 | 0 | 0 | 0 | 0 | 0 |
| 37.9 | -0.9 | 0.1 | 0 | 0 | 0 | 0 | 0 | 0 |
| 35.7 | -1.8 | -2.2 | 1 | 2 | 2 | 1 | 2 | 8 |
| 38.1 | -0.8 | 0.8 | 0 | 0 | 0 | 0 | 0 | 0 |
| 36.2 | -2.4 | -1.9 | 1 | 1 | 0 | 2 | 1 | 5 |
| 38.2 | -0.6 | 0.1 | 0 | 0 | 0 | 0 | 0 | 0 |
| 34.4 | -2 | -3.2 | 1 | 1 | 1 | 2 | 2 | 7 |
| 37.1 | -2 | -0.9 | 1 | 0 | 0 | 1 | 1 | 3 |
| 37.6 | 0.3 | 0 | 0 | 0 | 0 | 0 | 0 | 0 |
| 37.7 | -0.5 | 0.3 | 0 | 0 | 0 | 0 | 0 | 0 |
| 33.9 | -1.1 | -3.9 | 2 | 3 | 3 | 0 | 3 | 11 |
| 37.2 | -2.5 | -0.6 | 0 | 0 | 1 | 1 | 0 | 2 |
| 33.9 | -1 | -3.4 | 1 | 3 | 3 | 2 | 3 | 12 |
| 37.7 | 0.5 | 0.4 | 0 | 0 | 0 | 0 | 0 | 0 |
| 36.3 | -1.5 | -1.8 | 1 | 2 | 2 | 2 | 1 | 8 |
| 37.8 | -0.6 | 0.7 | 0 | 0 | 0 | 0 | 0 | 0 |
| 37.7 | -2 | 0.2 | 0 | 0 | 0 | 0 | 0 | 0 |
| 37.6 | 1.2 | -0.4 | 0 | 0 | 0 | 0 | 0 | 0 |
| 38.1 | 1.2 | 0.2 | 0 | 0 | 0 | 0 | 0 | 0 |
| 36.2 | -2.2 | -1.6 | 1 | 1 | 1 | 1 | 1 | 5 |
| 37.2 | -2.9 | -0.3 | 0 | 1 | 1 | 1 | 0 | 3 |
| 36.8 | -1.7 | -0.2 | 1 | 1 | 1 | 1 | 1 | 5 |
| 38.1 | 0.2 | 0.9 | 0 | 0 | 0 | 0 | 0 | 0 |
| 37.8 | 0.3 | 0.1 | 0 | 0 | 0 | 0 | 0 | 0 |
| 36.9 | -2.3 | -0.7 | 1 | 1 | 1 | 2 | 1 | 6 |
| 35.8 | -1.7 | -1.7 | 1 | 1 | 2 | 1 | 3 | 8 |
| 34.6 | -1.1 | -2.7 | 1 | 2 | 3 | 2 | 2 | 10 |
| 33.9 | -1.1 | -4 | 2 | 3 | 3 | 1 | 3 | 12 |
| 36.4 | -1.2 | -0.5 | 2 | 1 | 1 | 2 | 1 | 7 |
| 36.8 | -2 | -0.2 | 1 | 0 | 1 | 1 | 1 | 4 |
| 35.2 | -1.9 | -2.1 | 1 | 1 | 2 | 2 | 2 | 8 |
| 38 | 0.2 | 0.4 | 0 | 0 | 0 | 0 | 0 | 0 |
| 38 | 0.5 | 0.8 | 0 | 0 | 0 | 0 | 0 | 0 |
| 34.5 | -1.6 | -3.1 | 2 | 3 | 3 | 2 | 3 | 13 |
| 37.3 | -2.2 | -0.4 | 1 | 0 | 0 | 0 | 1 | 2 |
| 37.4 | -2.4 | 0.4 | 1 | 0 | 0 | 1 | 1 | 3 |
| 37.9 | 0.4 | 0.5 | 0 | 0 | 0 | 0 | 0 | 0 |

| WBC | NEU | LYM | MONO | EOS | BAS | RBC | HGB | PLT |
|---|---|---|---|---|---|---|---|---|
| 7.53 | 1.08 | 6.33 | 0.04 | 0.05 | 0.03 | 10.22 | 15.7 | 1013 |
| 8.21 | 3.47 | 4.41 | 0.18 | 0.06 | 0.09 | 8.87 | 14.6 | 1038 |
| 3.19 | 1.2 | 1.92 | 0.04 | 0.02 | 0.01 | 9.6 | 15 | 1044 |
| 3.98 | 0.72 | 3.12 | 0.04 | 0.09 | 0.01 | 8.17 | 13.3 | 659 |
| 6.78 | 1.29 | 5.34 | 0.08 | 0.05 | 0.02 | 10.27 | 15.7 | 1197 |
| 5.17 | 3.8 | 0.76 | 0.34 | 0.2 | 0.07 | 8.76 | 13.5 | 559 |
| 5.95 | 1.21 | 4.61 | 0.08 | 0.03 | 0.02 | 10.8 | 16.3 | 957 |
| 2.92 | 1.77 | 0.8 | 0.19 | 0.12 | 0.04 | 8.33 | 13 | 576 |
| 6.07 | 1.24 | 4.61 | 0.12 | 0.06 | 0.04 | 10.08 | 15.7 | 1213 |
| 2.63 | 1.32 | 1.08 | 0.15 | 0.06 | 0.02 | 8.8 | 13.2 | 399 |
| 1.85 | 1.45 | 0.25 | 0.06 | 0.08 | 0.01 | 8.93 | 13.8 | 473 |
| 4.73 | 0.93 | 3.59 | 0.12 | 0.08 | 0.01 | 9.38 | 14.6 | 868 |
| 4.11 | 1.2 | 2.77 | 0.06 | 0.05 | 0.03 | 9.63 | 15.5 | 1146 |
| 2.35 | 1.43 | 0.61 | 0.22 | 0.07 | 0.02 | 9.03 | 15.1 | 658 |
| 1.71 | 1.12 | 0.38 | 0.15 | 0.05 | 0.01 | 8.86 | 14.4 | 374 |
| 3.62 | 2.2 | 0.7 | 0.49 | 0.19 | 0.04 | 8.2 | 13 | 377 |
| 5.34 | 0.82 | 4.41 | 0.07 | 0.03 | 0.01 | 9.46 | 14.7 | 1027 |
| 1.58 | 0.97 | 0.49 | 0.09 | 0.02 | 0.01 | 8.94 | 14.6 | 377 |
| 3.17 | 0.86 | 2.2 | 0.08 | 0.02 | 0.01 | 9.28 | 15.2 | 834 |
| 5.93 | 3.5 | 2.12 | 0.2 | 0.06 | 0.05 | 8.69 | 14.2 | 776 |
| 5.92 | 1.16 | 4.59 | 0.07 | 0.07 | 0.03 | 9.47 | 15.2 | 1353 |
| 5.1 | 1.22 | 3.63 | 0.13 | 0.11 | 0.01 | 9.24 | 14.5 | 1344 |
| 3.49 | 2.55 | 0.66 | 0.17 | 0.09 | 0.02 | 8.62 | 13.6 | 612 |
| 3.3 | 2.57 | 0.47 | 0.16 | 0.06 | 0.04 | 8.62 | 13.8 | 527 |
| 3.76 | 2.82 | 0.75 | 0.11 | 0.06 | 0.02 | 8.46 | 13.5 | 645 |
| 10.6 | 2.61 | 7.34 | 0.28 | 0.3 | 0.07 | 8.92 | 14.3 | 1453 |
| 6.64 | 2.22 | 4.02 | 0.23 | 0.12 | 0.05 | 9.12 | 14.4 | 1397 |
| 0.35 | 0.24 | 0.07 | 0.03 | 0.01 | 0 | 1.05 | 1.7 | 48 |
| 2.12 | 1.3 | 0.62 | 0.09 | 0.08 | 0.03 | 8.4 | 13.8 | 287 |
| 4.28 | 2.99 | 1.03 | 0.16 | 0.08 | 0.02 | 8.02 | 12.7 | 671 |
| 6.45 | 3.4 | 1.93 | 0.57 | 0.43 | 0.12 | 10.98 | 17.8 | 443 |
| 18.05 | 13.04 | 1.83 | 2.18 | 0.66 | 0.34 | 8.23 | 13.4 | 674 |
| 2.61 | 1.57 | 0.8 | 0.14 | 0.07 | 0.03 | 8.44 | 13.8 | 630 |
| 2.15 | 0.93 | 1 | 0.13 | 0.07 | 0.02 | 8.39 | 13.5 | 351 |
| 8.27 | 1.81 | 5.8 | 0.32 | 0.18 | 0.16 | 8.86 | 14.8 | 1111 |
| 6.19 | 1.4 | 4.37 | 0.2 | 0.18 | 0.04 | 8.77 | 14.7 | 684 |
| 4.59 | 3.31 | 0.78 | 0.27 | 0.16 | 0.07 | 9.24 | 15 | 538 |
| 2.73 | 1.89 | 0.44 | 0.18 | 0.16 | 0.06 | 8.08 | 13.4 | 436 |
| 3.21 | 2.43 | 0.45 | 0.16 | 0.13 | 0.04 | 8.16 | 13.7 | 333 |
| 12.34 | 1.94 | 9.8 | 0.25 | 0.28 | 0.07 | 8.69 | 14.7 | 1357 |

| Plasma IL-6 | Plasma TNI | pH | Sodium | Potassium | Chloride | Calcium, io | Lactate | BUN |
|---|---|---|---|---|---|---|---|---|
| 50.28 | 165.09 | 7.266 | 143 | 5.8 | 123 | 0.78 | 8.84 | 31 |
| 0 | 145.56 | 7.121 | 146 | 4.8 | 114 | 1.2 | 9.97 | 24 |
| 19.17 | 177.96 | 7.298 | 143 | 5.7 | 112 | 0.81 | 8.01 | 30 |
| 1.53 | 125.09 | 7.155 | 139 | 5.3 | 114 | 1.01 | 8.57 | 38 |
| 21.39 | 188.8 | 7.354 | 140 | 7.3 | 115 | 0.81 | 3.11 | 49 |
| 14614.72 | 412.87 | 7.193 | 141 | 7.7 | 123 | 0.63 | 8.22 | 115 |
| 57.78 | 146.76 | 7.119 | 140 | 6.3 | 114 | 0.84 | 8.65 | 32 |
| 1915.83 | 462.87 | | | | | | | |
| 74.72 | 145.46 | 6.888 | 143 | 5.6 | 115 | 1.08 | 14.25 | 35 |
| 15394.44 | 1098.75 | | | | | | | |
| 901.11 | 361.57 | 7.367 | 144 | 6.5 | 123 | 0.7 | 5.26 | 72 |
| 67.78 | 176.67 | 7.273 | 147 | 7.6 | 120 | 0.76 | 8.59 | 29 |
| 62.22 | 414.75 | 7.448 | 141 | 5.9 | 117 | 0.54 | 7.97 | 31 |
| 16021.39 | 249.72 | 7.311 | 138 | 10.7 | 123 | 0.53 | 10.69 | 103 |
| 273.61 | 205.09 | 7.235 | 147 | 7.6 | 121 | 1.05 | 8.09 | 58 |
| 19471.39 | 591.76 | | | | | | | |
| 58.61 | 166.02 | 7.085 | 144 | 4.8 | 116 | 1.04 | 11.1 | 26 |
| 4780.56 | 242.22 | | | | | | | |
| 19.03 | 179.91 | | | | | | | |
| 177.22 | 103.8 | 7.305 | 150 | 6.3 | 116 | 0.98 | 8.48 | 59 |
| 44.17 | 97.59 | 6.969 | 148 | 4.8 | 117 | 1.11 | 14.39 | 25 |
| 26.67 | 162.96 | 7.221 | 147 | 6.2 | 122 | 0.8 | 9.74 | 31 |
| 2417.78 | 302.69 | 7.159 | 147 | 5.2 | 115 | 1.02 | 7.65 | 81 |
| 1017.5 | 346.2 | 7.3 | 148 | 5.3 | 115 | 1.03 | 4.43 | 56 |
| 2418.89 | 330.56 | 7.28 | 148 | 5 | 119 | 1 | 3.94 | 87 |
| 49.17 | 227.04 | 7.266 | 147 | 4.5 | 114 | 1 | 7.04 | 29 |
| 34.17 | 163.06 | 7.281 | 147 | 4.7 | 113 | 1.06 | 6.44 | 25 |
| 976.39 | 210.37 | 7.342 | 147 | 6.1 | 124 | 0.84 | 3.6 | 70 |
| 2265.83 | 319.81 | 7.321 | 149 | 6.3 | 122 | 0.93 | 4.22 | 89 |
| 15201.94 | 375.28 | 7.272 | 146 | 5.5 | 120 | 0.82 | 5.57 | 112 |
| 19066.67 | 908.33 | | | | | | | |
| 5830.83 | 296.67 | 7.251 | 150 | 6.1 | 118 | 1.03 | 5.52 | 120 |
| 1151.94 | 254.17 | 7.305 | 149 | 4.8 | 116 | 1.12 | 2.7 | 73 |
| 9171.39 | 308.52 | 7.35 | 143 | 6.6 | 124 | 0.59 | 4.23 | 120 |
| 212.22 | 207.41 | | | | | | | |
| 80 | 121.67 | 7.291 | 148 | 4.5 | 114 | 1.11 | 5.99 | 26 |
| 19425.56 | 365.93 | | | | | | | |
| 623.89 | 194.26 | 7.281 | 146 | 8.4 | 120 | 0.94 | 3.34 | 66 |
| 1295.83 | 237.69 | 7.309 | 148 | 6.5 | 128 | 0.81 | 3.81 | 76 |
| 383.33 | 245.93 | 7.212 | 148 | 4.3 | 116 | 1.05 | 8.6 | 23 |

| Creatinine | Glucose | Comments | KIDNEY-ICA | Kidney_SEL | Kidney_IL6 | Kidney_SO | Kidney_STA | Kidney_CEI |
|---|---|---|---|---|---|---|---|---|
| 0.54 | 323 | | 0.974237 | 2.852996 | 0.429458 | 0.917319 | 1.331688 | 4.059992 |
| 0.51 | 285 | | 1.208472 | 9.704373 | 0.970508 | 1.567294 | 1.441893 | 3.798868 |
| 0.65 | 290 | | 0.777296 | 2.334025 | 1.001053 | 1.499879 | 1.354241 | 3.372349 |
| 0.63 | 367 | | 1.39187 | 12.26424 | 1.208547 | 2.178168 | 1.523736 | 3.906482 |
| 0.61 | 351 | | 1.50124 | 3.955029 | 1.790699 | 2.391218 | 1.510737 | 2.534373 |
| 1.23 | 42 | | 4.593022 | 59.14627 | 165.4945 | 41.83384 | 5.45406 | 6.947351 |
| 0.61 | 334 | | 1 | 1 | 1 | 1 | 1 | 1 |
| | | | 8.689115 | 39.91817 | 45.61319 | 27.0153 | 3.591692 | 9.752334 |
| 0.71 | 367 | | 1.022538 | 1.98099 | 1.128824 | 1.765676 | 1.554816 | 3.98246 |
| | | | 10.62946 | 61.36169 | 243.0559 | 76.68324 | 6.911494 | 14.55335 |
| 0.86 | 47 | | 8.303351 | 35.71454 | 36.39814 | 19.79696 | 3.283171 | 4.846128 |
| 0.74 | 253 | | 1.417461 | 0.890607 | 1.141455 | 1.187038 | 0.660242 | 0.267909 |
| 0.86 | 134 | | 1.352659 | 5.277981 | 0.196408 | 1.817712 | 0.003825 | 0.018837 |
| 1.39 | 89 | | 46.60823 | 12.60698 | 1264.124 | 61.87883 | 5.80375 | 2.150206 |
| 0.69 | 116 | No respons | 2.274288 | 2.505605 | 70.36199 | 1.060961 | 0.432337 | 0.167342 |
| | | | 13.04893 | 0.264578 | 0.045541 | 12.51965 | 4.08E-05 | |
| 0.58 | 314 | | 1.179205 | 0.567816 | 0.706134 | 0.63405 | 0.455436 | 0.39091 |
| | | | 17.22841 | 5.541168 | 307.2423 | 46.08009 | 3.170833 | 2.70292 |
| | | | 1.65539 | 0.319231 | 1.398057 | 0.498688 | 0.586179 | 0.410729 |
| 0.87 | 90 | Ascites and | 3.129167 | 3.674782 | 40.7873 | 5.060939 | 1.78649 | 11.2968 |
| 0.52 | 336 | | 1 | 1 | 1 | 1 | 1 | 1 |
| 0.79 | 261 | | 0.846414 | 0.400316 | 1.535373 | 0.657658 | 0.412856 | 0.341662 |
| 0.92 | 87 | | 12.59868 | 4.021119 | 111.9814 | 18.37697 | 2.026462 | 0.986205 |
| 0.88 | 80 | | 3.966549 | 1.339715 | 71.51308 | 4.92946 | 3.128638 | 1.436778 |
| 0.93 | 72 | | 7.577332 | 0.437029 | 19.55778 | 12.6963 | 1.999775 | 0.710962 |
| 0.54 | 343 | | 0.607498 | 0.641538 | 0.251707 | 0.287852 | 1.539597 | 0.115715 |
| 0.46 | 283 | | 1.125886 | 2.324485 | 2.259068 | 0.445606 | 0.683977 | 0.643742 |
| 0.91 | 61 | | 5.337212 | 1.132363 | 15.35605 | 2.512897 | 9.780663 | 1.704734 |
| 1.24 | 64 | | 8.05127 | 6.065337 | 215.4659 | 10.42297 | 3.190871 | 12.3662 |
| 1.17 | 73 | | 12.23406 | 3.591231 | 246.6908 | 47.63027 | 8.080584 | 1.829973 |
| | | | 26.55998 | 1.913857 | 2246.9 | 32.4628 | 6.215235 | 1.909086 |
| 1.21 | 89 | | 3.439425 | 11.13132 | 231.4011 | 93.47093 | 2.991525 | 2.188826 |
| 0.82 | 73 | | 4.526962 | 0.385928 | 14.42871 | 19.13539 | 7.818978 | 0.670756 |
| 1.35 | 60 | | 7.593538 | 3.030757 | 174.2235 | 14.37975 | 4.448839 | 7.93033 |
| | | | 0.535226 | 1.064961 | 0.434314 | 1.57377 | 0.985239 | 3.859571 |
| 0.4 | 297 | | 0.365172 | 0.455991 | 0.842696 | 0.600299 | 0.658247 | 0.286764 |
| | | | 9.008218 | 14.41105 | 534.9411 | 160.5271 | 5.981587 | 2.318516 |
| 1.64 | 94 | | 3.599245 | 1.436857 | 25.26896 | 2.671797 | 3.418219 | 0.480678 |
| 0.88 | 70 | | 10.5264 | 1.34671 | 16.05367 | 19.77755 | 2.973867 | 1.812258 |
| 0.51 | 267 | | 0.88819 | 0.430203 | 0.44266 | 2.244137 | 1.462037 | 1.553416 |

| Kidney_F3 | Kidney_RH | Kidney_TSI | Kidney_PLA | Kidney_PLA | Kidney_THI | Kidney_PLA | Kidney_SYN | Kidney_ET1 |
|---|---|---|---|---|---|---|---|---|
| 1.057323 | 1.258691 | 1.217768 | 2.823779 | 0.649582 | 1.28054 | 1.202053 | 0.921469 | 2.141539 |
| 1.139266 | 1.016983 | 0.666185 | 1.850634 | 0.927991 | 1.110789 | 1.173216 | 3.88909 | 0.00345 |
| 2.356474 | 1.35769 | 1.881335 | 1.604092 | 0.898269 | 1.23302 | 2.066932 | 1.088149 | 1.60886 |
| 1.087614 | 1.230023 | 1.428342 | 2.283677 | 1.240926 | 1.570563 | 1.923369 | 2.118398 | 1.812708 |
| 1.933402 | 1.876746 | 1.801265 | 1.93075 | 0.956443 | 1.37209 | 2.786103 | 1.266815 | 1.235826 |
| 1.616027 | 3.104751 | 0.392981 | 1.69354 | 0.347262 | 1.180555 | 14.12524 | 0.536997 | 5.097314 |
| 1 | 1 | 1 | 1 | 1 | 1 | 1 | 1 | 1 |
| 0.40286 | 2.763151 | 0.497363 | 2.888864 | 0.114009 | 0.656635 | 7.658922 | 0.598956 | 4.919259 |
| 0.637245 | 1.597086 | 1.248829 | 1.698231 | 1.083827 | 1.003369 | 1.441698 | 1.545709 | 1.283116 |
| 1.163839 | 5.931259 | 0.499789 | 2.82787 | 0.097415 | 1.278814 | 16.42584 | 0.363401 | 9.970851 |
| 0.828219 | 1.964488 | 0.465636 | 1.665712 | 0.215573 | 0.819921 | 4.156163 | 0.596082 | 2.99914 |
| 1.713227 | 0.440546 | 0.64636 | 0.4145 | 0.581444 | 0.625248 | 0.982766 | 0.426872 | 0.8357 |
| 0.077925 | 0.031252 | 0.098548 | 0.959995 | 0.565425 | 1.454527 | 2.466106 | 1.441182 | 5.481573 |
| 12.25846 | 3.598869 | 0.626026 | 1.513406 | 0.136103 | 0.961505 | 13.04509 | 0.209897 | 9.129408 |
| 1.692363 | 0.251015 | 0.15085 | 0.293549 | 0.004438 | 0.177242 | 1.15352 | 0.019291 | 2.506805 |
| 27.75485 | | 0.501061 | 0.689292 | 0.197727 | 0.925946 | 18.00772 | 0.045066 | 6.135763 |
| 1.473692 | 0.35686 | 0.763613 | 0.657891 | 0.823481 | 0.939521 | 0.980837 | 0.483249 | 1.103314 |
| 0.742628 | 2.641937 | 0.590959 | 1.26816 | 0.209662 | 1.52954 | 12.15742 | 0.368184 | 11.79939 |
| 2.021936 | 0.345358 | 0.70248 | 0.698905 | 0.84612 | 1.006105 | 0.814066 | 0.47342 | 0.820874 |
| 1.99935 | 1.440256 | 1.728904 | 0.710854 | 0.433919 | 1.56411 | 2.583207 | 1.333733 | 2.771811 |
| 1 | 1 | 1 | 1 | 1 | 1 | 1 | 1 | 1 |
| 0.743612 | 0.273703 | 0.585212 | 0.405551 | 0.280215 | 0.521544 | 1.355343 | 0.79091 | 0.737554 |
| 0.965542 | 1.419117 | 0.368562 | 0.943939 | 0.132544 | 0.918089 | 5.13636 | 0.464501 | 5.873538 |
| 3.668976 | 1.62366 | 1.016004 | 0.639548 | 0.67268 | 0.606868 | 1.068714 | 0.717379 | 2.272039 |
| 1.373564 | 1.239715 | 0.365896 | 0.459823 | 0.071816 | 0.730093 | 1.254842 | 0.540529 | 2.817334 |
| 1.446498 | 1.407074 | 1.864656 | 0.283917 | 0.521307 | 1.078218 | 0.74881 | 0.783938 | 0.489971 |
| 1.020426 | 0.732109 | 0.74284 | 0.826052 | 1.565189 | 0.731088 | 0.820866 | 0.973756 | 1.444527 |
| 1.494786 | 1.539379 | 0.580305 | 1.22718 | 0.281334 | 0.577986 | 0.564426 | 0.5561 | 4.262947 |
| 1.333841 | 2.793273 | 0.791069 | 1.4948 | 0.216151 | 0.82242 | 4.261499 | 1.028907 | 4.324138 |
| 1.023704 | 5.193143 | 1.047845 | 0.644021 | 0.066869 | 2.454119 | 12.87426 | 0.616505 | 5.158453 |
| 18.227 | 6.276613 | 1.317088 | 0.903343 | 0.173572 | 1.425006 | 8.254071 | 0.174486 | 3.48692 |
| 2.138984 | 4.502073 | 1.309968 | 2.802769 | 0.092359 | 1.482929 | 11.85937 | 0.545665 | 2.106023 |
| 1.849663 | 1.935095 | 0.668086 | 1.425753 | 0.317717 | 1.252098 | 3.659673 | 0.853306 | 3.259641 |
| 0.630713 | 3.62897 | 0.476838 | 1.815918 | 0.096868 | 0.869056 | 16.55658 | 0.509779 | 3.960636 |
| 0.898174 | 0.52835 | 1.5214 | 1.722858 | 1.490245 | 1.191348 | 1.189516 | 1.625266 | 0.667588 |
| 0.963784 | 0.29077 | 0.62443 | 0.439815 | 0.291037 | 0.298501 | 0.322227 | 0.718518 | 0.485793 |
| 10.81052 | 6.968803 | 2.031885 | 1.862846 | 0.388899 | 2.19637 | 44.53774 | 0.428191 | 5.376158 |
| 4.026236 | 1.189438 | 1.200352 | 1.159686 | 0.126971 | 0.522373 | 0.907386 | 0.286156 | 2.167945 |
| 4.171494 | 3.934029 | 1.80848 | 2.303274 | 0.150776 | 1.11594 | 1.386724 | 1.240083 | 6.349866 |
| 0.979983 | 1.365916 | 1.346184 | 1.210578 | 0.638901 | 1.367824 | 1.218226 | 1.026952 | 0.692268 |

| Kidney_PT | Kidney_VW | Kidney_PR | Kidney_CD | Kidney_TFF | Kidney_TFF | Kidney_TFF | Kidney_SEL | Kidney_C5A |
|---|---|---|---|---|---|---|---|---|
| 1.618082 | 3.578411 | 0.914892 | 5.692106 | 10.98911 | 9.162648 | 3.879646 | 2.410364 | 4.475729 |
| 0.689771 | 3.031839 | 1.024933 | 2.45636 | 5.830498 | 4.678356 | 3.294358 | 5.119811 | 4.79008 |
| 20.15517 | 12.87478 | 18.50614 | 62.74389 | 13.68899 | 16.74009 | 7.582413 | 24.90384 | 28.8247 |
| 1.932857 | 3.704499 | 0.878263 | 7.747412 | 6.90427 | 9.699717 | 2.744013 | 4.90011 | 9.966502 |
| 3.908874 | 4.562978 | 2.171122 | 10.86409 | 2.690986 | 3.198444 | 2.922025 | 3.795133 | 29.52239 |
| 16.90554 | 1.525868 | 87.92412 | 1.879674 | 13.0141 | 7.31785 | 5.386229 | 86.50613 | 46.47293 |
| 1 | 1 | 1 | 1 | 1 | 1 | 1 | 1 | 1 |
| 1.343108 | 0.25401 | 0.999894 | 0.971571 | 0.909511 | 0.166786 | 0.260678 | 14.5582 | 3.633242 |
| 1.149133 | 3.202734 | 1.960042 | 3.025912 | 5.70843 | 19.04964 | 4.776212 | 1.467577 | 8.60211 |
| 69.65625 | 2.774638 | 49.00389 | 21.74304 | 7.498534 | 1.033336 | 9.363383 | 297.1112 | 15.49427 |
| 0.892476 | 1.152249 | 0.415304 | 0.766974 | 0.542381 | 0.641368 | 0.570602 | 2.291713 | 1.341238 |
| 2.201334 | 2.860043 | 1.256756 | 2.304419 | 1.41225 | 0.789347 | 1.011219 | 3.774213 | 1.669259 |
| 14.94644 | 0.549022 | 2.375008 | 1.701399 | 0.332788 | 0.069952 | 0.443939 | 209.2635 | 3.04939 |
| 0.305012 | 0.058495 | 0.78011 | 1.351231 | 0.405097 | | | 132.4545 | 2.358206 |
| 9.71698 | 0.17073 | 1.710726 | 0.200592 | 0.16825 | 0.01563 | 0.235575 | 203.5777 | 2.328539 |
| 0.645483 | 0.972269 | 0.747019 | 0.172887 | 0.532885 | 0.58522 | 0.501866 | 2.407997 | 5.005608 |
| 13.47148 | 0.362357 | 0.943442 | 2.402734 | 0.688791 | 0.116292 | 0.816895 | 76.65233 | 2.179344 |
| 0.542088 | 0.54277 | 0.652085 | 0.219051 | 0.624258 | 0.333437 | 0.437656 | 0.991598 | 1.574796 |
| 2.58413 | 3.48513 | 4.484655 | 4.067675 | 1.085865 | 0.567882 | 0.869214 | 62.71843 | 3.593193 |
| 1 | 1 | 1 | 1 | 1 | 1 | 1 | 1 | 1 |
| 0.972512 | 0.780784 | 0.693175 | 0.562732 | 0.656045 | 0.370692 | 0.522011 | 1.88584 | 4.421217 |
| 4.226379 | 0.31727 | 1.278539 | 3.264792 | 0.237774 | 0.120042 | 0.429084 | 49.60468 | 2.166732 |
| 0.159596 | 3.220097 | 0.024355 | 0.843389 | 0.225396 | 0.15232 | 0.896937 | 6.011864 | 0.974537 |
| 3.644582 | 1.545261 | 0.082605 | 0.178603 | 0.284421 | 0.061313 | 0.939236 | 2.537934 | 0.150516 |
| 0.598155 | 6.385696 | 0.113213 | 0.294861 | 0.613223 | 1.575364 | 3.291672 | 0.557653 | 0.207232 |
| 0.871969 | 5.262016 | 0.274602 | 4.674749 | 0.562231 | 1 | 0.184859 | 1.12626 | 1.532326 |
| 0.6305 | 1.001584 | 0.035107 | 0.282064 | 0.715993 | 0.300644 | 2.526779 | 2.475167 | 1.328957 |
| 5.765329 | 3.788974 | 0.192434 | 1.253932 | 1.25172 | 0.71412 | 2.424352 | 22.02519 | 2.267829 |
| 14.27075 | 2.530843 | 0.079257 | 0.341681 | 0.923592 | 0.056179 | 2.09529 | 3.010327 | 0.581103 |
| 44.98771 | 3.771622 | 0.356396 | 6.933409 | 1.367373 | 0.382925 | 1.915303 | 12.801 | 0.870125 |
| 3.846302 | 0.978158 | 0.166436 | 2.35881 | 0.795261 | 0.105139 | 0.476462 | 7.014522 | 0.607962 |
| 1.447856 | 2.510743 | 0.189983 | 0.770692 | 0.235737 | 0.289793 | 2.145779 | 1.816069 | 1.171026 |
| 0.959044 | 3.016033 | 0.138868 | 0.929928 | 0.167691 | 0.052145 | 1.030552 | 8.584739 | 0.252176 |
| 0.528047 | 13.74778 | 0.097398 | 0.875716 | 1.780753 | 0.507452 | 1.965262 | 0.589438 | 0.366859 |
| 0.55933 | 4.677747 | 0.034909 | 0.083931 | 0.947957 | 0.31362 | 1.127393 | 0.451839 | 0.76774 |
| 36.10502 | 2.298122 | 1.498555 | 4.601005 | 0.375025 | 0.101269 | 2.756 | 68.53741 | 0.451712 |
| 0.762018 | 4.316045 | 0.124136 | 0.138187 | 0.607114 | 0.185265 | 0.969611 | 14.72554 | 2.569537 |
| 2.985954 | 1.807629 | 0.225189 | 0.365955 | 0.977111 | 0.473702 | 1.216085 | 10.89623 | 0.938146 |
| 1.146829 | 0.190041 | 0.060422 | 0.213915 | 1.778628 | | 5.409529 | 0.887895 | 0.652602 |

| Kidney_VE | Kidney_SE | Kidney_ST | Kidney_OA | Kidney_OA | Kidney_OA | Kidney_OA | Kidney_FO | Kidney_MX |
|---|---|---|---|---|---|---|---|---|
| 7.550772 | 0.785563 | 0.970788 | 1.355517 | 1.65738 | 1.129459 | 3.83364 | 1.229824 | 5.938431 |
| 5.737226 | 0.7429 | 1.409685 | 1.282556 | 2.457377 | 1.456103 | 2.41842 | 0.830025 | 5.132477 |
| 113.3095 | 11.37003 | 9.368323 | 3.290067 | 20.64553 | 10.20125 | 7.570303 | 12.45538 | 27.79988 |
| 7.499575 | 1.553318 | 1.672566 | 2.982856 | 4.063455 | 2.254855 | 12.34681 | 2.629598 | 8.345778 |
| 14.2351 | 5.75104 | 2.162551 | 3.062379 | 5.804372 | 4.384893 | 50.93285 | 1.726207 | 15.0424 |
| 18.74968 | 61.29244 | 22.4392 | 6.101658 | 24.7359 | 31.28116 | 65.00273 | 2.271371 | 377.3544 |
| 1 | 1 | 1 | 1 | 1 | 1 | 1 | 1 | 1 |
| 1.919823 | 4.030449 | 2.107893 | 2.982706 | 2.702402 | 5.758777 | 16.06108 | 0.179873 | 13.31603 |
| 20.42152 | 1.197071 | 3.89831 | 1.514393 | 2.393733 | 2.060194 | 1.923194 | 0.638893 | 9.813186 |
| 3.847415 | 72.90743 | 47.0799 | 83.49464 | 27.70148 | 20.69409 | 83.87313 | 5.005311 | 614.4603 |
| 1.737274 | 0.863579 | 0.014596 | 1.113297 | 0.433684 | 0.684212 | 0.299975 | 13.32225 | 1.112526 |
| 2.630672 | 0.793016 | 1.414524 | 1.630194 | 0.656333 | 3.500858 | 0.84348 | 18.08933 | 1.54928 |
| 0.655842 | 8.10844 | 4.487407 | 53.2961 | 3.094854 | 8.616383 | 4.635211 | 253.4276 | 509.9571 |
| 1.134609 | 0.095207 | 3.051156 | 3.358378 | 2.57671 | 3.545762 | 3.858326 | 8.401141 | 6.757729 |
| 0.274931 | 18.13624 | 4.938547 | 45.42375 | 0.652782 | 6.094467 | 9.99277 | 51.36713 | 57.3555 |
| 1.80827 | 0.767276 | 0.879088 | 3.345849 | 0.73639 | 0.472996 | 1.174374 | 19.54166 | 2.903521 |
| 0.579752 | 16.5402 | 9.351271 | 45.01132 | 1.390126 | 3.865663 | 4.387363 | 27.58515 | 20.70556 |
| 1.113328 | 0.414558 | 1.015427 | 12.42047 | 2.953844 | 1.1787 | 4.335767 | 9.36337 | 2.844199 |
| 1.298859 | 4.580058 | 0.838315 | 1.637266 | 0.733265 | 0.786157 | 0.809477 | 44.02319 | 0.923478 |
| 1 | 1 | 1 | 1 | 1 | 1 | 1 | 1 | 1 |
| 1.521513 | 1.098912 | 0.355783 | 1.348389 | 0.533049 | 0.977021 | 2.827769 | 25.66615 | 0.565964 |
| 0.477581 | 33.94944 | 1.898226 | 8.810006 | 4.079782 | 8.014408 | 2.625408 | 12.16294 | 21.48021 |
| 0.223136 | 0.445942 | 2.769634 | 5.428658 | 3.69269 | 5.39426 | 5.860832 | 4.214173 | 5.050679 |
| 0.154062 | 0.832791 | 2.135341 | 6.074734 | 2.178765 | 4.183174 | 3.263528 | 2.295268 | 8.018011 |
| 0.362859 | 0.094668 | 0.572418 | 0.676701 | 0.770857 | 0.709319 | 0.860643 | 2.69994 | 0.881915 |
| 0.483791 | 0.96708 | 0.828633 | 1.105317 | 0.96387 | 0.948466 | 1.034855 | 0.628787 | 0.639584 |
| 0.546899 | 1.036801 | 2.207552 | 6.590964 | 2.855589 | 3.252454 | 3.586691 | 0.808226 | 3.696715 |
| 0.784273 | 0.811855 | 1.84974 | 5.002817 | 3.389983 | 4.176974 | 4.432923 | 2.660776 | 3.924636 |
| 0.675913 | 2.131662 | 7.410609 | 55.54061 | 5.851673 | 13.18632 | 10.35894 | 4.641929 | 62.77758 |
| 0.43121 | 2.37201 | 6.889722 | 176.232 | 3.451295 | 3.915458 | 7.419172 | 32.70618 | 190.4934 |
| 0.413112 | 2.339845 | 1.973598 | 10.77244 | 3.022698 | | 3.396686 | 2.587232 | 8.747368 |
| 0.30964 | 2.363511 | 2.379186 | 4.432782 | 2.374057 | 3.576378 | 4.321529 | 1.168514 | 3.481418 |
| 0.274532 | 6.582252 | 1.900199 | 7.367572 | 2.039263 | 3.675006 | 1.682235 | 3.301288 | 34.23755 |
| 0.637229 | 0.589871 | 1.310141 | 1.187248 | 1.468511 | 0.89569 | 0.942749 | 0.915309 | 1.893131 |
| 0.864737 | 0.128453 | 0.429966 | 0.400504 | 0.460961 | 0.38722 | 0.38133 | 0.425156 | 0.940895 |
| 0.367233 | 2.827165 | 3.849412 | 16.8803 | 4.598807 | 6.506636 | 10.53265 | 9.926493 | 43.40602 |
| 0.218315 | 3.446665 | 2.971856 | 8.383599 | 4.800325 | 4.216299 | 3.296894 | 2.871544 | 8.281086 |
| 0.820601 | 1.959315 | 2.488977 | 7.070918 | 3.804859 | 3.454468 | 5.139757 | 3.087716 | 8.136296 |
| 2.06701 | 1.034041 | 1.206807 | 0.904718 | 1.037484 | 1.054334 | 0.966319 | 1.590363 | 1.563517 |

| Kidney_IL1 | Kidney_IRF | Kidney_IRF | Kidney_IRF | Kidney_CX( | Kidney_CX( | Kidney_IFII | Kidney_CL( | Kidney_HA |
|---|---|---|---|---|---|---|---|---|
| 1.135122 | 3.602059 | 1.644555 | 2.135557 | 1.832207 | 4.503615 | 3.289684 | 3.380111 | 0.243592 |
| 0.436549 | 1.686533 | 1.730471 | 1.225431 | 1.262316 | 1.05348 | 1.316296 | 1.424945 | 0.309917 |
| 4.040917 | 9.686005 | 8.026846 | 11.28967 | 14.10998 | 5.333953 | 10.27407 | 10.4203 | 22.51867 |
| 0.607427 | 4.650833 | 3.93549 | 1.391198 | 9.560688 | 2.615224 | 4.426814 | 6.471454 | 0.572959 |
| 1.071471 | 1.450692 | 3.100944 | 1.511708 | 8.099558 | 8.223731 | 1.567028 | 1.052703 | 2.442122 |
| 5.520664 | 15.43884 | 129.2883 | 22.81724 | 328.0641 | 727.1164 | 14.16893 | 78.09421 | 587.405 |
| 1 | 1 | 1 | 1 | 1 | 1 | 1 | 1 | 1 |
| 0.358944 | 1.889755 | 6.248297 | 0.996596 | 12.9667 | 88.66794 | 1.335729 | 7.047968 | 15.08155 |
| 1.448632 | 3.411605 | 2.722124 | 2.3078 | 3.189412 | 3.923905 | 2.230911 | 7.926777 | 0.183987 |
| 2.81681 | 36.9188 | 68.33384 | 25.01193 | 269.2039 | 1230.576 | 22.5029 | 89.65955 | 37.84671 |
| 1.755665 | 1.08721 | 0.532624 | 0.465317 | 10.62138 | 2.948518 | 1.210731 | 2.093598 | 2.603401 |
| 2.820414 | 1.363016 | 1.394126 | 0.870426 | 2.210994 | 2.670536 | 3.457726 | 3.87261 | 0.896762 |
| 2.151224 | 4.285979 | 9.087297 | 5.210411 | 130.9162 | 1014.821 | 13.3362 | 5.171677 | 11.08423 |
| 1.568388 | 1.445519 | 16.18403 | 0.320952 | 6.697089 | 142.7109 | 6.442898 | 10.4112 | 43.98008 |
| 4.265352 | 6.059991 | 16.09158 | 27.63754 | 1456.449 | 258.6611 | 9.471777 | 8.111839 | 175.2228 |
| 1.414208 | 0.433079 | 0.487169 | 1.49369 | 1.40714 | 1.423081 | 3.534006 | 2.134811 | 4.470602 |
| 5.015081 | 3.880005 | 28.37303 | 1.381057 | 35.01613 | 262.7096 | 22.1466 | 8.075992 | 39.53079 |
| 4.935949 | 0.526246 | 24.59161 | 0.560037 | 1.295844 | 51.99972 | 4.138797 | 1.385961 | 9.77013 |
| 2.333754 | 1.118267 | 0.801261 | 0.821021 | 3.992155 | 3.833995 | 2.062194 | 6.032653 | 11.51217 |
| 1 | 1 | 1 | 1 | 1 | 1 | 1 | 1 | 1 |
| 2.51034 | 0.690341 | 1.536053 | 0.950353 | 0.535558 | 0.783014 | 5.956618 | 3.05474 | 7.658426 |
| 0.969144 | 2.215597 | 8.046736 | 2.53057 | 92.04742 | 53.82385 | 8.8686 | 14.78371 | 136.6838 |
| 0.746381 | 1.430896 | 11.32888 | 0.702857 | 3.048654 | 30.34377 | 0.743885 | 0.315852 | 40.86505 |
| 0.55541 | 1.785752 | 16.40534 | 1.826591 | 6.341074 | 12.54088 | 1.376417 | 1.011957 | 55.46184 |
| 0.730385 | 1.382726 | 0.761613 | 1.18462 | 0.136248 | 0.83431 | 0.421012 | 0.193561 | 2.959964 |
| 0.525959 | 1.264125 | 1.537592 | 0.619315 | 1.488776 | 1.153328 | 0.401946 | 0.231809 | 0.898862 |
| 0.751678 | 1.489504 | 23.87347 | 1.654972 | 1.17298 | 12.43774 | 1.210856 | 1.703059 | 37.36908 |
| 0.54122 | 1.644478 | 10.8668 | 0.657098 | 7.594115 | 43.02949 | 0.569307 | 2.711824 | 21.59044 |
| 0.748664 | 4.088753 | 10.30953 | 1.707854 | 56.25378 | 321.6508 | 2.635084 | 1.512995 | 3.593746 |
| 0.542223 | 10.67984 | 5.066211 | 6.902737 | 45.90221 | 167.0014 | 5.011439 | 3.308253 | 1.224871 |
| 0.730642 | 0.838608 | 4.505553 | 2.33101 | 11.50431 | 50.95279 | 1.49148 | 3.419559 | 81.63431 |
| 0.716748 | 1.267642 | 9.2796 | 1.230836 | 3.908571 | 40.34837 | 0.742297 | 2.250267 | 72.69756 |
| 0.74422 | 4.587812 | 13.89017 | 2.0204 | 7.657814 | 71.69143 | 1.677958 | 7.501712 | 41.54528 |
| 1.229446 | 3.343009 | 1.79173 | 2.280087 | 0.26602 | 0.950608 | 1.640406 | 0.657623 | 0.505857 |
| 1.244154 | 1.765281 | 1.041269 | 1.158905 | 0.1658 | 3.109936 | 1.054382 | 0.673194 | 6.889959 |
| 1.898824 | 4.937506 | 7.627653 | 6.685005 | 93.39046 | 311.1584 | 3.296465 | 7.250023 | 195.5063 |
| 1.040131 | 1.864114 | 14.34431 | 0.987879 | 2.345847 | 31.3259 | 1.264659 | 1.948561 | 111.8757 |
| 1.01998 | 2.265803 | 12.45596 | 1.554711 | 7.80922 | 55.52307 | 2.350681 | 3.243923 | 184.2342 |
| 1.901291 | 3.428522 | 0.650367 | 1.614686 | 0.671693 | 0.867056 | 2.487898 | 4.313905 | 1.112518 |

| Kidney_LCN | Kidney_DD | Kidney_KLF | Lung_ICAM | Lung_SELE | Lung_IL6 | Lung_SOCS | Lung_STAT | Lung_CEMI |
|---|---|---|---|---|---|---|---|---|
| 5.200164 | 5.13886 | 1.40389 | 0.621084 | 0.351414 | 1.152132 | 0.565454 | 0.700751 | 0.486167 |
| 10.73199 | 2.743846 | 0.773832 | 0.651046 | 0.367264 | 1.011383 | 1.042525 | 1.722024 | 0.735775 |
| 28.34154 | 15.258 | 12.35783 | 0.594348 | 0.516205 | 0.89365 | 1.670416 | 0.885036 | 1.093703 |
| 5.300043 | 4.886144 | 1.99551 | 1.394163 | 0.285482 | 0.984176 | 0.704382 | 1.178391 | 0.667091 |
| 10.98367 | 3.334315 | 1.89081 | 0.868789 | 0.258318 | 0.392398 | 1.0206 | 1.249934 | 0.511702 |
| 33197.38 | 17.98816 | 2.912641 | 2.367168 | 13.11634 | 164.8695 | 10.27047 | 3.827644 | 21.51511 |
| 1 | 1 | 1 | 1 | 1 | 1 | 1 | 1 | 1 |
| 11547.12 | 2.664411 | 0.447268 | 11.39956 | 36.31614 | 101.9544 | 6.781499 | 1.291817 | 19.03781 |
| 9.348403 | 4.260629 | 1.335439 | 1.019757 | 0.508402 | 0.808003 | 0.702066 | 0.556043 | 0.771481 |
| 59505.17 | 24.32259 | 3.090663 | 2.716508 | 9.982392 | 326.9565 | 16.12985 | 2.446129 | 16.06376 |
| | | | 6.629145 | 20.52717 | 71.38962 | 6.175641 | 1.363394 | 14.11264 |
| 0.345695 | 0.592299 | 0.374333 | 0.958377 | 2.14425 | 0.432103 | 1.43424 | 0.571679 | 0.49359 |
| 0.749282 | 0.855562 | 0.857116 | 1.154005 | 1.058821 | 0.706892 | 1.397534 | 1.083612 | 0.999717 |
| 2739.989 | 14.44044 | 0.815355 | 7.298854 | 18.59433 | 259.349 | 36.65689 | 4.251097 | 15.23139 |
| 292.9192 | 1.925682 | 0.578188 | 30.41739 | 1.727627 | 7.055725 | | | |
| 1720.834 | 4.228446 | 0.820591 | 2.017122 | 39.2774 | 926.9563 | 19.80015 | 2.906463 | 15.37839 |
| 0.242277 | 1.255387 | 0.60743 | 0.948037 | 2.079197 | 0.942379 | 1.39885 | 1.014777 | 0.588622 |
| 6446.186 | 6.24284 | 1.396912 | 5.362327 | 31.50528 | 214.6922 | 12.05236 | 1.992884 | 33.26403 |
| 7.403231 | 2.485056 | 0.394313 | 1.865343 | 1.174215 | 1.168124 | 1.478285 | 1.765224 | 1.072512 |
| 118.4285 | 0.813299 | 1.060428 | 1.047213 | 3.776918 | 1.270384 | 3.267707 | 2.022249 | 2.336057 |
| 1 | 1 | 1 | 1 | 1 | 1 | 1 | 1 | 1 |
| 1.300479 | 0.843965 | 1.277393 | 2.353416 | 7.210053 | 1.90512 | 3.472316 | 1.101267 | 1.583726 |
| 2442.109 | 1.918481 | 1.162541 | 2.353416 | 7.210053 | 103.2129 | 3.472316 | 2.433456 | 14.84042 |
| 227.8639 | 0.853746 | 0.550768 | 4.059203 | 51.37915 | 145.9443 | 6.119209 | 2.850642 | 12.69425 |
| 267.6129 | 2.429125 | 0.468264 | 2.247911 | 36.91323 | 96.67953 | 9.053534 | 2.333404 | 14.61675 |
| 0.125777 | 1.55645 | 0.698151 | 1.169524 | 1.686078 | 2.219355 | 3.263237 | 1.243452 | 2.583754 |
| 0.52939 | 1.404186 | 0.703559 | 0.973357 | 1.505768 | 0.936536 | 1.409803 | 1.563828 | 0.507164 |
| 370.5148 | 0.665032 | 0.453514 | 5.613391 | 100.4459 | 54.23271 | 3.585972 | 2.40997 | 21.20079 |
| 612.128 | 0.427278 | 0.386843 | 3.15277 | 20.07462 | 84.74296 | 3.98934 | 1.741543 | 9.029514 |
| 459.9302 | 3.066318 | 1.371643 | 2.278363 | 20.13014 | 335.0484 | 11.3093 | 2.340049 | 24.15548 |
| 404.695 | 5.192288 | 1.304132 | 3.501664 | 59.46798 | 5087.134 | 32.66111 | 5.388775 | 52.90717 |
| 444.5444 | 0.977295 | 0.62469 | 2.461637 | 36.15207 | 83.57891 | 4.834408 | 1.173409 | 7.262508 |
| 665.3206 | 1.149242 | 0.258841 | 5.492868 | 51.89752 | 76.59388 | 7.276247 | 2.739948 | 9.899358 |
| 2050.22 | 2.166876 | 0.416342 | 5.665259 | 36.57416 | 216.3096 | 22.5418 | 4.089543 | 51.44162 |
| 0.767225 | 3.359108 | 1.376708 | 0.64533 | 1.231414 | 0.59554 | 1.618408 | 1.464158 | 0.779468 |
| 0.821296 | 0.627104 | 0.456728 | 0.837679 | 0.974851 | 0.873268 | 1.228944 | 1.264836 | 0.89023 |
| 1601.139 | 1.445369 | 0.249562 | 1.374992 | 10.80178 | 912.7319 | 16.63748 | 5.011048 | 31.02326 |
| 580.7737 | 1.782308 | 0.265786 | 3.512876 | 23.65379 | 23.71977 | 5.756026 | 2.344706 | 18.24352 |
| 746.8935 | 1.032717 | 0.601628 | 3.311173 | 40.91779 | 130.6038 | 6.115723 | 1.565559 | 26.28923 |
| 1.888965 | 1.503687 | 1.421345 | 0.438912 | 0.664113 | 1.067764 | 0.709319 | 0.639457 | 1.971748 |

| Lung_F3 | Lung_RHOI | Lung_TSP1 | Lung_PLAT | Lung_PLAU | Lung_THBD | Lung_PLAU | Lung_PTGS | Lung_VWF |
|---|---|---|---|---|---|---|---|---|
| 1.008212 | 0.429947 | 0.548266 | 0.761477 | 0.97766 | 0.691595 | 0.624948 | 1.047059 | 0.807341 |
| 1.00533 | 2.077038 | 0.830573 | 1.082035 | 0.942485 | 0.902756 | 0.904664 | 1.61296 | 2.048201 |
| 1.942887 | 0.691314 | 1.136824 | 1.362688 | 0.611988 | 0.879946 | 1.098489 | 1.827451 | 0.836768 |
| 0.982752 | 1.25269 | 1.011491 | 0.844521 | 0.639872 | 0.578091 | 0.563315 | 2.623861 | 2.134258 |
| 0.623162 | 0.87686 | 0.573215 | 1.330934 | 0.469761 | 1.468612 | 1.244162 | 1.11282 | 1.625369 |
| 1.959117 | 3.54081 | 0.848389 | 6.961253 | 0.129693 | 0.14522 | 6.289172 | 2.905875 | 0.147471 |
| 1 | 1 | 1 | 1 | 1 | 1 | 1 | 1 | 1 |
| 0.979284 | 1.3992 | 0.61784 | 6.41223 | 0.593206 | 0.161156 | 2.622702 | 8.918183 | 0.296829 |
| 0.877194 | 0.809876 | 0.877147 | 0.60083 | 0.344317 | 0.570867 | 0.452686 | 1.900672 | 1.249303 |
| 1.178513 | 3.735182 | 0.684173 | 6.566507 | 0.228906 | 0.132421 | 3.359537 | 2.746183 | 0.163805 |
| 1.108493 | 1.840734 | 0.643267 | 9.020696 | 0.100843 | 0.207836 | 1.978521 | 3.568797 | 0.235574 |
| 0.569387 | 0.602406 | 0.922521 | 0.514921 | 0.472801 | 0.729099 | 0.611973 | 1.551305 | 0.960865 |
| 0.634117 | 0.940477 | 1.851745 | 0.842927 | 0.812816 | 1.354592 | 1.047163 | 1.061734 | 0.837064 |
| 1.038079 | 6.049721 | 2.918542 | 4.769618 | 0.449043 | 0.144033 | 7.573677 | 2.762837 | 0.343502 |
| 0.299648 | 0.468684 | 1.535858 | 2.080363 | 0.373625 | 0.241998 | 1.221207 | 1.202141 | 0.323466 |
| 1.699282 | 3.490968 | 5.122838 | 3.447943 | 0.310968 | 0.177401 | 9.580325 | 1.534121 | 0.162718 |
| 0.690255 | 0.806792 | 1.209367 | 1.01179 | 1.020558 | 1.103705 | 0.735846 | 0.985028 | 0.842836 |
| 0.773889 | 1.701685 | 0.993323 | 6.436819 | 0.631449 | 0.191044 | 4.639785 | 7.229181 | 0.546705 |
| 0.724722 | 1.26436 | 1.130966 | 1.073734 | 2.601537 | 1.839934 | 1.426636 | 1.978545 | 1.888172 |
| 0.655082 | 1.712002 | 2.706557 | 1.672171 | 0.24149 | 0.733284 | 1.486933 | 1.23688 | 2.499035 |
| 1 | 1 | 1 | 1 | 1 | 1 | 1 | 1 | 1 |
| 0.853566 | 1.164457 | 1.693874 | 1.064179 | 1.29687 | 1.090616 | 0.937516 | 2.438954 | 1.141362 |
| 0.74122 | 1.860151 | 0.82018 | 8.720695 | 0.728309 | 0.238184 | 3.457411 | 3.428117 | 0.460046 |
| 0.858794 | 2.784236 | 0.841584 | 5.974723 | 0.863941 | 0.474451 | 2.161019 | 5.058818 | 0.342742 |
| 1.356004 | 3.495221 | 0.97124 | 7.417441 | 0.376587 | 0.258369 | 3.997975 | 3.607997 | 0.34789 |
| 1.234751 | 1.755106 | 2.505522 | 0.613768 | 1.0589 | 0.964774 | 1.850414 | 1.787655 | 1.517994 |
| 2.040473 | 1.820025 | 1.107402 | 0.769225 | 1.272602 | 2.088113 | 0.970852 | 2.054294 | 2.279332 |
| 1.879406 | 3.366513 | 1.754259 | 15.94382 | 0.577034 | 0.460654 | 4.610916 | 3.138655 | 0.487414 |
| 1.008477 | 2.226862 | 0.621196 | 7.231145 | 0.441708 | 0.238321 | 2.279593 | 2.586348 | 0.220932 |
| 1.212091 | 2.659614 | 0.919301 | 14.76665 | 0.321156 | 0.273647 | 5.582337 | 3.948264 | 0.398476 |
| 3.920089 | 14.45242 | 7.478782 | 12.87351 | 0.574642 | 0.367996 | 16.63485 | 9.929505 | 0.323448 |
| 0.969442 | 1.154742 | 0.594398 | 11.17564 | 0.305123 | 0.22377 | 4.272135 | 3.072529 | 0.176596 |
| 1.889521 | 1.769551 | 0.948453 | 9.920248 | 1.157422 | 0.445757 | 4.129828 | 4.008709 | 0.459137 |
| 4.152373 | 3.761207 | 1.337263 | 24.25441 | 0.883023 | 0.391402 | 8.809086 | 3.770807 | 0.242346 |
| 1.965699 | 1.618109 | 1.257743 | 4.436924 | 1.338017 | 1.997367 | 1.726844 | 0.874597 | 1.657853 |
| 1.460904 | 0.84363 | 0.727117 | 3.270207 | 0.581408 | 1.23836 | 1.276666 | 0.854441 | 1.531053 |
| 4.864892 | 3.848367 | 2.066156 | 10.66873 | 0.469997 | 0.146534 | 10.81498 | 2.871971 | 0.100756 |
| 1.264402 | 1.527847 | 1.754885 | 8.08821 | 0.388881 | 0.504043 | 2.811267 | 2.395303 | 0.442612 |
| 0.826384 | 1.558415 | 1.653277 | 7.887486 | 0.278642 | 0.399146 | 3.369634 | 1.633756 | 0.220406 |
| 0.490082 | 0.549443 | 0.903015 | 1.300009 | 0.785792 | 0.478901 | 1.030023 | 0.486785 | 0.438725 |

| Lung_PRO( | Lung_CD44 | Lung_TFPIA | Lung_TFPIE | Lung_TFPI( | Lung_SELP | Lung_C5AR | Lung_VEGF | Lung_SERP |
|---|---|---|---|---|---|---|---|---|
| 1.583058 | 1.139176 | 1.488493 | 1.403676 | 1.529417 | 0.31811 | 0.383304 | 0.20945 | 0.435087 |
| 1.637281 | 0.904038 | 1.717188 | 1.278593 | 1.577407 | 1.578212 | 0.482245 | 1.152854 | 0.500959 |
| 1.265927 | 1.919529 | 1.081591 | 1.149887 | 1.003072 | 1.4969 | 4.88721 | 0.396189 | 0.679428 |
| 1.361604 | 0.731589 | 1.779 | 1.512112 | 1.415588 | 3.958487 | 0.928404 | 2.159048 | 0.626927 |
| 1.02966 | 0.621874 | 1.614524 | 1.551883 | 1.038414 | 0.925527 | 0.8601 | 0.820833 | 0.908807 |
| 13.47138 | 1.494245 | 1.089698 | 0.284942 | 1.350779 | 98.9173 | 7.596611 | 0.46498 | 20.83983 |
| 1 | 1 | 1 | 1 | 1 | 1 | 1 | 1 | 1 |
| 7.495997 | 0.816356 | 0.743481 | 0.213531 | 0.77069 | 43.78746 | 7.474888 | 0.150072 | 19.71226 |
| 0.924786 | 1.823337 | 0.944689 | 0.864337 | 0.7917 | 0.953951 | 0.993344 | 0.315472 | 0.599222 |
| 5.834948 | 0.992532 | 0.76359 | 0.103945 | 0.955972 | 50.02585 | 6.772746 | 0.179954 | 6.457361 |
| 5.643699 | 0.892716 | 0.858795 | 0.324501 | 0.756201 | 17.51195 | 9.497119 | 0.198239 | 11.0948 |
| | 0.932581 | 1.230682 | 1.170234 | 0.992928 | 1.955491 | 1.010247 | 0.667893 | 1.027792 |
| | 1.2117 | 1.373404 | 1.158135 | 0.7968 | 1.824415 | 0.64335 | 1.031661 | 1.674551 |
| | 1.741777 | 0.578324 | 0.197301 | 0.676662 | 55.39461 | 12.49747 | 0.645716 | 11.2436 |
| | 0.641924 | 0.965449 | 0.610126 | 0.649485 | 24.70005 | 1.716076 | 0.200011 | 20.99388 |
| | 1.992465 | 0.579278 | 0.132602 | 0.549259 | 106.2707 | 8.342281 | 0.264151 | 21.19955 |
| | 0.761556 | 1.058128 | 0.973591 | 0.977885 | 2.95124 | 0.57285 | 0.740254 | 1.245793 |
| | 1.082374 | 0.646769 | 0.177154 | 0.612451 | 23.48229 | 7.221186 | 0.689398 | 23.77842 |
| | 0.621869 | 0.816403 | 0.952249 | 0.822702 | 0.913389 | 2.091222 | 1.150515 | 1.04153 |
| | 0.730355 | 1.343445 | 0.801787 | 1.147498 | 5.306311 | 3.270944 | 0.772779 | 2.235449 |
| | 1 | 1 | 1 | 1 | 1 | 1 | 1 | 1 |
| | 1.757065 | 0.780746 | 0.804772 | 0.736591 | 0.960045 | 1.194037 | 1.052191 | 1.091321 |
| | 1.543663 | 0.89433 | 0.287474 | 0.886158 | 27.47807 | 13.0911 | 0.742827 | 26.42723 |
| 8.483696 | 1.180014 | 1.1519 | 0.405851 | 0.95352 | 34.30583 | 1.394397 | 0.167299 | 25.73082 |
| 9.105125 | 2.019425 | 0.615625 | 0.324689 | 0.720989 | 43.91123 | 3.741908 | 0.106999 | 23.47194 |
| 2.563212 | 3.334646 | 1.304417 | 0.785299 | 1.523763 | 0.798449 | 0.147588 | 0.661616 | 2.017196 |
| 1.5947 | 3.334002 | 2.40473 | 1.392206 | 2.128683 | 1.603543 | 1.58374 | 1.418535 | 1.667049 |
| 13.41694 | 3.043924 | 0.70477 | 0.564896 | 1.052556 | 24.63421 | 10.30214 | 0.459601 | 61.01154 |
| 9.958829 | 1.616175 | 1.514261 | 0.450622 | 1.071138 | 45.77661 | 0.871604 | 0.337727 | 25.49211 |
| 36.46589 | 3.806922 | 0.869945 | 0.117478 | 0.582321 | 81.02728 | 5.459564 | 0.257797 | 76.54726 |
| 18.46905 | 0.38287 | 0.19484 | 0.180944 | 1.211379 | 102.9318 | 1.574685 | 0.172506 | 31.79112 |
| 31.32646 | 0.607778 | 1.069438 | 0.129185 | 0.166011 | 156.2101 | 6.911668 | 0.305012 | 45.75123 |
| 10.62479 | 2.125462 | 0.748717 | 0.430124 | 1.456805 | 44.06564 | 2.618978 | 0.495431 | 30.35386 |
| 28.2467 | 3.359946 | 1.188439 | 0.160832 | 0.894179 | 151.8738 | 2.67823 | 0.756111 | 23.71455 |
| 4.692415 | 3.319743 | 2.13789 | 1.338492 | 0.903074 | 1.336593 | 1.945005 | 1.818039 | 5.062488 |
| 1.556427 | 1.392251 | 1.933491 | 1.381818 | 1.076996 | 1.385561 | 1.26145 | 0.465198 | 1.116888 |
| 15.99673 | 2.732967 | 0.820032 | 0.233229 | 1.29474 | 68.86903 | 0.394697 | 0.383298 | 29.8431 |
| 5.700794 | 1.035634 | 1.565117 | 0.650633 | 0.690651 | 19.75798 | 0.971496 | 0.07677 | 23.73079 |
| 4.759914 | 0.704002 | 0.414215 | 0.535238 | 0.633364 | 6.011435 | 3.752798 | 0.1961 | 25.77914 |
| 0.627077 | 0.29994 | 0.415847 | 0.718284 | 0.469774 | 0.623619 | 0.631417 | 0.704952 | 0.599863 |

| Lung_STAT | Lung_OASL | Lung_OAS2 | Lung_OASL | Lung_OAS3 | Lung_FOS | Lung_MX1 | Lung_IL18 | Lung_IRF1 |
|---|---|---|---|---|---|---|---|---|
| 0.813316 | 0.64326 | 0.693336 | 0.948527 | 0.424692 | 0.630025 | 1.009758 | 1.250724 | 1.065526 |
| 1.074643 | 1.532526 | 2.256412 | 1.713796 | 1.736932 | 0.943891 | 2.221763 | 1.246571 | 1.045307 |
| 0.807384 | 1.053378 | 1.414791 | 1.507059 | 1.643338 | 0.835492 | 0.664995 | 1.534024 | 0.99487 |
| 1.352835 | 1.017932 | 1.094384 | 1.329647 | 0.835512 | 0.960296 | 1.271511 | 1.033523 | 1.44707 |
| 1.161819 | 2.181376 | 2.582326 | 1.740152 | 1.589219 | 0.700275 | 0.828816 | 1.156067 | 1.213614 |
| 3.172866 | 9.067052 | 2.839938 | 4.210862 | 5.823419 | 1.38096 | 9.182159 | 0.639519 | 1.832912 |
| 1 | 1 | 1 | 1 | 1 | 1 | 1 | 1 | 1 |
| 2.544314 | 9.177183 | 3.909525 | 4.201304 | 10.85614 | 0.624529 | 5.241675 | 0.571189 | 1.782514 |
| 0.762441 | 0.59618 | 0.709209 | 0.795743 | 0.459252 | 0.789178 | 0.68815 | 1.13331 | 0.860677 |
| 4.408499 | 22.71788 | 5.681791 | 5.547926 | 8.546993 | 1.835735 | 31.35689 | 0.433555 | 3.777702 |
| 2.560779 | 11.07217 | 4.006155 | 5.391622 | 10.69505 | 0.329479 | 5.718718 | 0.815578 | 1.629474 |
| 1.041738 | 1.295844 | 1.785994 | 1.054683 | 1.356709 | 3.365325 | 0.743469 | 0.902196 | 1.064441 |
| 1.267642 | 1.138456 | 1.638912 | 1.390589 | 1.2292 | 2.208429 | 0.753604 | 0.712555 | 1.054762 |
| 5.719489 | 93.39447 | 9.094917 | 7.075874 | 15.68993 | 10.86526 | 79.95781 | 0.551978 | 6.602616 |
| 2.82326 | 9.088943 | 5.379084 | 5.50712 | 9.450951 | 3.809797 | 5.916387 | 0.79396 | 1.496876 |
| 4.333521 | 131.2546 | 7.69941 | 5.129906 | 20.25966 | 13.14314 | 64.01015 | 0.44895 | 3.540843 |
| 1.027147 | 1.13986 | 1.414853 | 1.261367 | 0.644719 | 2.917339 | 1.043987 | 0.950819 | 1.020959 |
| 3.78778 | 18.9789 | 4.156272 | 7.635437 | 16.78108 | 7.334597 | 14.49344 | 0.709412 | 2.594818 |
| 1.476049 | 2.306443 | 2.096942 | 1.94222 | 2.35402 | 2.267314 | 2.726413 | 1.58225 | 0.987344 |
| 0.926214 | 0.923538 | 1.812454 | 1.46787 | 3.095451 | 6.724788 | 0.572032 | 0.550571 | 1.07672 |
| 1 | 1 | 1 | 1 | 1 | 1 | 1 | 1 | 1 |
| 0.873955 | 1.522014 | 1.167155 | 1.165177 | 0.863157 | 3.479343 | 0.948364 | 1.373996 | 0.80459 |
| 4.481532 | 19.77959 | 7.107743 | 7.569893 | 29.7575 | 5.799186 | 8.398786 | 0.599967 | 2.081455 |
| 2.921612 | 12.84233 | 5.823241 | 7.909149 | 9.480604 | 6.301278 | 5.705438 | 0.990302 | 1.540872 |
| 3.649717 | 12.40583 | 5.315951 | 6.794009 | 5.344318 | 4.548935 | 6.28394 | 0.579153 | 1.34016 |
| 2.702638 | 2.198475 | 2.613205 | 2.898486 | 1.644326 | 5.46794 | 3.573095 | 1.68096 | 0.641092 |
| 1.772227 | 1.987448 | 1.943083 | 1.444453 | 1.596188 | 1.991884 | 1.800188 | 1.59336 | 0.848926 |
| 5.890732 | 14.47093 | 7.468299 | 10.83565 | 3.588377 | 3.624709 | 9.412426 | 1.072252 | 1.086151 |
| 3.368383 | 7.769434 | 4.56704 | 6.148382 | 3.992221 | 2.391783 | 5.548194 | 0.611753 | 1.011892 |
| 43.58324 | 160.3653 | 41.54739 | 52.76907 | 9.023493 | 28.1565 | 94.24741 | 1.677455 | 2.971029 |
| 16.60374 | 233.8148 | 17.62383 | 13.41633 | 13.26843 | 27.27057 | 172.8386 | 0.908543 | 7.830647 |
| 5.18147 | 11.8372 | 9.632193 | 8.945507 | 4.045116 | 7.454573 | 9.171198 | 1.071553 | 0.830699 |
| 4.771574 | 5.21778 | 5.056478 | 7.621237 | 4.105426 | 2.056078 | 3.32813 | 1.128657 | 1.029404 |
| 10.348 | 18.98841 | 5.998106 | 10.53161 | 6.68828 | 5.371114 | 27.06444 | 1.114389 | 2.587696 |
| 8.346507 | 4.803925 | 7.607415 | 6.389306 | 1.631023 | 4.751709 | 1.683632 | | 1.177606 |
| 2.347981 | 2.627419 | 2.430218 | 2.583006 | 0.534787 | 1.183345 | 1.552645 | 1.273918 | 0.697299 |
| 2.768274 | 13.41105 | 3.042274 | 4.505044 | 9.655059 | 6.562041 | 12.24573 | 0.368553 | 2.77076 |
| 3.707968 | 10.66287 | 5.742356 | 7.340282 | 8.679844 | 1.992261 | 5.793887 | 0.903641 | 1.204185 |
| 2.181145 | 7.340388 | 4.62071 | 4.563509 | 4.782598 | 1.603879 | 3.468245 | 0.42747 | 1.239861 |
| 0.564262 | 0.503158 | 0.514646 | 0.692304 | 0.626493 | 0.502037 | 0.555498 | 0.627604 | 1.17796 |

| Lung_IRF7 | Lung_IRF8 | Lung_CXCL | Lung_CXCL | Lung_IFIH1 | Lung_DDX5 | Lung_KLF2 | Liver_ICAM | Liver_SELE |
|---|---|---|---|---|---|---|---|---|
| 0.912463 | 0.911668 | 0.317822 | 3.505235 | 1.204914 | 0.899921 | 0.750718 | 1.097665 | 1.142831 |
| 1.427107 | 1.215659 | 0.65303 | 2.044911 | 1.383065 | 1.158654 | 0.611308 | 0.668434 | 0.967041 |
| 0.872648 | 0.621996 | 1.03921 | 1.081554 | 1.053674 | 0.984547 | 1.094319 | 1.015899 | 1.36882 |
| 1.252294 | 1.174271 | 0.282102 | 1.22795 | 1.434801 | 1.375432 | 0.985508 | 1.363407 | 0.439799 |
| 1.524784 | 0.791593 | 1.090819 | 0.906489 | 1.351816 | 1.323575 | 0.844685 | 0.852297 | 0.753091 |
| 12.01161 | 1.418695 | 30.1209 | 593.5298 | 2.588578 | 1.622529 | 0.760578 | 7.224862 | 1.91586 |
| 1 | 1 | 1 | 1 | 1 | 1 | 1 | 1 | 1 |
| 9.393189 | 0.808105 | 38.65234 | 301.0334 | 2.228809 | 1.938172 | 0.629909 | 1.547331 | 0.960612 |
| 0.677468 | 0.77829 | 0.380347 | 0.478701 | 0.99137 | 0.789661 | 0.627549 | 4.721572 | 0.933415 |
| 17.31768 | 1.939226 | 40.66857 | 1692.092 | 3.659878 | 2.326459 | 0.783172 | 7.963836 | 1.378339 |
| 10.75717 | 0.71524 | 25.25159 | 415.7267 | 2.522673 | 1.811711 | 0.507417 | 3.649977 | 1.335602 |
| 0.745183 | 1.21742 | 0.424843 | 0.661389 | 1.417546 | 1.81726 | 1.056556 | 1.354365 | 0.484804 |
| 0.865455 | 1.086826 | 0.812863 | 0.810433 | 1.47726 | 1.785408 | 0.735132 | 1.962121 | 0.573426 |
| 15.89637 | 3.477641 | 518.3292 | 1847.898 | 5.053892 | 4.314661 | 2.532123 | 15.22397 | 6.412264 |
| 15.65428 |  | 166.8273 | 156.0345 | 2.637564 | 2.487741 | 0.409979 | 7.88156 | 2.319015 |
| 20.99388 | 3.158411 | 2437.928 | 1852.861 | 4.257813 | 3.328537 | 1.690183 | 15.10438 | 5.131249 |
| 1.471406 | 1.091656 | 1.390174 | 0.596445 | 1.169229 | 1.293403 | 0.535123 | 1.013136 | 1.093706 |
| 16.52967 | 1.844535 | 258.2695 | 1688.293 | 3.052177 | 2.328364 | 3.826819 | 10.61421 | 1.17318 |
| 2.738016 | 1.165759 | 5.636355 | 1.359853 | 1.431005 | 1.217544 | 2.834124 | 2.251394 | 0.290246 |
| 0.87147 | 0.703151 | 10.2078 | 0.48206 | 1.140128 | 1.081759 | 1.673456 | 0.327189 | 0.627676 |
| 1 | 1 | 1 | 1 | 1 | 1 | 1 | 1 | 1 |
| 1.054567 | 0.877805 | 2.420026 | 0.700314 | 0.921972 | 0.888898 | 1.607908 | 1.009709 | 0.171895 |
| 11.93393 | 1.44693 | 144.3432 | 838.3742 | 3.811409 | 3.884219 | 4.971886 | 4.848416 | 0.981759 |
| 19.12926 | 1.164591 | 4.8367 | 53.17853 | 2.408311 | 2.162445 | 5.336936 | 3.757301 | 1.70472 |
| 15.6283 | 1.219753 | 8.425333 | 36.16973 | 1.974848 | 1.637168 | 0.683755 | 2.117994 | 3.093067 |
| 1.092875 | 0.663626 | 0.658403 | 1.023685 | 0.826588 | 0.738696 | 1.153112 | 0.956435 | 1.958879 |
| 1.067235 | 0.813844 | 1.787118 | 1.35138 | 0.75305 | 0.807878 | 0.921554 | 1.383957 | 0.327738 |
| 11.65716 | 0.861678 | 2.543722 | 12.64766 | 1.734281 | 1.403825 | 2.390058 | 5.975232 | 2.03219 |
| 11.93081 | 0.95041 | 5.577867 | 32.3697 | 1.661967 | 1.110118 | 0.761932 | 5.771125 | 4.41065 |
| 10.45852 | 2.954121 | 29.01146 | 89.4303 | 4.873122 | 2.938054 | 0.053263 | 22.49837 | 15.54818 |
| 6.939723 | 6.942476 | 77.18795 | 257.2242 | 8.213913 | 6.842198 | 7.791137 | 29.32253 | 50.06773 |
| 7.349691 | 1.193599 | 16.08654 | 27.97721 | 1.749422 | 1.271273 | 4.375321 | 4.469591 | 2.84844 |
| 12.46221 | 1.05063 | 2.342678 | 12.70522 | 1.508001 | 1.270323 | 0.631734 | 1.373827 | 2.458365 |
| 14.70265 | 2.125993 | 20.32246 | 94.9036 | 2.799996 | 2.455555 | 8.897173 | 10.17732 | 3.595288 |
| 1.59191 | 0.986201 | 0.370543 | 1.204003 | 1.181574 | 1.170189 | 6.945358 | 1.236005 | 2.50345 |
| 0.967372 | 0.689389 | 0.657346 | 1.370978 | 0.809893 | 0.627895 | 3.42635 | 1.236628 | 0.634857 |
| 13.88376 | 4.624707 | 44.12656 | 117.5193 | 2.337866 | 1.798032 | 6.138773 | 21.39476 | 7.586534 |
| 15.9829 | 1.049052 | 3.99621 | 14.26988 | 1.507413 | 1.632334 | 4.710817 | 5.65675 | 2.21195 |
| 12.12261 | 0.893291 | 3.891708 | 15.36225 | 1.633958 | 1.581729 | 2.415854 | 1.616502 | 3.466331 |
| 0.937001 | 1.228737 | 0.55956 | 0.739984 | 1.327933 | 1.237811 | 1.085123 | 0.722566 | 3.051216 |

| Liver_IL6 | Liver_SOCS | Liver_STAT | Liver_CEMI | Liver_F3 | Liver_RHOI | Liver_TSP1 | Liver_PLAT | Liver_PLAU |
|---|---|---|---|---|---|---|---|---|
| 0.345844 | 0.973153 | 0.498498 | 0.194902 | 4.942041 | 2.704289 | 1.032015 | 1.516717 | 1.551397 |
| 1.025394 | 1.698094 | 1.259763 | 1.750906 | 1.693952 | 1.168934 | 2.00828 | 1.229924 | 1.517022 |
| 0.681423 | 1.602833 | 2.706507 | 0.231667 | 1.348059 | 3.218861 | 0.817259 | 0.345934 | 0.199947 |
| 1.629294 | 0.922549 | 1.174951 | 0.191519 | 3.293104 | 2.366473 | 1.580296 | 0.267519 | 0.421906 |
| 0.689398 | 0.29362 | 0.82454 | 1.580375 | 0.767496 | 1.97057 | 1.700821 | 0.750612 | 1.103284 |
| 13.68776 | 30.43622 | 2.087857 | 52.18084 | 0.418558 | 0.878534 | 1.806806 | 2.614432 | 0.524168 |
| 1 | 1 | 1 | 1 | 1 | 1 | 1 | 1 | 1 |
| 1.030334 | 0.963092 | 1.318332 | 1.918909 | 0.68318 | 1.47596 | 0.73878 | 1.954949 | 1.290005 |
| 5.214167 | 12.14035 | 1.342916 | 28.54234 | 0.376163 | 0.641533 | 0.878804 | 1.318653 | 0.741147 |
| 14.3275 | 28.93313 | 2.071137 | 64.56097 | 0.564757 | 0.695973 | 1.204706 | 2.331999 | 0.388934 |
| 2.381299 | 9.644336 | 0.735658 | 4.586912 | 0.164369 | 0.379888 | 0.356745 | 0.318337 | 0.117652 |
| 2.466546 | 8.825814 | 1.906188 | 2.116284 | 0.531025 | 1.470634 | 0.264304 | 2.145209 | 0.36268 |
| 2.817756 | 9.323619 | 2.148561 | 4.818077 | 0.690958 | 2.869299 | 1.274111 | 1.591008 | 0.820385 |
| 73.4939 | 86.18866 | 2.803788 | 57.76101 | 0.649423 | 0.766666 | 2.701127 | 0.709944 | 0.583698 |
| 5.607961 | 30.83598 | 1.682867 | 2.234066 | 0.484862 | 1.333947 | 0.781284 | 5.407856 | 1.25241 |
| 128.3318 | 88.44058 | 3.663703 | 68.40126 | 1.081533 | 1.324308 | 8.057616 | 1.748015 | 0.406642 |
| 0.912531 | 2.806002 | 1.055568 | 1.502207 | 1.135157 | 1.143413 | 0.320799 | 1.234958 | 0.478674 |
| 104.0114 | 62.02085 | 1.702184 | 47.26019 | 0.438919 | 1.024179 | 1.388473 | 1.177717 | 0.523193 |
| 1.253116 | 1.424169 | 1.136653 | | 1.174549 | 1.264673 | 0.442923 | 1.107198 | 2.181979 |
| 1.211601 | 19.06834 | 1.199058 | 38.54589 | 0.384725 | 0.777456 | 1.024902 | 0.676765 | 0.434813 |
| 1 | 1 | 1 | 1 | 1 | 1 | 1 | 1 | 1 |
| 1.141016 | 3.895996 | 1.103864 | 0.895449 | 0.957496 | 1.075213 | 0.269872 | 1.433321 | 0.542506 |
| 17.50751 | 31.64454 | 0.848809 | 6.489343 | 0.132671 | 0.465951 | 0.366681 | 0.976477 | 0.133829 |
| 4.776257 | 6.90979 | 0.891108 | 0.250317 | 0.389862 | 0.453756 | 1.83208 | 1.358644 | 0.251944 |
| 2.619996 | 9.86349 | 1.379249 | 1.97013 | 0.227177 | 0.311517 | 2.30944 | 1.353017 | 0.619648 |
| 1.311282 | 1.954286 | 1.226506 | 1.841966 | 0.529038 | 1.905098 | 2.813623 | 0.155995 | 1.450412 |
| 0.60917 | 0.929806 | 1.148306 | 1 | 0.773073 | 0.799458 | 0.779032 | 0.040074 | 0.251319 |
| 3.088396 | 2.684712 | 0.50677 | 0.114207 | 0.331154 | 0.287506 | 0.905186 | 0.627949 | 0.172898 |
| 11.42562 | 9.535965 | 1.039304 | 0.664591 | 0.713538 | 0.728727 | 1.746353 | 13.00698 | 0.331001 |
| 29.32412 | 14.04696 | 0.82243 | 7.64091 | 0.444156 | 0.25983 | 3.163975 | 0.609976 | 0.508453 |
| 130.4162 | 81.70223 | 3.387168 | 31.65894 | 1.468842 | 2.953082 | 16.42203 | 9.235265 | 1.021094 |
| 5.138846 | 7.047562 | 0.612625 | 4.172845 | 0.398231 | 0.340432 | 2.360763 | 1.577563 | 0.202614 |
| 2.43977 | 4.961154 | 0.317325 | 0.338024 | 0.267198 | 0.257973 | 0.339792 | 5.582204 | 0.11108 |
| 14.24785 | 13.04434 | 0.567616 | 4.191207 | 0.834802 | 0.300024 | 2.612195 | 12.6759 | 0.291915 |
| 3.424878 | 2.346431 | 1.106153 | 1.186551 | 1.249781 | 1.177946 | 1.702467 | 12.70451 | 0.767181 |
| 1.665459 | 2.625035 | 0.952081 | 0.275595 | 2.236766 | 0.551439 | 0.794417 | 1.597244 | 0.287698 |
| 68.23002 | 52.82236 | 1.795477 | 25.23554 | 0.795557 | 2.058536 | 4.764712 | 12.19213 | 0.534184 |
| 7.958584 | 15.9924 | 1.199457 | 1.713216 | 0.553925 | 0.60654 | 1.927023 | 7.572845 | 1.213902 |
| 2.906263 | 12.18332 | 1.060653 | 0.216711 | 0.409842 | 0.580555 | 1.66998 | 9.714219 | 0.519601 |
| 1.641579 | 1.075493 | 0.870848 | | 1.293539 | 1.250848 | 1.283644 | 24.9536 | 3.979008 |

| Liver_THBD | Liver_PLAU | Liver_ET1 | Liver_PTGS | Liver_VWF | Liver_PROC | Liver_CD44 | Liver_TFPIA | Liver_TFPIB |
|---|---|---|---|---|---|---|---|---|
| 0.91594 | 0.498558 | 1.077652 | 0.398927 | 2.675456 | 0.709068 | 0.134515 | 1.528053 | 1.687396 |
| 0.673938 | 4.197656 | 3.101252 | 2.152627 | 5.06389 | 1.279465 | 0.801129 | 2.376343 | 2.421789 |
| 1.637956 | 0.801111 | 2.151201 | 0.474178 | 2.096895 | 0.504966 | 0.191189 | 1.474949 | 2.069817 |
| 0.702612 | 0.404083 | 1.26357 | 2.875467 | 1.158356 | 3.184402 | 0.263164 | 5.369783 | 5.923039 |
| 2.533429 | 1.007535 | 1.818483 | 2.160673 | 3.11812 | 0.511627 | 0.222233 | 2.457945 | 1.909962 |
| 6.518742 | 13.38637 | 6.941512 | 11.94055 | 0.368442 | 9.034708 | 2.587993 | 0.995003 | 0.358223 |
| 1 | 1 | 1 | 1 | 1 | 1 | 1 | 1 | 1 |
| 0.790413 | 0.833485 | 3.057674 | 3.998631 | 2.260601 | 2.740757 | 0.864771 | 1.417158 | 2.992795 |
| 5.462676 | 5.056893 | 2.431732 | 8.440507 | 0.470084 | 10.81428 | 0.297707 | 1.771932 | 0.508305 |
| 2.341935 | 8.422994 | 5.643901 | 99.81758 | 0.639495 | 10.48134 | 1.550888 | 0.826679 | 0.119323 |
| 2.214912 | 2.198676 | 1.220913 | 3.21253 | 0.385477 | 14.0149 | 1.693028 | 1.896112 | 0.861148 |
| 0.883944 | 0.434953 | 2.441348 | | 2.450752 | 0.645475 | | 0.965439 | 1.161019 |
| 2.069921 | 1.092876 | 2.83087 | | 1.586332 | 0.684763 | | 1.105132 | 1.446664 |
| 4.71723 | 27.29118 | 9.797876 | | 0.821745 | 0.949835 | | 0.897891 | 0.229697 |
| 4.96451 | 4.703511 | 2.985725 | | 0.80783 | 8.935224 | | 0.981796 | 1.720718 |
| 3.202709 | 42.51704 | 54.50823 | | 2.162025 | 18.31135 | | 1.597521 | 0.365248 |
| 1.263887 | 0.286825 | 1.29472 | | 0.633221 | 1.724806 | | 0.699659 | 0.51536 |
| 2.348018 | 9.349541 | 3.982086 | | 1.053009 | 28.26558 | | 0.918109 | 0.402149 |
| 1.47363 | 0.905331 | 2.082595 | | 2.284876 | 2.057175 | | 3.260729 | 1.227013 |
| 2.755046 | 2.12943 | 0.841852 | | 1.929536 | 3.083704 | | 1.948871 | 1.549573 |
| 1 | 1 | 1 | | 1 | 1 | | 1 | 1 |
| 0.859045 | 0.332758 | 1.107138 | | 1.225165 | 1.207906 | | 1.03426 | 0.964626 |
| 3.428874 | 3.228117 | 1.349159 | | 0.214107 | 4.74748 | | 0.345583 | 0.116182 |
| 4.257217 | 1.26082 | 1.591372 | 1.414307 | 0.186358 | 10.4118 | 0.299273 | 0.943677 | 0.558202 |
| 2.667636 | 1.72247 | 0.697055 | 3.909721 | 0.183751 | 3.623909 | | 0.671104 | 0.14201 |
| 1.377998 | 0.297956 | 0.508596 | 0.852038 | 1.149163 | 1.346823 | | 0.473142 | 0.967389 |
| 1.563399 | 0.142205 | 0.813592 | 0.839041 | 0.927541 | 1.150729 | 1.068913 | 0.911 | 0.751614 |
| 1.864517 | 1.050101 | 1.716285 | 0.829853 | 0.13638 | 6.248615 | 0.094132 | 0.621855 | 0.350159 |
| 5.488353 | 2.665186 | 3.306327 | 4.56893 | 0.575964 | 18.47513 | 0.350679 | 1.000571 | 0.722872 |
| 4.680559 | 3.632397 | 4.926854 | 30.46722 | 0.306627 | 22.23633 | 0.745844 | 0.764431 | 0.119283 |
| 5.333459 | 28.03888 | 19.69874 | 204.0951 | 2.867034 | 13.77944 | 4.5699 | 0.914133 | 0.332269 |
| 1.728513 | 2.199472 | 1.42493 | 2.848877 | 0.18603 | 4.768814 | 0.347293 | 0.44766 | 0.120114 |
| 3.047318 | 0.581513 | 0.964318 | 0.431397 | 0.085803 | 3.740818 | 0.099008 | 0.734755 | 0.183977 |
| 2.517502 | 3.804381 | 3.395985 | 3.374686 | 0.721737 | 28.243 | 0.340264 | 0.766421 | 0.277824 |
| 2.740837 | 0.383163 | 0.785846 | 0.131636 | 1.341615 | 2.022788 | 0.690432 | 0.929963 | 1.465874 |
| 0.803624 | 0.261391 | 0.597014 | 1.89714 | 0.812951 | 0.993945 | | 0.862864 | 1.623966 |
| 6.445394 | 4.96014 | 10.24857 | 51.78265 | 0.472911 | 21.65685 | 0.892924 | 1.274981 | 0.206312 |
| 2.982497 | 0.92216 | 1.775433 | 1.831403 | 0.590243 | 8.521184 | 0.057867 | 1.642687 | 0.779684 |
| 5.828317 | 1.24341 | 1.371887 | 1.197362 | 0.388576 | 6.696792 | | 1.133162 | 0.728816 |
| 0.639632 | 0.3825 | 1.229117 | 1.191837 | 1.07812 | 0.869014 | 0.93553 | 1.097695 | 1.330471 |

| Liver_TFPI( | Liver_SELP | Liver_C5AF | Liver_VEGF | Liver_SERP | Liver_STAT | Liver_OASL | Liver_OAS2 | Liver_OASL |
|---|---|---|---|---|---|---|---|---|
| 1.555231 | 1.938008 | 0.617183 | 1.649269 | 0.645737 | 5.551837 | 1.597215 | 1.200502 | 0.962322 |
| 1.631156 | 22.85341 | 4.501502 | 0.979981 | 1.453843 | 3.171436 | 3.876841 | 1.469433 | 2.087435 |
| 0.825859 | 4.704176 | 1.05264 | 1.519001 | 0.464396 | 5.866918 | 1.492726 | 1.325698 | 0.98182 |
| 4.248457 | 16.00637 | 0.586359 | 0.605423 | 1.209479 | 4.378081 | 0.621837 | 0.765335 | 0.542621 |
| 1.090506 | 11.89314 | 0.474452 | 0.966482 | 0.624028 | 2.375815 | 2.841962 | 1.377099 | 2.228597 |
| 1.508197 | 99.17513 | 0.881251 | 0.979425 | 377.6404 | 14.50943 | 1.822459 | 1.573647 | 1.158385 |
| 1 | 1 | 1 | 1 | 1 | 1 | 1 | 1 | 1 |
| 1.182993 | 3.87348 | 3.269021 | 0.562024 | 0.461688 | 1.96401 | 0.593275 | 0.468714 | 1.993316 |
| 1.460172 | 34.16972 | 5.05118 | 0.639681 | 34.45815 | 2.850029 | 0.984027 | 1.319246 | 0.97734 |
| 0.620811 | 45.56931 | 3.388258 | 1.077878 | 93.86038 | 1.464143 | 3.436203 | 0.625025 | 0.549396 |
| 1.792516 | 17.85286 | 1.744432 | 0.331883 | 144.1111 | 0.679388 | 0.56087 | 0.128636 | 1.022981 |
| 1.226786 | 10.3092 | 3.966114 | 1.803538 | 1.002717 | 1.855669 | 1.537423 | 1.304752 | 0.941613 |
| 1.346029 | 28.77923 | 3.534613 | 3.765496 | 1.268492 | 0.471211 | 0.54971 | 2.386456 | 1.386587 |
| 1.590291 | 194.5141 | 23.47912 | 3.123806 | 533.1372 | 5.51846 | 11.10099 | 6.42885 | 8.768302 |
| 2.620519 | 162.2713 | 17.13605 | 4.553354 | 13.1486 | 5.524636 | 2.76455 | 10.86455 | 7.901655 |
| 2.235011 | 559.9715 | 38.87409 | 5.364702 | 388.6463 | 0.815741 | 10.96954 | 15.05092 | 9.375671 |
| 0.89643 | 2.540469 | 1.513848 | 1.606465 | 0.283184 | 0.803781 | 0.710554 | 0.277187 | 0.328828 |
| 1.893149 | 24.37068 | 6.33605 | 7.698779 | 200.3893 | 3.793117 | 2.969533 | 3.094437 | 6.743315 |
| 1.874061 | 2.781973 | 2.526572 | 2.058432 | 1.313763 | 1.580291 | 2.269369 | 0.333619 | 2.712994 |
| 1.704538 | 105.2351 | 2.326801 | 6.973347 | 63.18141 | 2.625589 | 0.276736 | 2.262625 | 0.534429 |
| 1 | 1 | 1 | 1 | 1 | 1 | 1 | 1 | 1 |
| 0.787879 | 1.723566 | 2.415241 | 4.14207 | 0.751299 | 1.082554 | 1.66426 | 0.703597 | 0.775877 |
| 0.509602 | 93.75013 | 13.14171 | 1.682729 | 150.1046 | 1.472204 | 0.585358 | 0.459959 | 2.298276 |
| 1.302085 | 17.93907 | 6.689709 | 0.285867 | 36.83418 | 0.939387 | 0.711162 | 1.129693 | 2.625408 |
| 0.829696 | 40.40751 | 10.36071 | 0.874983 | 122.5121 | 0.740323 | 0.350577 | 0.636691 | 1.53814 |
| 0.819188 | 0.820835 | 1.049582 | 1.169284 | 0.917747 | 0.846797 | 1.469405 | 0.480727 | 0.85117 |
| 0.572689 | 0.318613 | 0.828918 | 0.649072 | 0.829537 | 0.995165 | 0.760937 | 0.912565 | 0.86682 |
| 0.92353 | 4.115759 | 2.673475 | 0.648592 | 81.68667 | 0.795716 | 0.564558 | 0.887421 | 1.973558 |
| 1.418748 | 8.727269 | 17.21849 | 0.509279 | 133.6908 | 0.83964 | 1.030505 | 1.799255 | 3.102224 |
| 0.889312 | 4.692083 | 34.0166 | 0.152629 | 358.9577 | 1.14269 | 3.116919 | 2.524733 | 5.419451 |
| 0.716072 | 129.6149 | 50.7674 | 7.824396 | 1196.502 | 0.944297 | 15.7397 | 2.78221 | 3.322245 |
| 0.409478 | 1.350203 | 6.47975 | 0.234475 | 101.4608 | 0.517784 | 0.384001 | 0.554499 | 1.192548 |
| 0.775825 | 8.377826 | 8.529237 | 0.45708 | 21.15947 | 0.693228 | 0.339275 | 0.453957 | 1.020698 |
| 0.942271 | 26.39099 | 29.27612 | 0.340622 | 156.9959 | 0.963446 | 1.663697 | 1.841661 | 3.668341 |
| 1.309607 | 1.413929 | 17.94353 | 0.479553 | 0.744056 | 1.034528 | 0.937547 | 1.735964 | 1.491095 |
| 1.177126 | 2.330566 | 5.33475 | 0.302772 | 0.950248 | 1.395033 | 0.888602 | 0.807236 | 1.374283 |
| 1.677435 | 31.32296 | 23.90722 | 2.029022 | 1436.009 | 1.072626 | 1.768799 | 2.107011 | 3.468586 |
| 1.340191 | 11.13712 | 3.567325 | 1.904544 | 81.58867 | 1.785418 | 1.172735 | 1.692384 | 3.404986 |
| 1.307286 | 42.55202 | 19.6298 | 1.213226 | 200.7347 | 1.252875 | 0.67247 | 1.08895 | 2.013432 |
| 1.746147 | 3.138601 | 1.206392 | 1.54066 | 1.205492 | 1.004858 | 1.314169 | 1.095812 | 1.153642 |

| Liver_OAS3 | Liver_FOS | Liver_MX1 | Liver_IL18 | Liver_IRF1 | Liver_IRF7 | Liver_IRF8 | Liver_CXCL | Liver_CXCL |
|---|---|---|---|---|---|---|---|---|
| 4.561965 | 0.073312 | 0.76362 | 1.535785 | 1.346032 | 1.151529 | 1.062377 | 0.388727 | 1.941686 |
| 4.757501 | 0.228238 | 2.941793 | 1.018623 | 1.419049 | 1.356938 | 1.577505 | 1.029166 | 2.983807 |
| 1.910174 | 0.155917 | 1.275658 | 1.170777 | 1.150111 | 1.033764 | 1.426547 | 0.979183 | 1.588011 |
| 1.800803 | 0.106104 | 0.559134 | 1.088711 | 1.178477 | 1.113827 | 1.182652 | 0.686334 | 1.551298 |
| 6.103103 | 0.020547 | 1.253739 | 1.684811 | 0.928448 | 1.743231 | 2.001772 | 0.726405 | 2.73301 |
| 2.342576 | 0.007328 | 2.003785 | 0.203879 | 0.686115 | 1.644635 | 2.040646 | 13.348 | 133.9218 |
| 1 | 1 | 1 | 1 | 1 | 1 | 1 | 1 | 1 |
| 0.786727 | 0.02014 | 0.445992 | 1.123152 | 0.977514 | 0.854139 | 0.962288 | 1.321189 | 0.829501 |
| 3.3669 | 0.008115 | 0.954712 | 0.176804 | 0.384188 | 0.735083 | 0.976374 | 4.05931 | 22.59905 |
| 3.377123 | 0.00582 | 5.236301 | 0.175396 | 0.903353 | 1.365727 | 2.625131 | 17.05327 | 97.67908 |
| 7.038637 | 0.0022 | 1.262853 | 0.106113 | 0.385789 | 0.669852 | 0.626611 | 3.822233 | 61.55588 |
| 0.276456 | 3.768006 | 6.274638 | 2.30268 | 2.708647 | 3.236929 | 2.223397 | | 0.416272 |
| 2.428359 | 13.07247 | 6.203561 | 2.936617 | 2.345142 | 2.431851 | 3.517584 | 8.935118 | 1.023729 |
| 3.064541 | 7.442003 | 73.33579 | 0.558008 | 6.04418 | 3.377623 | 13.90917 | 349.48 | 488.7991 |
| 5.216656 | 7.851577 | 8.2423 | 1.04211 | 2.785716 | 9.418532 | 1.393462 | 29.07643 | 96.34828 |
| 11.45391 | 17.84809 | 179.4998 | 0.978999 | 2.460322 | 7.777265 | 42.76096 | 2253.427 | 501.3947 |
| 0.469933 | 0.621298 | 3.047193 | 0.56195 | 0.909439 | 1.430031 | 5.503626 | 5.184725 | 0.562461 |
| 7.207356 | 5.195299 | 9.661462 | 0.263211 | 2.109616 | 2.598807 | 9.85449 | 249.8566 | 110.2935 |
| 0.995657 | 1.03986 | 2.367046 | 1.384093 | 1.02654 | 3.190989 | 4.102488 | 9.492562 | 53.4252 |
| 0.732954 | 33.18378 | 0.585405 | 0.460053 | 0.765752 | 0.313057 | 1.863817 | 9.062678 | 1.621585 |
| 1 | 1 | 1 | 1 | 1 | 1 | 1 | 1 | 1 |
| 0.394095 | 1.975534 | 3.187177 | 1.128561 | 1.392455 | 2.238107 | 2.20376 | 1.241897 | 0.595519 |
| 2.079832 | 2.047443 | 1.76302 | 0.171572 | 1.245725 | 1.499901 | 2.262523 | 82.29745 | 53.52184 |
| 5.151999 | 3.12437 | 1.172067 | 0.280131 | 0.546567 | 1.556349 | 1.279366 | 10.27426 | 130.5625 |
| 2.419586 | 1.326891 | 0.884113 | 0.088989 | 0.426205 | 1.206318 | 1.195426 | 12.13478 | 134.8918 |
| 0.706509 | 3.877772 | 1.313637 | 0.64341 | 0.459492 | 1.170617 | 0.978329 | 3.504211 | 0.664265 |
| 0.704556 | 0.691911 | 1.039492 | 0.843446 | 0.964534 | 0.914982 | 0.602342 | 1.132203 | 0.488681 |
| 2.612944 | 0.863247 | 1.064377 | 0.197746 | 0.352773 | 1.192724 | 1.00516 | 8.533117 | 40.44771 |
| 5.766941 | 2.867534 | 1.315059 | 0.377234 | 0.622693 | 1.488315 | 1.006486 | 18.58369 | 47.44248 |
| 12.04996 | 2.562396 | 9.68322 | 0.195292 | 0.913259 | 1.728805 | 2.662228 | 38.22076 | 101.201 |
| 8.294974 | 7.614987 | 72.22842 | 0.281314 | 1.693551 | 1.202158 | 5.01351 | 422.9397 | 534.5318 |
| 1.845524 | 0.797466 | 0.575037 | 0.155209 | 0.267307 | 0.607496 | 1.068801 | 12.33398 | 59.08374 |
| 1.160642 | 0.609042 | 0.774739 | 0.175314 | 0.33253 | 0.94805 | 0.622703 | 3.902234 | 50.51199 |
| 8.830395 | 2.086369 | 7.274775 | 0.290187 | 0.877628 | 2.000699 | 3.629742 | 27.74491 | 62.52677 |
| 3.081147 | 2.084965 | 1.44488 | 1.583965 | 1.320494 | 1.895607 | 2.277818 | 2.335797 | 2.241601 |
| 0.56448 | 0.948179 | 1.521603 | 1.008914 | 1.279083 | 1.357602 | 0.815914 | 2.64243 | 1.120156 |
| 6.241457 | 6.622789 | 5.18682 | 0.303705 | 1.169275 | 1.743082 | 5.331964 | 128.062 | 151.6876 |
| 5.60154 | 1.53839 | 2.414828 | 0.299664 | 0.994354 | 2.33828 | 1.989575 | 19.92399 | 87.12362 |
| 3.704668 | 1.240149 | 1.55157 | 0.249145 | 0.654257 | 1.549381 | 1.02442 | 13.65099 | 74.43902 |
| 1.419334 | 1.445272 | 0.962008 | 1.185613 | 1.03677 | 1.092917 | 1.660185 | 0.883234 | 2.046325 |

| Liver_IFIH1 | Liver_DDX5 | Liver_KLF2 | Liver_HNF1 |
|---|---|---|---|
| 1.407919 | 1.238724 | 0.909539 | 1.05739 |
| 1.447384 | 1.404987 | 0.880276 | 1.259822 |
| 1.35081 | 1.870749 | 3.427519 | 1.288668 |
| 1.216836 | 1.072773 | 0.737294 | 0.787958 |
| 1.82272 | 2.006452 | 2.176902 | 1.507968 |
| 0.533641 | 0.542672 | 1.831985 | 0.91133 |
| 1 | 1 | 1 | 1 |
| 0.922347 | 1.135824 | 0.971922 | 0.849919 |
| 0.293966 | 0.381305 | 0.748178 | 0.621281 |
| 0.542544 | 0.391232 | 10.23667 | 0.63476 |
| 0.302831 | 0.337728 | 2.977163 | 0.353006 |
| 0.966616 | 0.859037 | 3.273839 | 2.562631 |
| 0.998053 | 1.285024 | 2.880546 | 1.663199 |
| 1.428671 | 1.140455 | 4.050856 | 1.831064 |
| 0.762935 | 1.467435 | 2.39222 | 2.105832 |
| 1.62555 | 0.909296 | 9.502708 | 2.168841 |
| 0.862711 | 0.617206 | 0.321588 | 0.619223 |
| 0.547277 | 0.610517 | 3.512133 | 1.742708 |
| 1.165027 | 1.316728 | 1.363529 | 1.266497 |
| 0.426058 | 0.620286 | 1.577947 | 1.192167 |
| 1 | 1 | 1 | 1 |
| 1.020528 | 1.071316 | 0.994239 | 2.099874 |
| 0.246212 | 0.247538 | 1.048564 | 1.360049 |
| 0.593841 | 0.888622 | 0.997219 | 0.653571 |
| 0.336015 | 0.609406 | 1.026397 | 0.766231 |
| 0.826929 | 1.64158 | 1.175846 | 1.292318 |
| 0.938891 | 0.58242 | 0.439121 | 0.629095 |
| 0.437439 | 0.383464 | 1.62814 | 0.273309 |
| 0.501329 | 0.673539 | 1.734863 | 0.714958 |
| 0.936579 | 0.527358 | 2.301555 | 0.422349 |
| 1.573088 | 2.328082 | 8.036089 | 0.224717 |
| 0.318589 | 0.342923 | 1.18745 | 0.569503 |
| 0.523348 | 0.395953 | 0.791079 | 0.379035 |
| 0.614405 | 0.594651 | 1.74727 | 0.665762 |
| 1.275806 | 1.744142 | 1.149437 | 1.290505 |
| 1.362586 | 0.716858 | 0.341969 | 0.851684 |
| 0.839301 | 0.783682 | 3.463626 | 1.538454 |
| 0.901741 | 1.531206 | 0.978531 | 0.977878 |
| 0.588796 | 0.780075 | 1.045923 | 1.02817 |
| 1.065086 | 1.716974 | 2.277278 | 1.589584 |

**Supplemental Table 7**

| Mouse ID | Gender | Genotype | Tx | Group | KIDNEY-ICA | Kidney_SEL | Kidney_IL6 | Kidney_SO |
|---|---|---|---|---|---|---|---|---|
| 5129 | F | fl/+ | Saline | Saline_fl/+ | -0.03765 | 1.512478 | -1.21941 | -0.1245 |
| 5130 | F | fl/fl | Saline | Saline_fl/fl | 0.273184 | 3.278635 | -0.04319 | 0.648275 |
| 5132 | M | fl/fl | Saline | Saline_fl/fl | -0.36346 | 1.22282 | 0.001518 | 0.584846 |
| 5133 | F | fl/fl | Saline | Saline_fl/fl | 0.477024 | 3.616386 | 0.273273 | 1.123116 |
| 5135 | M | fl/fl | Saline | Saline_fl/fl | 0.586155 | 1.983688 | 0.840523 | 1.257746 |
| 5137 | M | fl/fl | LPS | LPS_fl/fl | 2.199444 | 5.886215 | 7.37064 | 5.386599 |
| 5139 | M | +/+ | Saline | Saline_+/+ | 0 | 0 | 0 | 0 |
| 5140 | F | +/+ | LPS | LPS_+/+ | 3.119209 | 5.318974 | 5.511379 | 4.755705 |
| 5142 | F | fl/+ | Saline | Saline_fl/+ | 0.032154 | 0.986221 | 0.17482 | 0.820221 |
| 5143 | F | fl/+ | LPS | LPS_fl/+ | 3.409996 | 5.939266 | 7.925144 | 6.260839 |
| 5144 | M | fl/+ | LPS | LPS_fl/+ | 3.053694 | 5.15844 | 5.185793 | 4.307207 |
| 5150 | M | fl/fl | Saline | Saline_fl/fl | 0.503309 | -0.16714 | 0.190874 | 0.247366 |
| 5151 | F | fl/fl | Saline | Saline_fl/fl | 0.435799 | 2.399986 | -2.34808 | 0.862123 |
| 5155 | F | fl/fl | LPS | LPS_fl/fl | 5.542513 | 3.656151 | 10.30392 | 5.951374 |
| 5156 | F | fl/fl | LPS | LPS_fl/fl | 1.185415 | 1.325159 | 6.136724 | 0.085371 |
| 5178 | M | fl/fl | LPS | LPS_fl/fl | 3.70586 | -1.91824 | -4.45667 | 3.646122 |
| 5179 | M | fl/fl | Saline | Saline_fl/fl | 0.237814 | -0.8165 | -0.50199 | -0.65733 |
| 5180 | F | +/+ | LPS | LPS_+/+ | 4.106718 | 2.47019 | 8.263233 | 5.526072 |
| 5181 | M | fl/+ | LPS | LPS_fl/+ | 0.727171 | -1.64733 | 0.483423 | -1.00379 |
| 5182 | F | fl/fl | Saline | Saline_fl/fl | 1.645779 | 1.877659 | 5.350048 | 2.339405 |
| 5183 | F | +/+ | Saline | Saline_+/+ | 0 | 0 | 0 | 0 |
| 5184 | M | fl/+ | Saline | Saline_fl/+ | -0.24056 | -1.32079 | 0.618589 | -0.60459 |
| 5185 | M | fl/+ | LPS | LPS_fl/+ | 3.655201 | 2.007597 | 6.807116 | 4.199827 |
| 5219 | M | +/+ | LPS | LPS_+/+ | 1.987885 | 0.421926 | 6.160135 | 2.30143 |
| 5220 | M | fl/+ | LPS | LPS_fl/+ | 2.92169 | -1.1942 | 4.289671 | 3.666336 |
| 5221 | M | fl/+ | saline | Saline_fl/+ | -0.71905 | -0.64039 | -1.99018 | -1.7966 |
| 5226 | M | +/+ | saline | Saline_+/+ | 0.171061 | 1.216911 | 1.175728 | -1.16616 |
| 5228 | M | fl/+ | LPS | LPS_fl/+ | 2.416086 | 0.179337 | 3.940735 | 1.329351 |
| 5230 | F | +/+ | LPS | LPS_+/+ | 3.009216 | 2.600588 | 7.751316 | 3.381695 |
| 5231 | F | fl/+ | LPS | LPS_fl/+ | 3.612831 | 1.844479 | 7.94656 | 5.573807 |
| 5232 | F | fl/fl | LPS | LPS_fl/fl | 4.731182 | 0.936483 | 11.13372 | 5.020716 |
| 5233 | M | fl/+ | LPS | LPS_fl/+ | 1.782167 | 3.476553 | 7.854252 | 6.546446 |
| 5234 | M | +/+ | LPS | LPS_+/+ | 2.178543 | -1.3736 | 3.85087 | 4.258171 |
| 5239 | F | fl/+ | LPS | LPS_fl/+ | 2.924772 | 1.599678 | 7.444796 | 3.845966 |
| 5240 | F | fl/+ | saline | Saline_fl/+ | -0.90178 | 0.090801 | -1.20319 | 0.654224 |
| 5241 | M | fl/+ | saline | Saline_fl/+ | -1.45335 | -1.13292 | -0.24692 | -0.73625 |
| 5242 | M | fl/fl | LPS | LPS_fl/fl | 3.171242 | 3.849104 | 9.063236 | 7.326674 |
| 5243 | M | +/+ | LPS | LPS_+/+ | 1.847694 | 0.522917 | 4.659294 | 1.41781 |
| 5244 | M | +/+ | LPS | LPS_+/+ | 3.395941 | 0.42944 | 4.004831 | 4.305792 |
| 5245 | F | +/+ | saline | Saline_+/+ | -0.17106 | -1.21691 | -1.17573 | 1.166161 |

| Kidney_STA | Kidney_CEI | Kidney_F3 | Kidney_RH | Kidney_TSI | Kidney_PLA | Kidney_PLA | Kidney_THI | Kidney_PLA |
|---|---|---|---|---|---|---|---|---|
| 0.413256 | 2.021477 | 0.080416 | 0.331924 | 0.284239 | 1.497627 | -0.62242 | 0.356752 | 0.265501 |
| 0.527964 | 1.92557 | 0.188105 | 0.024296 | -0.586 | 0.88802 | -0.10782 | 0.151585 | 0.230469 |
| 0.437485 | 1.753754 | 1.236629 | 0.441154 | 0.911757 | 0.681757 | -0.15478 | 0.302197 | 1.047491 |
| 0.607613 | 1.96587 | 0.121166 | 0.298685 | 0.514341 | 1.191359 | 0.311417 | 0.651281 | 0.943636 |
| 0.595253 | 1.341629 | 0.951141 | 0.908234 | 0.84901 | 0.949162 | -0.06425 | 0.456375 | 1.478249 |
| 2.44733 | 2.796463 | 0.692451 | 1.634478 | -1.34747 | 0.760042 | -1.52591 | 0.239466 | 3.820204 |
| 0 | 0 | 0 | 0 | 0 | 0 | 0 | 0 | 0 |
| 1.844664 | 3.285748 | -1.31165 | 1.466314 | -1.00763 | 1.530502 | -3.13278 | -0.60684 | 2.937141 |
| 0.636744 | 1.99366 | -0.65008 | 0.675442 | 0.320576 | 0.764032 | 0.116135 | 0.004852 | 0.527769 |
| 2.788998 | 3.863279 | 0.218891 | 2.568338 | -1.00061 | 1.499716 | -3.35971 | 0.354807 | 4.037895 |
| 1.71509 | 2.276833 | -0.27192 | 0.974154 | -1.10272 | 0.736139 | -2.21375 | -0.28644 | 2.055252 |
| -0.59893 | -1.90018 | 0.776716 | -1.18264 | -0.62959 | -1.27056 | -0.78229 | -0.6775 | -0.02508 |
| -8.0304 | -5.73028 | -3.68177 | -4.99992 | -3.34303 | -0.0589 | -0.82259 | 0.54055 | 1.302235 |
| 2.536985 | 1.104475 | 3.615705 | 1.847544 | -0.6757 | 0.597799 | -2.87723 | -0.05663 | 3.705435 |
| -1.20977 | -2.57913 | 0.759039 | -1.99416 | -2.72882 | -1.76833 | -7.81576 | -2.49621 | 0.206043 |
| -14.5811 | | 4.794668 | | -0.99694 | -0.53681 | -2.33842 | -0.111 | 4.170544 |
| -1.13468 | -1.35509 | 0.559435 | -1.48657 | -0.38909 | -0.60408 | -0.28019 | -0.09 | -0.02791 |
| 1.664862 | 1.434519 | -0.42929 | 1.401596 | -0.75887 | 0.342737 | -2.25386 | 0.613098 | 3.603765 |
| -0.77059 | -1.28374 | 1.015738 | -1.53384 | -0.50947 | -0.51683 | -0.24107 | 0.008781 | -0.29678 |
| 0.837128 | 3.497843 | 0.999531 | 0.526325 | 0.789858 | -0.49237 | -1.2045 | 0.645342 | 1.369164 |
| 0 | 0 | 0 | 0 | 0 | 0 | 0 | 0 | 0 |
| -1.27629 | -1.54936 | -0.42738 | -1.86932 | -0.77297 | -1.30205 | -1.83539 | -0.93914 | 0.438658 |
| 1.018963 | -0.02004 | -0.05059 | 0.504993 | -1.44002 | -0.08323 | -2.91546 | -0.12329 | 2.360746 |
| 1.645535 | 0.522837 | 1.875378 | 0.699249 | 0.022906 | -0.64488 | -0.57201 | -0.72055 | 0.095876 |
| 0.999838 | -0.49216 | 0.457924 | 0.310009 | -1.4505 | -1.12085 | -3.79955 | -0.45385 | 0.327506 |
| 0.622553 | -3.11135 | 0.532564 | 0.492699 | 0.89891 | -1.81646 | -0.93979 | 0.108649 | -0.41733 |
| -0.54798 | -0.63544 | 0.029171 | -0.44987 | -0.42888 | -0.2757 | 0.646337 | -0.45188 | -0.28478 |
| 3.289932 | 0.769547 | 0.579939 | 0.622349 | -0.78512 | 0.295347 | -1.82964 | -0.79089 | -0.82514 |
| 1.67395 | 3.62833 | 0.415586 | 1.481956 | -0.33812 | 0.579952 | -2.20989 | -0.28205 | 2.091361 |
| 3.01446 | 0.871822 | 0.033798 | 2.376608 | 0.067426 | -0.63482 | -3.90252 | 1.295205 | 3.686418 |
| 2.635809 | 0.932882 | 4.188005 | 2.649986 | 0.397351 | -0.14665 | -2.5264 | 0.510968 | 3.045106 |
| 1.580881 | 1.130157 | 1.096926 | 2.170589 | 0.389531 | 1.486853 | -3.4366 | 0.56845 | 3.567956 |
| 2.96698 | -0.57614 | 0.887262 | 0.952404 | -0.58189 | 0.511724 | -1.65419 | 0.324347 | 1.871715 |
| 2.153429 | 2.987381 | -0.66494 | 1.85956 | -1.06843 | 0.860699 | -3.36784 | -0.20248 | 4.049333 |
| -0.02145 | 1.948441 | -0.15493 | -0.92043 | 0.605399 | 0.784803 | 0.575549 | 0.252595 | 0.250375 |
| -0.6033 | -1.80206 | -0.05322 | -1.78205 | -0.67939 | -1.18503 | -1.78073 | -1.74419 | -1.63385 |
| 2.580528 | 1.213202 | 3.434364 | 2.800911 | 1.022819 | 0.897509 | -1.36253 | 1.135121 | 5.476956 |
| 1.773245 | -1.05686 | 2.009432 | 0.25028 | 0.263457 | 0.213734 | -2.97743 | -0.93685 | -0.14021 |
| 1.57234 | 0.857788 | 2.060564 | 1.976007 | 0.854777 | 1.203686 | -2.72952 | 0.158259 | 0.471681 |
| 0.547979 | 0.635445 | -0.02917 | 0.449869 | 0.428876 | 0.275696 | -0.64634 | 0.451882 | 0.284781 |

| Kidney_SY | Kidney_ET | Kidney_PT | Kidney_VW | Kidney_PR | Kidney_CD | Kidney_TF | Kidney_TF | Kidney_TF |
|---|---|---|---|---|---|---|---|---|
| -0.11799 | 1.098648 | 0.694284 | 1.839319 | -0.12833 | 2.508963 | 3.458002 | 3.195765 | 1.955925 |
| 1.959433 | -8.17899 | -0.53581 | 1.600193 | 0.03553 | 1.296522 | 2.543619 | 2.226002 | 1.719997 |
| 0.121876 | 0.686039 | 4.333078 | 3.686476 | 4.209932 | 5.971403 | 3.774944 | 4.065235 | 2.922657 |
| 1.082973 | 0.858147 | 0.950735 | 1.889278 | -0.18727 | 2.953714 | 2.787489 | 3.277943 | 1.456287 |
| 0.341206 | 0.305475 | 1.966753 | 2.189976 | 1.118441 | 3.441496 | 1.428135 | 1.67737 | 1.546968 |
| -0.89701 | 2.349737 | 4.079424 | 0.609631 | 6.458187 | 0.910482 | 3.702003 | 2.87142 | 2.429276 |
| 0 | 0 | 0 | 0 | 0 | 0 | 0 | 0 | 0 |
| -0.73948 | 2.298441 | 0.425575 | -1.97705 | -0.00015 | -0.04161 | -0.13684 | -2.58393 | -1.93966 |
| 0.628269 | 0.359652 | 0.200546 | 1.679304 | 0.970884 | 1.59737 | 2.513094 | 4.251692 | 2.255867 |
| -1.46037 | 3.317717 | 6.122181 | 1.4723 | 5.614824 | 4.442482 | 2.906609 | 0.04731 | 3.22703 |
| -0.74642 | 1.584549 | | | | | | | |
| -1.22812 | -0.25894 | -0.16411 | 0.204453 | -1.26776 | -0.38275 | -0.88262 | -0.64078 | -0.80944 |
| 0.527252 | 2.45459 | 1.138378 | 1.516037 | 0.329704 | 1.204403 | 0.497995 | -0.34127 | 0.016096 |
| -2.25225 | 3.190521 | 3.90173 | -0.86506 | 1.247932 | 0.766722 | -1.58732 | -3.83748 | -1.17157 |
| -5.69592 | 1.32585 | -1.71306 | -4.09555 | -0.35825 | 0.434275 | -1.30366 | | |
| -4.4718 | 2.617243 | 3.280508 | -2.55021 | 0.774609 | -2.31766 | -2.57133 | -5.99958 | -2.08574 |
| -1.04916 | 0.141844 | -0.63155 | -0.04057 | -0.42078 | -2.5321 | -0.9081 | -0.77295 | -0.99463 |
| -1.4415 | 3.56064 | 3.751837 | -1.46452 | -0.08399 | 1.264677 | -0.53786 | -3.10418 | -0.29178 |
| -1.07881 | -0.28477 | -0.8834 | -0.88159 | -0.61687 | -2.19066 | -0.67979 | -1.58451 | -1.19213 |
| 0.41547 | 1.470829 | 1.369678 | 1.801212 | 2.164997 | 2.024204 | 0.118845 | -0.81634 | -0.20222 |
| 0 | 0 | 0 | 0 | 0 | 0 | 0 | 0 | 0 |
| -0.33842 | -0.43918 | -0.04021 | -0.357 | -0.52871 | -0.82948 | -0.60813 | -1.43171 | -0.93785 |
| -1.10625 | 2.55423 | 2.079422 | -1.65622 | 0.354496 | 1.706991 | -2.07234 | -3.05839 | -1.22067 |
| -0.47919 | 1.183988 | -2.6475 | 1.687104 | -5.35966 | -0.24573 | -2.14947 | -2.71482 | -0.15692 |
| -0.88756 | 1.49433 | 1.865753 | 0.627851 | -3.59762 | -2.48517 | -1.8139 | -4.02765 | -0.09044 |
| -0.35119 | -1.02923 | -0.74141 | 2.674844 | -3.14289 | -1.7619 | -0.70552 | 0.655685 | 1.718821 |
| -0.03837 | 0.530597 | -0.19765 | 2.395616 | -1.86458 | 2.224889 | -0.83076 | 0 | -2.4355 |
| -0.84658 | 2.091851 | -0.66543 | 0.002284 | -4.8321 | -1.82591 | -0.48198 | -1.73387 | 1.337299 |
| 0.041113 | 2.112412 | 2.527403 | 1.921807 | -2.37757 | 0.326459 | 0.323912 | -0.48576 | 1.277599 |
| -0.69781 | 2.366939 | 3.83499 | 1.339618 | -3.65731 | -1.54928 | -0.11467 | -4.15382 | 1.06715 |
| -2.51882 | 1.801953 | 5.491459 | 1.915185 | -1.48845 | 2.793565 | 0.451406 | -1.38487 | 0.937572 |
| -0.87391 | 1.074521 | 1.943472 | -0.03186 | -2.58696 | 1.238059 | -0.3305 | -3.24963 | -1.06957 |
| -0.22886 | 1.704713 | 0.533918 | 1.328115 | -2.39606 | -0.37577 | -2.08475 | -1.78691 | 1.101501 |
| -0.97206 | 1.985732 | -0.06033 | 1.592652 | -2.84822 | -0.10481 | -2.57612 | -4.26134 | 0.043417 |
| 0.700676 | -0.58297 | -0.92126 | 3.781127 | -3.35996 | -0.19146 | 0.832487 | -0.97866 | 0.974722 |
| -0.4769 | -1.04158 | -0.83823 | 2.225814 | -4.84028 | -3.57465 | -0.07711 | -1.67291 | 0.172991 |
| -1.22367 | 2.426576 | 5.174128 | 1.200456 | 0.583572 | 2.201949 | -1.41494 | -3.30374 | 1.462576 |
| -1.80513 | 1.116328 | -0.3921 | 2.10971 | -3.01001 | -2.8553 | -0.71996 | -2.43234 | -0.04452 |
| 0.310436 | 2.666726 | 1.578192 | 0.854098 | -2.15079 | -1.45026 | -0.03341 | -1.07795 | 0.282244 |
| 0.038368 | -0.5306 | 0.197651 | -2.39562 | -4.04878 | -2.22489 | 0.830765 | | 2.435503 |

| Kidney_SEI | Kidney_C5/ | Kidney_VE( | Kidney_SEI | Kidney_ST/ | Kidney_OA | Kidney_OA | Kidney_OA | Kidney_OA |
|---|---|---|---|---|---|---|---|---|
| 1.269251 | 2.162123 | 2.916624 | -0.3482 | -0.04277 | 0.438843 | 0.728905 | 0.175632 | 1.938715 |
| 2.356091 | 2.26005 | 2.520353 | -0.42876 | 0.495373 | 0.359022 | 1.297119 | 0.542112 | 1.274065 |
| 4.638296 | 4.849234 | 6.824125 | 3.507164 | 3.227791 | 1.718117 | 4.367758 | 3.350674 | 2.920351 |
| 2.292814 | 3.317087 | 2.906809 | 0.635353 | 0.742064 | 1.576694 | 2.022707 | 1.173035 | 3.626066 |
| 1.92415 | 4.883738 | 3.831381 | 2.523823 | 1.112734 | 1.614653 | 2.53714 | 2.132542 | 5.670525 |
| 6.434731 | 5.538319 | 4.228794 | 5.937637 | 4.487949 | 2.609201 | 4.628534 | 4.967222 | 6.022429 |
| 0 | 0 | 0 | 0 | 0 | 0 | 0 | 0 | 0 |
| 3.86376 | 1.861258 | 0.940973 | 2.010941 | 1.075802 | 1.576622 | 1.434242 | 2.525763 | 4.005497 |
| 0.553436 | 3.104691 | 4.352018 | 0.259508 | 1.962849 | 0.59874 | 1.259262 | 1.04278 | 0.943504 |
| 8.214859 | 3.953663 | 1.94389 | 6.187994 | 5.557039 | 6.383612 | 4.791891 | 4.371147 | 6.390137 |
| 1.196426 | 0.423565 | 0.796825 | -0.2116 | -6.09826 | 0.154839 | -1.20528 | -0.54749 | -1.73708 |
| 1.916176 | 0.739208 | 1.395432 | -0.33458 | 0.500317 | 0.705044 | -0.6075 | 1.807709 | -0.24557 |
| 7.709177 | 1.608521 | -0.60858 | 3.019424 | 2.165882 | 5.735958 | 1.629871 | 3.107082 | 2.212635 |
| 7.049353 | 1.23769 | 0.182196 | -3.39279 | 1.609356 | 1.747765 | 1.36553 | 1.826096 | 1.947975 |
| 7.669436 | 1.219425 | -1.86286 | 4.180803 | 2.304087 | 5.505375 | -0.61533 | 2.6075 | 3.320885 |
| 1.267834 | 2.323545 | 0.85461 | -0.38218 | -0.18592 | 1.742373 | -0.44146 | -1.0801 | 0.231892 |
| 6.260258 | 1.123894 | -0.78649 | 4.047905 | 3.225163 | 5.492216 | 0.475216 | 1.950716 | 2.133354 |
| -0.01217 | 0.655165 | 0.154879 | -1.27036 | 0.022087 | 3.634647 | 1.562593 | 0.237196 | 2.116287 |
| 5.970818 | 1.845266 | 0.377245 | 2.195366 | -0.25444 | 0.711288 | -0.44759 | -0.34711 | -0.30494 |
| 0 | 0 | 0 | 0 | 0 | 0 | 0 | 0 | 0 |
| 0.915207 | 2.144444 | 0.605507 | 0.136076 | -1.49093 | 0.431236 | -0.90766 | -0.03354 | 1.499664 |
| 5.632404 | 1.11552 | -1.06618 | 5.085316 | 0.924652 | 3.139143 | 2.028492 | 3.002596 | 1.392542 |
| 2.587812 | -0.03721 | -2.16401 | -1.16507 | 1.469695 | 2.440596 | 1.884672 | 2.431425 | 2.551105 |
| 1.343655 | -2.73201 | -2.69841 | -0.26397 | 1.094466 | 2.602821 | 1.12351 | 2.064598 | 1.706432 |
| -0.84256 | -2.27068 | -1.46252 | -3.40099 | -0.80486 | -0.56341 | -0.37547 | -0.49549 | -0.21651 |
| 0.171539 | 0.615724 | -1.04755 | -0.04829 | -0.2712 | 0.144461 | -0.05309 | -0.07633 | 0.049429 |
| 1.307526 | 0.410295 | -0.87065 | 0.052138 | 1.142447 | 2.72049 | 1.513788 | 1.701529 | 1.842653 |
| 4.461082 | 1.181312 | -0.35057 | -0.30071 | 0.887322 | 2.322741 | 1.761278 | 2.062458 | 2.148258 |
| 1.58992 | -0.78314 | -0.56509 | 1.091979 | 2.889592 | 5.795471 | 2.548849 | 3.72097 | 3.372805 |
| 3.678185 | -0.20071 | -1.21354 | 1.24611 | 2.784446 | 7.461332 | 1.787138 | 1.969181 | 2.891258 |
| 2.810345 | -0.71795 | -1.2754 | 1.226413 | 0.980828 | 3.429274 | 1.595837 | | 1.764128 |
| 0.860819 | 0.227773 | -1.69134 | 1.240932 | 1.250468 | 2.148212 | 1.247355 | 1.838499 | 2.111542 |
| 3.101774 | -1.9875 | -1.86495 | 2.718581 | 0.92615 | 2.881189 | 1.028048 | 1.877747 | 0.75038 |
| -0.76259 | -1.4467 | -0.65012 | -0.76153 | 0.389722 | 0.247622 | 0.554354 | -0.15893 | -0.08505 |
| -1.14612 | -0.38131 | -0.20967 | -2.96068 | -1.21771 | -1.32011 | -1.11728 | -1.36878 | -1.39089 |
| 6.09882 | -1.14652 | -1.44523 | 1.499356 | 1.944638 | 4.077269 | 2.20126 | 2.701912 | 3.396797 |
| 3.880249 | 1.361508 | -2.19552 | 1.785201 | 1.571364 | 3.06757 | 2.263132 | 2.075977 | 1.721107 |
| 3.445757 | -0.09212 | -0.28525 | 0.970349 | 1.315553 | 2.821898 | 1.927843 | 1.788464 | 2.3617 |
| -0.17154 | -0.61572 | 1.047545 | 0.048293 | 0.271195 | -0.14446 | 0.053089 | 0.076332 | -0.04943 |

| Kidney_FO! | Kidney_MX | Kidney_IL1 | Kidney_IRF | Kidney_IRF | Kidney_IRF | Kidney_CX( | Kidney_CX( | Kidney_IFIH |
|---|---|---|---|---|---|---|---|---|
| 0.298452 | 2.570082 | 0.182848 | 1.848822 | 0.717697 | 1.094612 | 0.873583 | 2.171083 | 1.717949 |
| -0.26877 | 2.359655 | -1.19579 | 0.754061 | 0.791164 | 0.293289 | 0.336073 | 0.075163 | 0.396484 |
| 3.638697 | 4.797007 | 2.014683 | 3.275902 | 3.004833 | 3.496931 | 3.818644 | 2.415205 | 3.360935 |
| 1.394842 | 3.061047 | -0.71922 | 2.217489 | 1.976543 | 0.476328 | 3.257114 | 1.386934 | 2.146269 |
| 0.787605 | 3.910963 | 0.099592 | 0.536741 | 1.632708 | 0.59618 | 3.017843 | 3.039793 | 0.648031 |
| 1.183563 | 8.559776 | 2.464842 | 3.948492 | 7.014448 | 4.512053 | 8.357834 | 9.506042 | 3.824659 |
| 0 | 0 | 0 | 0 | 0 | 0 | 0 | 0 | 0 |
| -2.47495 | 3.735092 | -1.47817 | 0.9182 | 2.643463 | -0.00492 | 3.696739 | 6.470341 | 0.417627 |
| -0.64635 | 3.294722 | 0.534691 | 1.770451 | 1.444733 | 1.206518 | 1.67329 | 1.97229 | 1.157633 |
| 2.32346 | 9.263176 | 1.494062 | 5.206284 | 6.094528 | 4.644545 | 8.072556 | 10.26512 | 4.492039 |
| 3.735765 | 0.153839 | 0.812017 | 0.12063 | -0.90881 | -1.10371 | 3.408899 | 1.55999 | 0.275879 |
| 4.177067 | 0.631598 | 1.495907 | 0.446802 | 0.479361 | -0.20021 | 1.144695 | 1.41713 | 1.789824 |
| 7.98543 | 8.994232 | 1.105158 | 2.099625 | 3.183851 | 2.381397 | 7.032499 | 9.987009 | 3.737276 |
| 3.070585 | 2.756538 | 0.649282 | 0.531588 | 4.016499 | -1.63957 | 2.743534 | 7.156952 | 2.68771 |
| 5.682774 | 5.84186 | 2.092665 | 2.599316 | 4.008234 | 4.788557 | 10.50824 | 8.014919 | 3.243635 |
| 4.288481 | 1.537804 | 0.499994 | -1.2073 | -1.03751 | 0.57888 | 0.492765 | 0.509018 | 1.821304 |
| 4.78582 | 4.371946 | 2.326273 | 1.956059 | 4.826448 | 0.465773 | 5.129948 | 8.037325 | 4.469013 |
| 3.227028 | 1.508022 | 2.303328 | -0.92619 | 4.620094 | -0.83641 | 0.373892 | 5.700432 | 2.049212 |
| 5.460192 | -0.11485 | 1.222652 | 0.161264 | -0.31966 | -0.28451 | 1.997168 | 1.938848 | 1.04418 |
| 0 | 0 | 0 | 0 | 0 | 0 | 0 | 0 | 0 |
| 4.681795 | -0.82122 | 1.327883 | -0.53462 | 0.619228 | -0.07347 | -0.90089 | -0.35289 | 2.574493 |
| 3.60442 | 4.424936 | -0.04522 | 1.147696 | 3.008404 | 1.339462 | 6.524305 | 5.750174 | 3.148706 |
| 2.07525 | 2.336477 | -0.42202 | 0.516919 | 3.501933 | -0.5087 | 1.608172 | 4.923328 | -0.42685 |
| 1.198663 | 3.003244 | -0.84838 | 0.836532 | 4.036094 | 0.869154 | 2.664727 | 3.648567 | 0.460917 |
| 1.432927 | -0.18129 | -0.45327 | 0.467515 | -0.39287 | 0.244425 | -2.8757 | -0.26134 | -1.24807 |
| -0.66936 | -0.64479 | -0.92698 | 0.33814 | 0.620673 | -0.69125 | 0.574126 | 0.205803 | -1.31493 |
| -0.30717 | 1.886244 | -0.41181 | 0.574832 | 4.577336 | 0.726807 | 0.230178 | 3.636652 | 0.276027 |
| 1.411847 | 1.972559 | -0.88571 | 0.717629 | 3.441855 | -0.60582 | 2.924882 | 5.427254 | -0.81272 |
| 2.214725 | 5.972178 | -0.41761 | 2.031661 | 3.365907 | 0.772184 | 5.813878 | 8.329351 | 1.397849 |
| 5.031491 | 7.573597 | -0.88304 | 3.416818 | 2.340907 | 2.787169 | 5.520492 | 7.383717 | 2.325225 |
| 1.371409 | 3.128849 | -0.45276 | -0.25393 | 2.171704 | 1.220955 | 3.524102 | 5.671089 | 0.576745 |
| 0.224675 | 1.799675 | -0.48046 | 0.342147 | 3.214063 | 0.299639 | 1.966641 | 5.334438 | -0.42993 |
| 1.723029 | 5.097507 | -0.4262 | 2.197806 | 3.795993 | 1.014641 | 2.936933 | 6.163729 | 0.746707 |
| -0.12767 | 0.920774 | 0.298008 | 1.741147 | 0.841353 | 1.189089 | -1.91039 | -0.07308 | 0.714053 |
| -1.23394 | -0.08789 | 0.315165 | 0.819898 | 0.058343 | 0.212763 | -2.59248 | 1.636885 | 0.076398 |
| 3.311284 | 5.439823 | 0.925106 | 2.303782 | 2.931239 | 2.740929 | 6.545203 | 8.281506 | 1.72092 |
| 1.521827 | 3.04982 | 0.056765 | 0.89849 | 3.842406 | -0.01759 | 1.230109 | 4.969284 | 0.338748 |
| 1.62654 | 3.024372 | 0.028542 | 1.180022 | 3.638764 | 0.636646 | 2.965178 | 5.795015 | 1.233079 |
| 0.669356 | 0.644794 | 0.926979 | 1.777587 | -0.62067 | 0.691254 | -0.57413 | -0.2058 | 1.314927 |

| Kidney_CL | Kidney_HA | Kidney_LC | Kidney_DD | Kidney_KL | Lung_ICAM | Lung_SELE | Lung_IL6 | Lung_SOCS |
|---|---|---|---|---|---|---|---|---|
| 1.757071 | -2.03746 | 2.378557 | 2.361448 | 0.489429 | -0.68714 | -1.50876 | 0.204306 | -0.82252 |
| 0.510906 | -1.69005 | 3.423845 | 1.4562 | -0.36991 | -0.61917 | -1.44511 | 0.016329 | 0.060081 |
| 3.381325 | 4.49305 | 4.824846 | 3.931494 | 3.627354 | -0.75062 | -0.95399 | -0.16222 | 0.740208 |
| 2.69409 | -0.8035 | 2.406004 | 2.288696 | 0.996758 | 0.479399 | -1.80853 | -0.02301 | -0.50557 |
| 0.074099 | 1.288136 | 3.457289 | 1.737391 | 0.919004 | -0.20292 | -1.95278 | -1.34961 | 0.029417 |
| 6.287144 | 9.198212 | 15.01878 | 4.168976 | 1.542328 | 1.243162 | 3.713293 | 7.365181 | 3.36043 |
| 0 | 0 | 0 | 0 | 0 | 0 | 0 | 0 | 0 |
| 2.817207 | 3.914713 | 13.49524 | 1.413816 | -1.16079 | 3.510906 | 5.182539 | 6.67178 | 2.761604 |
| 2.986734 | -2.44232 | 3.22472 | 2.091066 | 0.417315 | 0.028225 | -0.97596 | -0.30757 | -0.51032 |
| 6.486385 | 5.242096 | 15.86073 | 4.604225 | 1.627916 | 1.441753 | 3.319386 | 8.352955 | 4.011662 |
| | | | | | 2.728823 | 4.359463 | 6.157642 | 2.626589 |
| 1.065985 | 1.380398 | -1.53243 | -0.7556 | -1.41761 | -0.06133 | 1.100473 | -1.21055 | 0.520287 |
| 1.953306 | -0.1572 | -0.41642 | -0.22506 | -0.22244 | 0.20665 | 0.082458 | -0.50044 | 0.482883 |
| 2.370632 | 3.470436 | 11.41995 | 3.852043 | -0.2945 | 2.86767 | 4.216791 | 8.018751 | 5.196012 |
| 3.380064 | 5.458778 | 8.194359 | 0.94537 | -0.79039 | 4.926825 | 0.788792 | 2.818794 | |
| 3.020029 | 7.453047 | 10.74889 | 2.080128 | -0.28526 | 1.012299 | 5.295628 | 9.856358 | 4.30744 |
| 1.094109 | 2.160469 | -2.04527 | 0.328133 | -0.71921 | -0.07698 | 1.056026 | -0.08562 | 0.484241 |
| 3.013639 | 5.304905 | 12.65423 | 2.642202 | 0.482241 | 2.422859 | 4.977522 | 7.746126 | 3.591244 |
| 0.470886 | 3.288378 | 2.888155 | 1.313278 | -1.34259 | 0.899441 | 0.231697 | 0.224194 | 0.563925 |
| 2.592793 | 3.525087 | 6.887873 | -0.29814 | 0.084646 | 0.066555 | 1.91721 | 0.345264 | 1.708279 |
| 0 | 0 | 0 | 0 | 0 | 0 | 0 | 0 | 0 |
| 1.61105 | 2.937048 | 0.379044 | -0.24475 | 0.353203 | 1.234756 | 2.85001 | 0.929882 | 1.795898 |
| 3.885937 | 7.094698 | 11.25391 | 0.939964 | 0.217281 | 1.234756 | 2.85001 | 6.68948 | 1.795898 |
| -1.66268 | 5.352796 | 7.832028 | -0.22812 | -0.86048 | 2.021196 | 5.683111 | 7.189274 | 2.613345 |
| 0.017148 | 5.793424 | 8.064004 | 1.280437 | -1.09461 | 1.168585 | 5.206066 | 6.595139 | 3.178481 |
| -2.36914 | 1.565579 | -2.99107 | 0.638259 | -0.51839 | 0.225922 | 0.753672 | 1.150141 | 1.706304 |
| -2.10899 | -0.15383 | -0.9176 | 0.489734 | -0.50726 | -0.03896 | 0.5905 | -0.09459 | 0.495494 |
| 0.768128 | 5.223773 | 8.533387 | -0.5885 | -1.14078 | 2.488873 | 6.650274 | 5.761091 | 1.842364 |
| 1.439263 | 4.432321 | 9.257689 | -1.22675 | -1.37018 | 1.65662 | 4.327301 | 6.405022 | 1.99615 |
| 0.597407 | 1.845489 | 8.845271 | 1.616508 | 0.455905 | 1.187998 | 4.331285 | 8.388226 | 3.499438 |
| 1.726069 | 0.29263 | 8.660691 | 2.37637 | 0.38309 | 1.808041 | 5.894041 | 12.31264 | 5.029502 |
| 1.77381 | 6.351104 | 8.796184 | -0.03313 | -0.67879 | 1.299618 | 5.176006 | 6.385067 | 2.273339 |
| 1.170096 | 6.183835 | 9.377906 | 0.200683 | -1.94986 | 2.45756 | 5.697594 | 6.259157 | 2.863194 |
| 2.90722 | 5.376613 | 11.00156 | 1.115617 | -1.26416 | 2.502142 | 5.192753 | 7.756954 | 4.494531 |
| -0.60467 | -0.9832 | -0.38228 | 1.748078 | 0.461223 | -0.63189 | 0.300316 | -0.74773 | 0.694575 |
| -0.5709 | 2.784495 | -0.28403 | -0.67322 | -1.13059 | -0.25553 | -0.03675 | -0.1955 | 0.29742 |
| 2.857985 | 7.611072 | 10.64488 | 0.531438 | -2.00253 | 0.459423 | 3.433197 | 9.834047 | 4.056365 |
| 0.962409 | 6.805753 | 9.181832 | 0.833747 | -1.91166 | 1.812653 | 4.563999 | 4.568018 | 2.525073 |
| 1.69774 | 7.525397 | 9.544759 | 0.046445 | -0.73306 | 1.727343 | 5.354656 | 7.029053 | 2.612523 |
| 2.108994 | 0.153829 | 0.917596 | 0.588504 | 0.507257 | -1.188 | -0.5905 | 0.094593 | -0.49549 |

| Lung_STAT | Lung_CEMI | Lung_F3 | Lung_RHO( | Lung_TSP1 | Lung_PLAT | Lung_PLAU | Lung_THBD | Lung_PLAU |
|---|---|---|---|---|---|---|---|---|
| -0.51303 | -1.04048 | 0.011799 | -1.21777 | -0.86705 | -0.39313 | -0.03259 | -0.532 | -0.67819 |
| 0.784105 | -0.44266 | 0.007669 | 1.054527 | -0.26782 | 0.113747 | -0.08546 | -0.14759 | -0.14455 |
| -0.17619 | 0.129221 | 0.958202 | -0.53259 | 0.185009 | 0.446455 | -0.70843 | -0.18451 | 0.135521 |
| 0.236818 | -0.58404 | -0.0251 | 0.325029 | 0.016483 | -0.2438 | -0.64414 | -0.79063 | -0.82799 |
| 0.321852 | -0.96663 | -0.68232 | -0.18958 | -0.80285 | 0.412439 | -1.09 | 0.554453 | 0.315174 |
| 1.936457 | 4.427279 | 0.970203 | 1.82408 | -0.2372 | 2.799347 | -2.94683 | -2.78369 | 2.65287 |
| 0 | 0 | 0 | 0 | 0 | 0 | 0 | 0 | 0 |
| 0.369402 | 4.250795 | -0.0302 | 0.484602 | -0.69469 | 2.680826 | -0.7534 | -2.63347 | 1.391054 |
| -0.84673 | -0.3743 | -0.18903 | -0.30423 | -0.18911 | -0.73497 | -1.53819 | -0.80877 | -1.14342 |
| 1.290501 | 4.005737 | 0.236967 | 1.901178 | -0.54757 | 2.715126 | -2.12717 | -2.9168 | 1.748262 |
| 0.447203 | 3.818916 | 0.1486 | 0.880281 | -0.63651 | 3.173239 | -3.30982 | -2.26648 | 0.984423 |
| -0.80672 | -1.01861 | -0.81252 | -0.73119 | -0.11635 | -0.95758 | -1.08069 | -0.45581 | -0.70846 |
| 0.115849 | -0.00041 | -0.65718 | -0.08854 | 0.888885 | -0.24652 | -0.299 | 0.437859 | 0.066486 |
| 2.087835 | 3.928976 | 0.053917 | 2.596869 | 1.545248 | 2.253874 | -1.15507 | -2.79553 | 2.920994 |
| | | -1.73866 | -1.09331 | 0.619045 | 1.056835 | -1.42034 | -2.04693 | 0.288307 |
| 1.539265 | 3.942833 | 0.764925 | 1.803627 | 2.356943 | 1.785736 | -1.68516 | -2.49492 | 3.260075 |
| 0.021162 | -0.76459 | -0.5348 | -0.30973 | 0.274252 | 0.016911 | 0.029358 | 0.142355 | -0.44252 |
| 0.994858 | 5.055891 | -0.3698 | 0.766964 | -0.00966 | 2.686348 | -0.66326 | -2.38802 | 2.214058 |
| 0.819851 | 0.100994 | -0.4645 | 0.338408 | 0.177555 | 0.102636 | 1.379364 | 0.879654 | 0.512617 |
| 1.015961 | 1.224075 | -0.61025 | 0.775684 | 1.436459 | 0.741722 | -2.04996 | -0.44756 | 0.57234 |
| 0 | 0 | 0 | 0 | 0 | 0 | 0 | 0 | 0 |
| 0.139164 | 0.663322 | -0.22843 | 0.219658 | 0.760326 | 0.089741 | 0.375034 | 0.125143 | -0.09308 |
| 1.283007 | 3.89146 | -0.43203 | 0.89542 | -0.28599 | 3.124443 | -0.45738 | -2.06985 | 1.789692 |
| 1.511287 | 3.666103 | -0.21962 | 1.477282 | -0.24882 | 2.578872 | -0.211 | -1.07567 | 1.111712 |
| 1.222436 | 3.869551 | 0.439362 | 1.805384 | -0.0421 | 2.890922 | -1.40895 | -1.95249 | 1.999269 |
| 0.314351 | 1.369469 | 0.30422 | 0.811558 | 1.325111 | -0.70423 | 0.082566 | -0.05174 | 0.887848 |
| 0.645082 | -0.97948 | 1.028904 | 0.863958 | 0.147179 | -0.37852 | 0.347781 | 1.0622 | -0.04268 |
| 1.269015 | 4.406046 | 0.910276 | 1.751255 | 0.810862 | 3.994925 | -0.79327 | -1.11824 | 2.205053 |
| 0.800366 | 3.174648 | 0.012178 | 1.155012 | -0.68688 | 2.854224 | -1.17884 | -2.06902 | 1.188776 |
| 1.226539 | 4.594278 | 0.277498 | 1.411217 | -0.12139 | 3.884271 | -1.63865 | -1.86961 | 2.480869 |
| 2.429957 | 5.725391 | 1.970886 | 3.853239 | 2.902803 | 3.686334 | -0.79926 | -1.44224 | 4.056137 |
| 0.230706 | 2.860468 | -0.04477 | 0.207571 | -0.7505 | 3.482285 | -1.71254 | -2.15991 | 2.094957 |
| 1.454148 | 3.307335 | 0.91802 | 0.823383 | -0.07635 | 3.310376 | 0.210916 | -1.16567 | 2.046082 |
| 2.03194 | 5.684864 | 2.053936 | 1.911196 | 0.419283 | 4.600175 | -0.17948 | -1.35328 | 3.138992 |
| 0.550071 | -0.35944 | 0.975042 | 0.694309 | 0.330837 | 2.14956 | 0.420096 | 0.998099 | 0.788137 |
| 0.33895 | -0.16775 | 0.546862 | -0.24532 | -0.45974 | 1.709382 | -0.78238 | 0.308431 | 0.352381 |
| 2.325112 | 4.955278 | 2.282408 | 1.944246 | 1.046949 | 3.415317 | -1.08928 | -2.77069 | 3.434959 |
| 1.229407 | 4.189312 | 0.338455 | 0.6115 | 0.811377 | 3.015821 | -1.3626 | -0.98838 | 1.49122 |
| 0.646678 | 4.7164 | -0.27512 | 0.640079 | 0.725328 | 2.979566 | -1.84352 | -1.32501 | 1.752592 |
| -0.64508 | 0.979475 | -1.0289 | -0.86396 | -0.14718 | 0.378522 | -0.34778 | -1.0622 | 0.042677 |

| Lung_PTGS | Lung_VWF | Lung_PROC | Lung_CD44 | Lung_TFPIA | Lung_TFPIB | Lung_TFPIC | Lung_SELP | Lung_C5AR |
|---|---|---|---|---|---|---|---|---|
| 0.066343 | -0.30875 | 0.662714 | 0.18799 | 0.573853 | 0.48921 | 0.612982 | -1.6524 | -1.38344 |
| 0.689711 | 1.034357 | 0.711302 | -0.14554 | 0.780048 | 0.354557 | 0.657555 | 0.658291 | -1.05216 |
| 0.869833 | -0.2571 | 0.340195 | 0.940752 | 0.113155 | 0.201492 | 0.004425 | 0.581978 | 2.289011 |
| 1.391691 | 1.093735 | 0.445307 | -0.4509 | 0.831066 | 0.596565 | 0.501402 | 1.984949 | -0.10718 |
| 0.154221 | 0.700768 | 0.042168 | -0.6853 | 0.691109 | 0.63402 | 0.054382 | -0.11165 | -0.21742 |
| 1.538973 | -2.76149 | 3.751825 | 0.579416 | 0.123928 | -1.81126 | 0.433792 | 6.628151 | 2.925356 |
| 0 | 0 | 0 | 0 | 0 | 0 | 0 | 0 | 0 |
| 3.15675 | -1.7523 | 2.90612 | -0.29273 | -0.42763 | -2.22748 | -0.37578 | 5.452446 | 2.902052 |
| 0.92651 | 0.321123 | -0.11281 | 0.866581 | -0.08209 | -0.21033 | -0.33698 | -0.06801 | -0.00963 |
| 1.457428 | -2.60995 | 2.54472 | -0.01081 | -0.38913 | -3.26611 | -0.06496 | 5.644602 | 2.759741 |
| 1.835438 | -2.08575 | 2.496641 | -0.16373 | -0.21961 | -1.62371 | -0.40316 | 4.130268 | 3.24749 |
| 0.633482 | -0.05759 | | -0.1007 | 0.299458 | 0.226797 | -0.01024 | 0.967531 | 0.014708 |
| 0.086422 | -0.25659 | | 0.277033 | 0.457756 | 0.211803 | -0.32771 | 0.867434 | -0.63632 |
| 1.46615 | -1.54161 | | 0.80056 | -0.79005 | -2.34153 | -0.56349 | 5.791674 | 3.643564 |
| 0.265606 | -1.62831 | | -0.63953 | -0.05073 | -0.71282 | -0.62263 | 4.626442 | 0.779114 |
| 0.617413 | -2.61956 | | 0.994555 | -0.78767 | -2.91483 | -0.86444 | 6.7316 | 3.060442 |
| -0.02176 | -0.24668 | | -0.39298 | 0.081514 | -0.03861 | -0.03226 | 1.561321 | -0.80377 |
| 2.853832 | -0.87117 | | 0.114199 | -0.62868 | -2.49693 | -0.70733 | 4.553501 | 2.852236 |
| 0.98444 | 0.91699 | | -0.68532 | -0.29265 | -0.07059 | -0.28156 | -0.1307 | 1.064346 |
| 0.306705 | 1.321371 | | -0.45333 | 0.425938 | -0.31871 | 0.198492 | 2.407709 | 1.709707 |
| 0 | 0 | | 0 | 0 | 0 | 0 | 0 | 0 |
| 1.286263 | 0.190756 | | 0.813168 | -0.35707 | -0.31335 | -0.44106 | -0.05883 | 0.255848 |
| 1.777416 | -1.12015 | | 0.626358 | -0.16112 | -1.7985 | -0.17436 | 4.780209 | 3.710514 |
| 2.3388 | -1.54481 | 3.084693 | 0.238804 | 0.204016 | -1.30098 | -0.06867 | 5.100382 | 0.479641 |
| 1.851198 | -1.5233 | 3.186679 | 1.013945 | -0.69988 | -1.62287 | -0.47195 | 5.456518 | 1.903774 |
| 0.838068 | 0.602166 | 1.357953 | 1.737534 | 0.383406 | -0.34869 | 0.607638 | -0.32473 | -2.76035 |
| 1.038643 | 1.188611 | 0.673285 | 1.737255 | 1.265875 | 0.477373 | 1.089961 | 0.681263 | 0.663336 |
| 1.650146 | -1.03678 | 3.745984 | 1.605932 | -0.50478 | -0.82394 | 0.073897 | 4.622591 | 3.364872 |
| 1.370916 | -2.17833 | 3.315976 | 0.692583 | 0.598614 | -1.15001 | 0.099145 | 5.516539 | -0.19826 |
| 1.981218 | -1.32744 | 5.188476 | 1.928625 | -0.201 | -3.08954 | -0.78011 | 6.340336 | 2.448786 |
| 3.311722 | -1.62839 | 4.207038 | -1.38507 | -2.35964 | -2.46638 | 0.27665 | 6.685545 | 0.655064 |
| 1.619427 | -2.50147 | 4.96931 | -0.71838 | 0.096853 | -2.95249 | -2.59065 | 7.287344 | 2.789034 |
| 2.003138 | -1.123 | 3.409362 | 1.087776 | -0.41751 | -1.21717 | 0.542808 | 5.461582 | 1.389004 |
| 1.914873 | -2.04486 | 4.82001 | 1.748438 | 0.249067 | -2.63637 | -0.16136 | 7.246729 | 1.42128 |
| -0.19331 | 0.729316 | 2.23033 | 1.731071 | 1.096188 | 0.420609 | -0.14708 | 0.41856 | 0.959774 |
| -0.22695 | 0.614524 | 0.638238 | 0.477419 | 0.951208 | 0.466568 | 0.107013 | 0.47047 | 0.335083 |
| 1.522041 | -3.31107 | 3.999705 | 1.450468 | -0.28625 | -2.10018 | 0.372663 | 6.105783 | -1.34118 |
| 1.260208 | -1.17589 | 2.511163 | 0.050514 | 0.646271 | -0.62008 | -0.53397 | 4.304363 | -0.04172 |
| 0.708193 | -2.18176 | 2.250936 | -0.50635 | -1.27155 | -0.90175 | -0.65889 | 2.587709 | 1.907967 |
| -1.03864 | -1.18861 | -0.67329 | -1.73726 | -1.26587 | -0.47737 | -1.08996 | -0.68126 | -0.66334 |

| Lung_VEGF | Lung_SERP | Lung_STAT | Lung_OASL | Lung_OAS2 | Lung_OASL | Lung_OAS3 | Lung_FOS | Lung_MX1 |
|---|---|---|---|---|---|---|---|---|
| -2.25532 | -1.20062 | -0.29811 | -0.63653 | -0.52837 | -0.07624 | -1.23551 | -0.66652 | 0.014009 |
| 0.20521 | -0.99723 | 0.103857 | 0.615911 | 1.17403 | 0.777195 | 0.796541 | -0.08331 | 1.151705 |
| -1.33574 | -0.55761 | -0.30867 | 0.075024 | 0.500589 | 0.591736 | 0.716629 | -0.2593 | -0.58858 |
| 1.110395 | -0.67363 | 0.435986 | 0.02564 | 0.130119 | 0.411043 | -0.25927 | -0.05845 | 0.346544 |
| -0.28484 | -0.13795 | 0.216385 | 1.125238 | 1.368671 | 0.799213 | 0.668318 | -0.51401 | -0.27088 |
| -1.10476 | 4.381271 | 1.665787 | 3.180634 | 1.505859 | 2.074116 | 2.541866 | 0.465672 | 3.198833 |
| 0 | 0 | 0 | 0 | 0 | 0 | 0 | 0 | 0 |
| -2.73628 | 4.301022 | 1.347277 | 3.198051 | 1.966993 | 2.070837 | 3.440439 | -0.67916 | 2.390028 |
| -1.66442 | -0.73884 | -0.3913 | -0.74618 | -0.49572 | -0.32963 | -1.12264 | -0.34158 | -0.53921 |
| -2.4743 | 2.690945 | 2.140287 | 4.505756 | 2.506346 | 2.471949 | 3.095417 | 0.876358 | 4.970711 |
| -2.33469 | 3.471811 | 1.356583 | 3.468866 | 2.002218 | 2.430719 | 3.418871 | -1.60174 | 2.515692 |
| -0.58231 | 0.039549 | 0.058992 | 0.373892 | 0.836727 | 0.076809 | 0.440111 | 1.750746 | -0.42766 |
| 0.04497 | 0.743774 | 0.342148 | 0.187078 | 0.712738 | 0.475697 | 0.29772 | 1.143021 | -0.40812 |
| -0.63103 | 3.491032 | 2.515886 | 6.545265 | 3.185061 | 2.822908 | 3.971767 | 3.44165 | 6.321167 |
| -2.32185 | 4.391897 | 1.497362 | 3.184113 | 2.427361 | 2.461298 | 3.240459 | 1.929714 | 2.564716 |
| -1.92056 | 4.405962 | 2.11554 | 7.036224 | 2.944748 | 2.358932 | 4.340538 | 3.716238 | 6.000229 |
| -0.43391 | 0.317064 | 0.038643 | 0.188856 | 0.500652 | 0.334988 | -0.63326 | 1.544653 | 0.062103 |
| -0.53659 | 4.571581 | 1.921352 | 4.246325 | 2.05529 | 2.932711 | 4.068764 | 2.874718 | 3.857328 |
| 0.20228 | 0.058704 | 0.561741 | 1.205669 | 1.068287 | 0.957706 | 1.235126 | 1.180984 | 1.447004 |
| -0.37187 | 1.160564 | -0.11058 | -0.11476 | 0.857944 | 0.553724 | 1.63015 | 2.749489 | -0.80583 |
| 0 | 0 | 0 | 0 | 0 | 0 | 0 | 0 | 0 |
| 0.073397 | 0.126076 | -0.19437 | 0.605982 | 0.222996 | 0.220549 | -0.21231 | 1.798815 | -0.07649 |
| -0.4289 | 4.723953 | 2.163992 | 4.305941 | 2.829391 | 2.920273 | 4.895182 | 2.535851 | 3.070181 |
| -2.5795 | 4.685426 | 1.546764 | 3.682835 | 2.541822 | 2.983522 | 3.244979 | 2.655644 | 2.512338 |
| -3.22433 | 4.552865 | 1.867785 | 3.632946 | 2.410328 | 2.764263 | 2.418006 | 2.185529 | 2.65167 |
| -0.59593 | 1.012351 | 1.434368 | 1.136503 | 1.38582 | 1.535299 | 0.717496 | 2.450997 | 1.837174 |
| 0.504402 | 0.737296 | 0.825563 | 0.990917 | 0.958347 | 0.530523 | 0.67463 | 0.994134 | 0.848147 |
| -1.12155 | 5.93101 | 2.558447 | 3.855085 | 2.90078 | 3.437714 | 1.843331 | 1.857865 | 3.234567 |
| -1.56607 | 4.671979 | 1.752056 | 2.957809 | 2.191259 | 2.620207 | 1.997191 | 1.258086 | 2.472018 |
| -1.95569 | 6.258279 | 5.445702 | 7.325218 | 5.376686 | 5.721621 | 3.173686 | 4.815396 | 6.558381 |
| -2.53528 | 4.990552 | 4.053436 | 7.869223 | 4.139456 | 3.745918 | 3.729926 | 4.769273 | 7.433282 |
| -1.71306 | 5.515738 | 2.373362 | 3.565256 | 3.267864 | 3.161163 | 2.016181 | 2.898126 | 3.19711 |
| -1.01324 | 4.923808 | 2.254465 | 2.383436 | 2.338133 | 2.930025 | 2.037532 | 1.039895 | 1.734712 |
| -0.40333 | 4.5677 | 3.371281 | 4.247047 | 2.584507 | 3.396654 | 2.741635 | 2.425221 | 4.758327 |
| 0.862383 | 2.339847 | 3.061172 | 2.264214 | 2.927406 | 2.675659 | 0.705777 | 2.248446 | 0.751576 |
| -1.10408 | 0.159484 | 1.231421 | 1.393646 | 1.281086 | 1.369051 | -0.90296 | 0.242871 | 0.634728 |
| -1.38346 | 4.899325 | 1.468987 | 3.74535 | 1.60515 | 2.171541 | 3.271285 | 2.714145 | 3.614207 |
| -3.70331 | 4.568688 | 1.890629 | 3.414524 | 2.521643 | 2.875835 | 3.117669 | 0.994407 | 2.534532 |
| -2.35034 | 4.688132 | 1.125086 | 2.875856 | 2.208115 | 2.190144 | 2.257794 | 0.681565 | 1.794206 |
| -0.5044 | -0.7373 | -0.82556 | -0.99092 | -0.95835 | -0.53052 | -0.67463 | -0.99413 | -0.84815 |

| Lung_IL18 | Lung_IRF1 | Lung_IRF7 | Lung_IRF8 | Lung_CXCL | Lung_CXCL | Lung_IFIH1 | Lung_DDX5 | Lung_KLF2 |
|---|---|---|---|---|---|---|---|---|
| 0.322763 | 0.091566 | -0.13216 | -0.13342 | -1.65371 | 1.809511 | 0.26893 | -0.15213 | -0.41366 |
| 0.317965 | 0.063927 | 0.513094 | 0.281738 | -0.61478 | 1.032038 | 0.467869 | 0.21245 | -0.71003 |
| 0.617321 | -0.00742 | -0.19653 | -0.68502 | 0.055487 | 0.113106 | 0.075428 | -0.02247 | 0.130033 |
| 0.047571 | 0.533134 | 0.324574 | 0.231766 | -1.82571 | 0.296251 | 0.520851 | 0.459885 | -0.02106 |
| 0.209225 | 0.27931 | 0.608604 | -0.33717 | 0.125412 | -0.14164 | 0.434898 | 0.40444 | -0.24352 |
| -0.64494 | 0.874138 | 3.586357 | 0.504564 | 4.912693 | 9.213177 | 1.37216 | 0.698244 | -0.39483 |
| 0 | 0 | 0 | 0 | 0 | 0 | 0 | 0 | 0 |
| -0.80796 | 0.833914 | 3.231615 | -0.30738 | 5.272484 | 8.23378 | 1.156273 | 0.954697 | -0.66678 |
| 0.180542 | -0.21646 | -0.56178 | -0.36162 | -1.39461 | -1.0628 | -0.0125 | -0.34069 | -0.6722 |
| -1.20571 | 1.917509 | 4.114174 | 0.955481 | 5.345842 | 10.72459 | 1.871796 | 1.218136 | -0.3526 |
| -0.29411 | 0.704407 | 3.427227 | -0.4835 | 4.658302 | 8.699492 | 1.334953 | 0.857353 | -0.97876 |
| -0.14849 | 0.090096 | -0.42433 | 0.283827 | -1.235 | -0.59643 | 0.503395 | 0.861765 | 0.079369 |
| -0.48893 | 0.076918 | -0.20847 | 0.120121 | -0.29892 | -0.30324 | 0.562923 | 0.836254 | -0.44392 |
| -0.85732 | 2.723038 | 3.990625 | 1.798109 | 9.017725 | 10.85167 | 2.337395 | 2.109247 | 1.340347 |
| -0.33286 | 0.581955 | 3.968485 | | 7.382212 | 7.285721 | 1.399206 | 1.314837 | -1.28638 |
| -1.15537 | 1.824093 | 4.391897 | 1.659199 | 11.25144 | 10.85554 | 2.090113 | 1.734888 | 0.757179 |
| -0.07276 | 0.029924 | 0.557196 | 0.126518 | 0.475266 | -0.74554 | 0.225557 | 0.371172 | -0.90206 |
| -0.4953 | 1.375633 | 4.046986 | 0.883257 | 8.012733 | 10.72135 | 1.609838 | 1.219316 | 1.936146 |
| 0.661978 | -0.01838 | 1.453131 | 0.22127 | 2.494762 | 0.443451 | 0.517029 | 0.283974 | 1.502903 |
| -0.861 | 0.106644 | -0.19848 | -0.50809 | 3.351601 | -1.05272 | 0.189196 | 0.113379 | 0.74283 |
| 0 | 0 | 0 | 0 | 0 | 0 | 0 | 0 | 0 |
| 0.458378 | -0.31367 | 0.076651 | -0.18803 | 1.275023 | -0.51393 | -0.1172 | -0.16991 | 0.685184 |
| -0.73705 | 1.057592 | 3.576998 | 0.532995 | 7.173359 | 9.711451 | 1.930325 | 1.957624 | 2.313793 |
| -0.01406 | 0.623747 | 4.25771 | 0.219824 | 2.274023 | 5.732772 | 1.268022 | 1.112663 | 2.416012 |
| -0.78798 | 0.422405 | 3.966089 | 0.286589 | 3.074734 | 5.176711 | 0.981742 | 0.711203 | -0.54845 |
| 0.749286 | -0.6414 | 0.128128 | -0.59156 | -0.60296 | 0.033772 | -0.27476 | -0.43695 | 0.205532 |
| 0.672072 | -0.23629 | 0.093878 | -0.29718 | 0.837635 | 0.434434 | -0.40918 | -0.30779 | -0.11786 |
| 0.100644 | 0.119225 | 3.543144 | -0.21478 | 1.346941 | 3.660798 | 0.794337 | 0.489363 | 1.257046 |
| -0.70898 | 0.017056 | 3.57662 | -0.07338 | 2.479713 | 5.016572 | 0.732892 | 0.150713 | -0.39227 |
| 0.746274 | 1.570963 | 3.386606 | 1.562729 | 4.858551 | 6.482692 | 2.284846 | 1.554861 | -4.23071 |
| -0.13837 | 2.969131 | 2.794878 | 2.79545 | 6.270304 | 8.006883 | 3.03807 | 2.77446 | 2.961834 |
| 0.099704 | -0.2676 | 2.877684 | 0.255318 | 4.007782 | 4.80618 | 0.806878 | 0.346273 | 2.129389 |
| 0.174607 | 0.041809 | 3.639488 | 0.071255 | 1.228159 | 3.66735 | 0.592637 | 0.345196 | -0.66261 |
| 0.156253 | 1.371668 | 3.878004 | 1.088137 | 4.345003 | 6.568391 | 1.485425 | 1.296049 | 3.153347 |
| | 0.235857 | 0.670759 | -0.02005 | -1.43229 | 0.267838 | 0.24071 | 0.226742 | 2.796049 |
| 0.349273 | -0.52015 | -0.04786 | -0.53661 | -0.60528 | 0.455205 | -0.3042 | -0.67141 | 1.776672 |
| -1.44006 | 1.470282 | 3.795326 | 2.209362 | 5.463575 | 6.876754 | 1.225192 | 0.846418 | 2.61795 |
| -0.14618 | 0.268057 | 3.998457 | 0.069086 | 1.998632 | 3.834901 | 0.592074 | 0.706936 | 2.235977 |
| -1.22611 | 0.310179 | 3.599628 | -0.1628 | 1.960403 | 3.941318 | 0.708371 | 0.661503 | 1.272533 |
| -0.67207 | 0.23629 | -0.09388 | 0.297176 | -0.83764 | -0.43443 | 0.409183 | 0.307791 | 0.117859 |

| Liver_ICAM | Liver_SELE | Liver_IL6 | Liver_SOCS | Liver_STAT | Liver_CEM | Liver_F3 | Liver_RHO | Liver_TSP1 |
|---|---|---|---|---|---|---|---|---|
| 0.134438 | 0.192612 | -1.53181 | -0.03926 | -1.00434 | -2.35918 | 2.305107 | 1.435249 | 0.045464 |
| -0.58114 | -0.04835 | 0.036179 | 0.763916 | 0.333153 | 0.808102 | 0.760393 | 0.225193 | 1.00596 |
| 0.022757 | 0.452932 | -0.55338 | 0.680624 | 1.436432 | -2.10987 | 0.430883 | 1.68655 | -0.29113 |
| 0.447216 | -1.18509 | 0.704247 | -0.1163 | 0.232601 | -2.38444 | 1.719448 | 1.242739 | 0.660194 |
| -0.23057 | -0.4091 | -0.53659 | -1.76798 | -0.27834 | 0.660267 | -0.38177 | 0.978613 | 0.766232 |
| 2.85297 | 0.937992 | 3.774815 | 4.927717 | 1.062023 | 5.705448 | -1.2565 | -0.18683 | 0.853441 |
| 0 | 0 | 0 | 0 | 0 | 0 | 0 | 0 | 0 |
| 0.629782 | -0.05797 | 0.043112 | -0.05425 | 0.398714 | 0.940287 | -0.54966 | 0.561653 | -0.43678 |
| 2.239267 | -0.09941 | 2.382437 | 3.601738 | 0.425369 | 4.835032 | -1.41057 | -0.6404 | -0.18639 |
| 2.993464 | 0.462931 | 3.840715 | 4.85465 | 1.050423 | 6.01259 | -0.8243 | -0.5229 | 0.268681 |
| 1.867887 | 0.41749 | 1.251749 | 3.269682 | -0.44289 | 2.197523 | -2.60499 | -1.39635 | -1.48703 |
| 0.437616 | -1.04453 | 1.302492 | 3.141729 | 0.930691 | 1.081533 | -0.91315 | 0.556438 | -1.91973 |
| 0.972414 | -0.80232 | 1.494547 | 3.22089 | 1.103371 | 2.268457 | -0.53333 | 1.520699 | 0.349491 |
| 3.928272 | 2.680834 | 6.199553 | 6.429426 | 1.487377 | 5.852024 | -0.62277 | -0.38333 | 1.433561 |
| 2.978481 | 1.213512 | 2.487476 | 4.946543 | 0.750921 | 1.159672 | -1.04435 | 0.415701 | -0.35608 |
| 3.916895 | 2.35931 | 7.003735 | 6.466637 | 1.873302 | 6.095951 | 0.113077 | 0.405239 | 3.010353 |
| 0.018827 | 0.129225 | -0.13206 | 1.488516 | 0.07802 | 0.587084 | 0.182892 | 0.193346 | -1.64026 |
| 3.407925 | 0.230425 | 6.700598 | 5.954681 | 0.767387 | 5.562553 | -1.18797 | 0.034468 | 0.473499 |
| 1.170818 | -1.78465 | 0.32552 | 0.51012 | 0.184792 | | 0.232107 | 0.338764 | -1.17487 |
| -1.6118 | -0.67191 | 0.276915 | 4.253107 | 0.261902 | 5.268505 | -1.3781 | -0.36317 | 0.035486 |
| 0 | 0 | 0 | 0 | 0 | 0 | 0 | 0 | 0 |
| 0.013939 | -2.5404 | 0.190319 | 1.961992 | 0.142563 | -0.15932 | -0.06266 | 0.104622 | -1.88965 |
| 2.277514 | -0.02656 | 4.129902 | 4.983885 | -0.23649 | 2.698072 | -2.91407 | -1.10175 | -1.4474 |
| 1.909697 | 0.769535 | 2.25588 | 2.788642 | -0.16633 | -1.99817 | -1.35896 | -1.14001 | 0.873483 |
| 1.082699 | 1.629038 | 1.389565 | 3.302098 | 0.463883 | 0.978291 | -2.13811 | -1.68262 | 1.207543 |
| -0.06426 | 0.970028 | 0.390978 | 0.966641 | 0.294555 | 0.881247 | -0.91856 | 0.929865 | 1.492429 |
| 0.4688 | -1.60938 | -0.71508 | -0.105 | 0.199508 | 0 | -0.37132 | -0.32291 | -0.36025 |
| 2.578995 | 1.023035 | 1.626858 | 1.424767 | -0.9806 | -3.13028 | -1.59442 | -1.79834 | -0.14371 |
| 2.528852 | 2.140991 | 3.5142 | 3.253379 | 0.055617 | -0.58946 | -0.48694 | -0.45655 | 0.804345 |
| 4.491749 | 3.958673 | 4.874016 | 3.812186 | -0.28203 | 2.933744 | -1.17086 | -1.94436 | 1.661738 |
| 4.873938 | 5.645809 | 7.026979 | 6.352304 | 1.760079 | 4.984541 | 0.554679 | 1.562222 | 4.03756 |
| 2.160143 | 1.510172 | 2.361444 | 2.817124 | -0.70692 | 2.061031 | -1.32832 | -1.55456 | 1.239253 |
| 0.4582 | 1.297699 | 1.286745 | 2.310676 | -1.65597 | -1.5648 | -1.90402 | -1.95471 | -1.55727 |
| 3.347286 | 1.846107 | 3.832672 | 3.705352 | -0.81701 | 2.067366 | -0.26049 | -1.73685 | 1.385262 |
| 0.305685 | 1.323917 | 1.776052 | 1.230468 | 0.145551 | 0.246775 | 0.321675 | 0.236273 | 0.767627 |
| 0.306412 | -0.6555 | 0.73592 | 1.392337 | -0.07084 | -1.85938 | 1.161414 | -0.85873 | -0.33203 |
| 4.419186 | 2.923441 | 6.092335 | 5.723077 | 0.844367 | 4.657385 | -0.32996 | 1.041618 | 2.252389 |
| 2.499973 | 1.145319 | 2.992512 | 3.999314 | 0.262382 | 0.776707 | -0.85224 | -0.72132 | 0.946374 |
| 0.692875 | 1.793409 | 1.539165 | 3.606835 | 0.084952 | -2.20616 | -1.28686 | -0.78449 | 0.739831 |
| -0.4688 | 1.609385 | 0.715084 | 0.104999 | -0.19951 | | 0.371324 | 0.322906 | 0.360246 |

| Liver_PLAT | Liver_PLAU | Liver_THBD | Liver_PLAU | Liver_ET1 | Liver_PTGS | Liver_VWF | Liver_PRO( | Liver_CD44 |
|---|---|---|---|---|---|---|---|---|
| 0.600952 | 0.633568 | -0.12667 | -1.00417 | 0.107891 | -1.3258 | 1.419785 | -0.496 | -2.89417 |
| 0.298569 | 0.601242 | -0.56931 | 2.069584 | 1.632851 | 1.106098 | 2.340246 | 0.355541 | -0.31989 |
| -1.53143 | -2.32231 | 0.711897 | -0.31993 | 1.105143 | -1.0765 | 1.068254 | -0.98574 | -2.38693 |
| -1.90229 | -1.24501 | -0.5092 | -1.30728 | 0.337505 | 1.523796 | 0.212078 | 1.671022 | -1.92597 |
| -0.41386 | 0.141804 | 1.341091 | 0.01083 | 0.862736 | 1.111481 | 1.640676 | -0.96684 | -2.16986 |
| 1.386497 | -0.9319 | 2.704594 | 3.742693 | 2.79525 | 3.577797 | -1.44049 | 3.175478 | 1.371834 |
| 0 | 0 | 0 | 0 | 0 | 0 | 0 | 0 | 0 |
| 0.967131 | 0.367376 | -0.33932 | -0.26277 | 1.612434 | 1.999506 | 1.176706 | 1.454575 | -0.20961 |
| 0.399065 | -0.43217 | 2.449608 | 2.338251 | 1.281984 | 3.07733 | -1.08901 | 3.434866 | -1.74804 |
| 1.221567 | -1.3624 | 1.227701 | 3.074333 | 2.496693 | 6.641222 | -0.64499 | 3.389751 | 0.633095 |
| -1.65137 | -3.0874 | 1.147249 | 1.136635 | 0.28796 | 1.68371 | -1.37528 | 3.808889 | 0.759605 |
| 1.101118 | -1.46323 | -0.17797 | -1.20107 | 1.287678 | | 1.293224 | -0.63157 | |
| 0.669941 | -0.28563 | 1.049576 | 0.12813 | 1.501245 | | 0.665695 | -0.54632 | |
| -0.49422 | -0.7767 | 2.23794 | 4.770363 | 3.292469 | | -0.28324 | -0.07425 | |
| 2.435057 | 0.324707 | 2.311651 | 2.233738 | 1.578081 | | -0.30788 | 3.159504 | |
| 0.805717 | -1.29817 | 1.679293 | 5.409969 | 5.768402 | | 1.112383 | 4.194666 | |
| 0.304462 | -1.06288 | 0.337868 | -1.80176 | 0.372641 | | -0.65922 | 0.786434 | |
| 0.235992 | -0.93458 | 1.231443 | 3.224895 | 1.993525 | | 0.074518 | 4.820974 | |
| 0.146914 | 1.125637 | 0.559374 | -0.14348 | 1.058382 | | 1.192116 | 1.040665 | |
| -0.56327 | -1.20153 | 1.462076 | 1.090467 | -0.24836 | | 0.948254 | 1.624664 | |
| 0 | 0 | 0 | 0 | 0 | | 0 | 0 | |
| 0.519361 | -0.88229 | -0.21919 | -1.58746 | 0.146835 | | 0.292976 | 0.272509 | |
| -0.03434 | -2.90153 | 1.777735 | 1.690693 | 0.43206 | | -2.2236 | 2.247162 | |
| 0.442167 | -1.98882 | 2.089911 | 0.334362 | 0.670271 | 0.500095 | -2.42385 | 3.380147 | -1.74046 |
| 0.43618 | -0.69048 | 1.415562 | 0.784479 | -0.52066 | 1.967066 | -2.44417 | 1.857547 | |
| -2.68043 | 0.536463 | 0.462574 | -1.74683 | -0.97541 | -0.23101 | 0.200583 | 0.429561 | |
| -4.64118 | -1.99241 | 0.644686 | -2.81395 | -0.29762 | -0.25319 | -0.10852 | 0.202548 | 0.096145 |
| -0.67128 | -2.53201 | 0.898802 | 0.070528 | 0.779289 | -0.26907 | -2.8743 | 2.643537 | -3.40917 |
| 3.701214 | -1.59509 | 2.456373 | 1.414236 | 1.725229 | 2.191856 | -0.79595 | 4.207513 | -1.51178 |
| -0.71317 | -0.97581 | 2.226681 | 1.860922 | 2.300667 | 4.929186 | -1.70544 | 4.474847 | -0.42305 |
| 3.207153 | 0.030115 | 2.415071 | 4.809357 | 4.300032 | 7.673098 | 1.519559 | 3.784446 | 2.192163 |
| 0.657698 | -2.30319 | 0.789532 | 1.137157 | 0.510891 | 1.510393 | -2.4264 | 2.253631 | -1.52577 |
| 2.480835 | -3.17033 | 1.60754 | -0.78212 | -0.05242 | -1.21291 | -3.54283 | 1.903354 | -3.33631 |
| 3.664017 | -1.77638 | 1.331993 | 1.927662 | 1.76383 | 1.754753 | -0.47045 | 4.819821 | -1.55527 |
| 3.667269 | -0.38236 | 1.454617 | -1.38397 | -0.34768 | -2.92538 | 0.42397 | 1.016345 | -0.53443 |
| 0.675585 | -1.79737 | -0.31541 | -1.93572 | -0.74416 | 0.923826 | -0.29876 | -0.00876 | |
| 3.607878 | -0.90459 | 2.688269 | 2.310381 | 3.357351 | 5.694397 | -1.08036 | 4.436751 | -0.16339 |
| 2.920835 | 0.279652 | 1.576521 | -0.11691 | 0.828171 | 0.87295 | -0.76062 | 3.091054 | -4.11112 |
| 3.280098 | -0.94452 | 2.543079 | 0.314302 | 0.456161 | 0.259859 | -1.36373 | 2.74347 | |
| 4.641176 | 1.992409 | -0.64469 | -1.38647 | 0.297623 | 0.253187 | 0.108518 | -0.20255 | -0.09614 |

| Liver_TFPIA | Liver_TFPIB | Liver_TFPIC | Liver_SELP | Liver_C5AR | Liver_VEGF | Liver_SERP | Liver_STAT | Liver_OASL |
|---|---|---|---|---|---|---|---|---|
| 0.611694 | 0.754799 | 0.637129 | 0.954575 | -0.69623 | 0.721827 | -0.63098 | 2.472965 | 0.675558 |
| 1.248743 | 1.276073 | 0.705894 | 4.514338 | 2.170406 | -0.02917 | 0.539871 | 1.665136 | 1.954882 |
| 0.560665 | 1.049503 | -0.27603 | 2.233942 | 0.074013 | 0.603123 | -1.10657 | 2.552603 | 0.57795 |
| 2.424864 | 2.566338 | 2.086939 | 4.000574 | -0.77014 | -0.72399 | 0.274385 | 2.130299 | -0.68539 |
| 1.297453 | 0.933544 | 0.124998 | 3.572058 | -1.07566 | -0.04918 | -0.68032 | 1.248423 | 1.506887 |
| -0.00723 | -1.48107 | 0.592825 | 6.631907 | -0.18237 | -0.02999 | 8.560869 | 3.858919 | 0.865887 |
| 0 | 0 | 0 | 0 | 0 | 0 | 0 | 0 | 0 |
| 0.503 | 1.581493 | 0.242441 | 1.95363 | 1.708858 | -0.8313 | -1.11501 | 0.973803 | -0.75323 |
| 0.825323 | -0.97623 | 0.546139 | 5.094646 | 2.33662 | -0.64458 | 5.106773 | 1.510977 | -0.02323 |
| -0.2746 | -3.06705 | -0.68777 | 5.509991 | 1.760544 | 0.108194 | 6.552444 | 0.550056 | 1.780815 |
| 0.923044 | -0.21567 | 0.841986 | 4.158083 | 0.802757 | -1.59126 | 7.171038 | -0.55769 | -0.83426 |
| -0.05074 | 0.215391 | 0.294884 | 3.36586 | 1.987726 | 0.85083 | 0.003914 | 0.891939 | 0.620514 |
| 0.144218 | 0.53273 | 0.428709 | 4.846956 | 1.821552 | 1.91284 | 0.343115 | -1.08555 | -0.86326 |
| -0.15539 | -2.1222 | 0.669291 | 7.603731 | 4.553307 | 1.643305 | 9.058363 | 2.464266 | 3.472616 |
| -0.0265 | 0.78301 | 1.389853 | 7.342264 | 4.098963 | 2.18693 | 3.716837 | 2.465879 | 1.467045 |
| 0.675835 | -1.45305 | 1.160282 | 9.12921 | 5.280737 | 2.423498 | 8.602314 | -0.29382 | 3.455431 |
| -0.51528 | -0.95635 | -0.15774 | 1.345095 | 0.598221 | 0.683889 | -1.82019 | -0.31513 | -0.49298 |
| -0.12326 | -1.3142 | 0.920788 | 4.607075 | 2.663584 | 2.94463 | 7.646662 | 1.923384 | 1.570236 |
| 1.705194 | 0.295151 | 0.906168 | 1.476109 | 1.337181 | 1.041546 | 0.393705 | 0.660191 | 1.182291 |
| 0.962639 | 0.63187 | 0.769381 | 6.717472 | 1.218348 | 2.801851 | 5.981428 | 1.392641 | -1.85342 |
| 0 | 0 | 0 | 0 | 0 | 0 | 0 | 0 | 0 |
| 0.048599 | -0.05196 | -0.34395 | 0.785397 | 1.272167 | 2.050352 | -0.41254 | 0.114439 | 0.73488 |
| -1.53289 | -3.10555 | -0.97256 | 6.550749 | 3.716082 | 0.750803 | 7.229824 | 0.557978 | -0.77261 |
| -0.08363 | -0.84114 | 0.380824 | 4.165033 | 2.741943 | -1.80658 | 5.202973 | -0.09021 | -0.49175 |
| -0.57539 | -2.81593 | -0.26935 | 5.336552 | 3.373051 | -0.19267 | 6.93678 | -0.43377 | -1.5122 |
| -1.07965 | -0.04783 | -0.28773 | -0.28484 | 0.069815 | 0.225626 | -0.12383 | -0.23991 | 0.555232 |
| -0.13448 | -0.41194 | -0.80418 | -1.65012 | -0.2707 | -0.62355 | -0.26962 | -0.00699 | -0.39415 |
| -0.68535 | -1.51392 | -0.11477 | 2.041159 | 1.418716 | -0.62462 | 6.352029 | -0.32967 | -0.82481 |
| 0.000823 | -0.46819 | 0.504619 | 3.12553 | 4.105886 | -0.97347 | 7.062757 | -0.25216 | 0.043351 |
| -0.38754 | -3.06754 | -0.16924 | 2.230228 | 5.088167 | -2.7119 | 8.48767 | 0.192434 | 1.640121 |
| -0.12952 | -1.58958 | -0.48182 | 7.018087 | 5.665831 | 2.967979 | 10.22461 | -0.08269 | 3.976336 |
| -1.15953 | -3.05753 | -1.28814 | 0.433176 | 2.695938 | -2.09249 | 6.664779 | -0.94958 | -1.38082 |
| -0.44466 | -2.4424 | -0.3662 | 3.066576 | 3.092417 | -1.12948 | 4.403232 | -0.5286 | -1.55947 |
| -0.38379 | -1.84776 | -0.08579 | 4.721973 | 4.871653 | -1.55375 | 7.294583 | -0.05372 | 0.734393 |
| -0.10475 | 0.551761 | 0.389134 | 0.49971 | 4.165392 | -1.06024 | -0.42652 | 0.048973 | -0.09304 |
| -0.21279 | 0.699521 | 0.235269 | 1.22068 | 2.415421 | -1.7237 | -0.07362 | 0.480299 | -0.17039 |
| 0.350476 | -2.2771 | 0.746257 | 4.969149 | 4.579374 | 1.020784 | 10.48785 | 0.101147 | 0.82277 |
| 0.716058 | -0.35904 | 0.422439 | 3.477304 | 1.834843 | 0.929445 | 6.350297 | 0.836262 | 0.229877 |
| 0.180354 | -0.45637 | 0.386575 | 5.411156 | 4.294973 | 0.278849 | 7.649146 | 0.325243 | -0.57246 |
| 0.134477 | 0.411937 | 0.804175 | 1.650122 | 0.270699 | 0.623549 | 0.269622 | 0.006992 | 0.394151 |

| Liver_OAS2 | Liver_OASL | Liver_OAS3 | Liver_FOS | Liver_MX1 | Liver_IL18 | Liver_IRF1 | Liver_IRF7 | Liver_IRF8 |
|---|---|---|---|---|---|---|---|---|
| 0.263638 | -0.05541 | 2.189655 | -3.76981 | -0.38907 | 0.618977 | 0.428713 | 0.20355 | 0.087296 |
| 0.55526 | 1.061731 | 2.250204 | -2.13139 | 1.556696 | 0.026621 | 0.504925 | 0.440355 | 0.657644 |
| 0.406752 | -0.02647 | 0.933704 | -2.68115 | 0.351242 | 0.227467 | 0.201773 | 0.047907 | 0.512527 |
| -0.38584 | -0.88198 | 0.84864 | -3.23645 | -0.83873 | 0.122622 | 0.236923 | 0.155525 | 0.242025 |
| 0.461632 | 1.156136 | 2.609543 | -5.6049 | 0.326237 | 0.752586 | -0.10711 | 0.801764 | 1.001278 |
| 0.654112 | 0.212114 | 1.228096 | -7.09229 | 1.002728 | -2.29422 | -0.54348 | 0.717768 | 1.029026 |
| 0 | 0 | 0 | 0 | 0 | 0 | 0 | 0 | 0 |
| -1.09322 | 0.995171 | -0.34607 | -5.63383 | -1.16491 | 0.167553 | -0.03281 | -0.22746 | -0.05546 |
| 0.399714 | -0.03307 | 1.751421 | -6.94519 | -0.06686 | -2.49978 | -1.38012 | -0.44402 | -0.03449 |
| -0.67801 | -0.86408 | 1.755795 | -7.42485 | 2.388548 | -2.51131 | -0.14664 | 0.449669 | 1.392389 |
| -2.95863 | 0.03278 | 2.815296 | -8.82858 | 0.336687 | -3.23633 | -1.37411 | -0.57808 | -0.67436 |
| 0.383776 | -0.08679 | -1.85488 | 1.913801 | 2.649532 | 1.203314 | 1.437572 | 1.694626 | 1.152765 |
| 1.254869 | 0.471539 | 1.279982 | 3.70846 | 2.633097 | 1.554155 | 1.229675 | 1.282055 | 1.814585 |
| 2.684561 | 3.132298 | 1.615671 | 2.895691 | 6.196445 | -0.84164 | 2.595547 | 1.756008 | 3.797964 |
| 3.441557 | 2.982155 | 2.383125 | 2.972982 | 3.043047 | 0.059507 | 1.478048 | 3.235502 | 0.478674 |
| 3.911779 | 3.228922 | 3.517769 | 4.157698 | 7.487839 | -0.03062 | 1.298847 | 2.959263 | 5.418222 |
| -1.85107 | -1.6046 | -1.08947 | -0.68664 | 1.607481 | -0.83149 | -0.13695 | 0.516047 | 2.460382 |
| 1.629677 | 2.753458 | 2.84947 | 2.377207 | 3.272242 | -1.92571 | 1.076981 | 1.37785 | 3.300781 |
| -1.58373 | 1.439886 | -0.00628 | 0.056389 | 1.243088 | 0.468941 | 0.03779 | 1.674004 | 2.036499 |
| 1.177998 | -0.90393 | -0.44821 | 5.052406 | -0.77249 | -1.12013 | -0.38505 | -1.6755 | 0.89826 |
| 0 | 0 | 0 | 0 | 0 | 0 | 0 | 0 | 0 |
| -0.50718 | -0.3661 | -1.34338 | 0.982243 | 1.672279 | 0.174484 | 0.477631 | 1.162279 | 1.139967 |
| -1.12042 | 1.200552 | 1.056467 | 1.033823 | 0.818048 | -2.54311 | 0.316986 | 0.584867 | 1.177933 |
| 0.175931 | 1.392542 | 2.365132 | 1.643565 | 0.229054 | -1.83582 | -0.87153 | 0.638165 | 0.355429 |
| -0.65134 | 0.621187 | 1.27476 | 0.40805 | -0.1777 | -3.49022 | -1.23038 | 0.27061 | 0.257524 |
| -1.05671 | -0.23248 | -0.50122 | 1.955228 | 0.393567 | -0.63619 | -1.12189 | 0.227269 | -0.03161 |
| -0.132 | -0.2062 | -0.50521 | -0.53134 | 0.055879 | -0.24563 | -0.0521 | -0.12818 | -0.73134 |
| -0.17231 | 0.980799 | 1.385676 | -0.21216 | 0.090009 | -2.33828 | -1.50319 | 0.25426 | 0.007425 |
| 0.8474 | 1.633303 | 2.527806 | 1.519811 | 0.395127 | -1.40647 | -0.68341 | 0.57368 | 0.009327 |
| 1.336131 | 2.438147 | 3.590957 | 1.357493 | 3.275487 | -2.35629 | -0.1309 | 0.789775 | 1.412634 |
| 1.476232 | 1.732159 | 3.052238 | 2.928842 | 6.174495 | -1.82975 | 0.760052 | 0.265626 | 2.325821 |
| -0.85074 | 0.254047 | 0.88403 | -0.3265 | -0.79827 | -2.68772 | -1.90343 | -0.71905 | 0.095993 |
| -1.13937 | 0.029556 | 0.214923 | -0.71539 | -0.36822 | -2.51198 | -1.58844 | -0.07696 | -0.68338 |
| 0.881007 | 1.875128 | 3.142478 | 1.060994 | 2.862903 | -1.78495 | -0.18832 | 1.000504 | 1.859867 |
| 0.795737 | 0.576372 | 1.623467 | 1.060023 | 0.53095 | 0.663541 | 0.401077 | 0.92266 | 1.187653 |
| -0.30894 | 0.458679 | -0.82501 | -0.07677 | 0.605592 | 0.012803 | 0.35511 | 0.44106 | -0.29351 |
| 1.075198 | 1.794348 | 2.641883 | 2.727439 | 2.37485 | -1.71926 | 0.225615 | 0.801641 | 2.414667 |
| 0.759057 | 1.767649 | 2.485824 | 0.621421 | 1.27192 | -1.73858 | -0.00817 | 1.225448 | 0.99246 |
| 0.122937 | 1.009657 | 1.889344 | 0.310513 | 0.633729 | -2.00494 | -0.61207 | 0.631692 | 0.034807 |
| 0.132001 | 0.206196 | 0.505214 | 0.531341 | -0.05588 | 0.245633 | 0.052095 | 0.128184 | 0.731344 |

| Liver_CXCL | Liver_CXCL | Liver_IFIH1 | Liver_DDX5 | Liver_KLF2 | Liver_HNF1 |
|---|---|---|---|---|---|
| -1.36317 | 0.95731 | 0.493565 | 0.308855 | -0.13679 | 0.080507 |
| 0.041475 | 1.577154 | 0.533447 | 0.490557 | -0.18397 | 0.33322 |
| -0.03035 | 0.667221 | 0.433825 | 0.903616 | 1.777164 | 0.365881 |
| -0.54302 | 0.633476 | 0.283134 | 0.101345 | -0.43969 | -0.34381 |
| -0.46115 | 1.450491 | 0.866093 | 1.004646 | 1.122276 | 0.592606 |
| 3.738552 | 7.065247 | -0.90606 | -0.88185 | 0.873407 | -0.13396 |
| 0 | 0 | 0 | 0 | 0 | 0 |
| 0.401836 | -0.26968 | -0.11662 | 0.183739 | -0.04109 | -0.2346 |
| 2.021235 | 4.49819 | -1.76628 | -1.39098 | -0.41855 | -0.68668 |
| 4.091976 | 6.609978 | -0.88219 | -1.3539 | 3.355675 | -0.65572 |
| 1.934416 | 5.943825 | -1.72341 | -1.56607 | 1.573938 | -1.50224 |
| | -1.2644 | -0.04898 | -0.21921 | 1.710983 | 1.357626 |
| 3.159487 | 0.033834 | -0.00281 | 0.361795 | 1.526342 | 0.733961 |
| 8.449066 | 8.933098 | 0.514673 | 0.18961 | 2.018227 | 0.872683 |
| 4.861778 | 6.590187 | -0.39037 | 0.553297 | 1.25835 | 1.07439 |
| 11.13791 | 8.969803 | 0.700928 | -0.13718 | 3.248339 | 1.116924 |
| 2.374268 | -0.83018 | -0.21305 | -0.69618 | -1.63671 | -0.69147 |
| 7.964956 | 6.785204 | -0.86966 | -0.7119 | 1.812347 | 0.801331 |
| 3.246798 | 5.739449 | 0.220364 | 0.396957 | 0.447346 | 0.340843 |
| 3.179937 | 0.697405 | -1.23088 | -0.689 | 0.658049 | 0.253586 |
| 0 | 0 | 0 | 0 | 0 | 0 |
| 0.312546 | -0.74778 | 0.029316 | 0.099384 | -0.00834 | 1.070303 |
| 6.362776 | 5.742056 | -2.02203 | -2.01428 | 0.068415 | 0.443659 |
| 3.360962 | 7.028597 | -0.75185 | -0.17036 | -0.00402 | -0.61358 |
| 3.601076 | 7.075659 | -1.5734 | -0.71452 | 0.037589 | -0.38415 |
| 1.80909 | -0.59017 | -0.27417 | 0.715085 | 0.233699 | 0.369961 |
| 0.179132 | -1.03304 | -0.09097 | -0.77987 | -1.18731 | -0.66865 |
| 3.093073 | 5.337986 | -1.19285 | -1.38284 | 0.703225 | -1.8714 |
| 4.215965 | 5.568108 | -0.99617 | -0.57017 | 0.794822 | -0.48407 |
| 5.256285 | 6.661079 | -0.09453 | -0.92314 | 1.202609 | -1.24349 |
| 8.724308 | 9.062132 | 0.6536 | 1.219142 | 3.006494 | -2.15382 |
| 3.624567 | 5.884689 | -1.65023 | -1.54404 | 0.247867 | -0.81222 |
| 1.9643 | 5.658554 | -0.93416 | -1.3366 | -0.33811 | -1.3996 |
| 4.794151 | 5.966402 | -0.70274 | -0.74988 | 0.805102 | -0.58692 |
| 1.223915 | 1.16453 | 0.351409 | 0.802518 | 0.200927 | 0.367935 |
| 1.401865 | 0.1637 | 0.446347 | -0.48024 | -1.54806 | -0.23161 |
| 7.000698 | 7.244959 | -0.25274 | -0.35166 | 1.792283 | 0.621481 |
| 4.316435 | 6.444992 | -0.14921 | 0.614669 | -0.03131 | -0.03227 |
| 3.770933 | 6.217987 | -0.76416 | -0.35832 | 0.064776 | 0.040079 |
| -0.17913 | 1.033035 | 0.09097 | 0.779868 | 1.18731 | 0.66865 |
